# Human - murine concordance of molecular signatures in nerve-sparing murine partial bladder outlet obstruction (NeMO)

**DOI:** 10.1101/2021.09.15.460523

**Authors:** Martin Sidler, Abdalla Ahmed, Jia-Xin Jiang, Dursa Koshkebaghi, Priyank Yadav, Dariush Davani, Ryan Huang, Rosanna Weksberg, Paul Delgado-Olguin, KJ Aitken, Darius Bägli

**Author notes:** Martin Sidler and Abdalla Ahmed are of equal importance. Corresponding authors: Darius J. Bägli and KJ Aitken, Developmental and Stem Cell Biology, Research Institute, Hospital for Sick Children, 159420TUV PGCRL Building, 686 Bay Street, Toronto, ON M5G0A4, Canada.

## Abstract

Recently we demonstrated the utility of a nerve-sparing mid-urethra model of partial outlet obstruction (NeMO) that has high consistency and minimal mortalities, unlike the traditional model proximal to the bladder neck. Our goal was to uncover potential therapeutic targets by investigating the genome wide transcriptional changes and pathways altered in NeMO to compare with published human bladder obstruction data. We performed RNAseq and analysed the differentially upregulated and downregulated genes for associated pathways, transcription factor binding site analysis (TFBS), upstream regulators and Gene Set Enrichment Analysis (GSEA). NeMO increased bladder mass, relative bladder mass and hyperactivity, and decreased voiding efficiency. In NeMO vs. sham, 831 genes were differentially expressed (adjusted p<0.05) and correlated significantly with at least one physiologic parameter. Gene ontology revealed an enrichment for matrix pathways in the **up**regulated genes, and for cardiac contraction, oxidative phosphorylation and pyruvate metabolism in **down**regulated genes. TFBS analysis revealed a differential regulation of up vs downregulated genes, with KLF4 strongly associated with the downregulated genes. Downregulated genes of Human bladder obstruction were also associated with the TFBS of KLF4. GSEA of the NeMO gene set confirmed the DAVID results, but also showed a cluster of cytokine activation genes. In human bladder underactive obstruction, cytokines were also highly upregulated. The common cytokine pathway upregulation provided an example of the use of RNAseq for uncovering potential new therapeutic targets. As TNF and the innate immune pathways were strongly implicated in both human and mouse, and TNF is produced by macrophages, we depletion macrophages with clodronate (CL) during NeMO. Although CL did not block hypertrophy, it significantly decreased NeMO-induced hyperactive voiding (p<0.01) and increased voiding efficiency (p<0.05). The expression of several cytokines/chemokines correlated significantly with bladder functional parameters such as residual volumes, and hyperactivity. Conclusions: Gene expression signatures of NeMO were consistent with human bladder obstruction, supporting the use of the nerve-sparing mouse obstruction model for therapeutic exploration.

## INTRODUCTION

Partial bladder outlet obstruction (PBO) has a high prevalence and can affect people of any age and gender. The most common form of PBO results from benign prostatic enlargement (BPE), which affects 70% of men in the US at 60-69 years of age and 80% of men of 70 years or older(1–6). The obstruction-induced remodeling process in the urinary bladder follows a sequence of inflammation, compensatory smooth muscle hypertrophy and tissue fibrosis, and eventually leads to persistent hypertrophy and a functionally decompensated state(7–13). This latter state is marked by symptoms of storage dysfunction (reduction in bladder capacity and compliance), and voiding dysfunction (detrusor overactivity, increased residual volumes, urinary incontinence and incomplete voiding), which together fall under the umbrella term Lower Urinary Tract Symptoms (LUTS). LUTS are seen in many other urologic conditions, including obstructions due to other causes, such as neurogenic (spinal cord injury, myelomenigocoele) and congenital Posterior Urethral Valves.

Models that have attempted to replicate the human disease in mouse have included the hormonally activated models, spinal transections and anatomic obstructions at the bladder neck. However, in mice the latter two approaches have a great deal of variability, with some obstructions too minimal to reproduce the disease phenotype and other obstructions that are too stringent, leading to hydronephrosis and often acute kidney injury resulting in death. This leads to the necessary picking of “suitable” obstructions, often without any accompanying physiological data. To address this, we developed methods for a highly consistent nerve-sparing mid-urethral obstruction (NeMO) (14, 15) and for non-invasive detection of physiologic changes in mice (16). This NeMO model is more reproducible, with low mortality, less injury to shams, but still shows the pathophysiologic changes of detrusor hyperactivity, loss of contractile function and smooth muscle hypertrophy. This contractile dysfunction and hyperactivity is pertinent to human bladder obstruction and UA as well (17–21).

To uncover the global transcriptional responses to PBO and identify novel pathways for potential drug targeting, we performed unbiased transcriptional profiling of NeMO and identified all pathways involved. These pathways included downregulated muscle contractile and oxidative phosphorylation programs, and upregulated AKT, ECM and cytokines/inflammation pathways, as well as shared transcription factor binding sites in our model compared to published human datasets. We also identified activated proinflammatory pathways to be a highly conserved transcriptional response in NeMO and human bladder obstruction(41). We then targeted inflammation mediated by macrophages by chemical depletion, as proof of principle to study RNAseq-derived target effects on bladder function.

## MATERIALS AND METHODS

### Animal Model and Micturition Recording

C57bl/6 mice underwent mid-urethral partial bladder outlet obstruction while another set underwent sham surgery. Micturition patterns were recorded 2 weeks after the surgical procedure. The bladders were harvested for histologic workup and gene-expression analysis by RNAseq after recording of functional parameters. For histology, bladders were embedded in OCT compound while bladders for RNAseq were stored in RNAlater for 24 hours at 4°C before storage at −80°C, then isolation of the RNA with Trizol as previously described(22).

### RNA extraction and RNA sequencing for differential gene expression analysis

RNA from bladder tissue was extracted using Trizol(23) and DNase digestion was performed. RNA quality was determined using the Agilent Bioanalyzer as previously described(24). RNA-seq libraries were made using the NEBNext® Ultra™ Directional RNA Library Prep Kit for Illumina with a starting amount of 500ng RNA. Libraries were sequenced on the Illumina HiSeq 2500. Differential expression was analyzed using DESeq2 with batch correction. Heatmaps showing sample Euclidian distance were generated on R. Heatmaps showing individual gene expression changes were generated using Cluster3.0 and visualized using JavaTreeview. GO term enrichment was performed using DAVID (for Fig.1) and gprofiler (version e98_eg45_p14_ce5b097) with all significant genes as input (adjust p-value<0.05). GO terms encompassing less than 5, or more than 500 genes were excluded from the analysis. Enrichment gene networks were generated by combined both GO term for biological process and molecular function using the Cytoscape plugin Enrichment Map and were annotated using the AutoAnnotate plugin. All labels were edited for readability and comprehension. Pathway analysis was performed on Ingenuity Pathway Analysis (graciously provided as a trial program) using significance values to analyse results. Opossum 3.0 was utilized with default settings for analysis of TFBS in upregulated or downregulated genes. To test frequency of macrophage associated genes from NeMO vs. sham bladders, an *a priori* analysis of macrophage M1 or M2-associated genes(25), was performed. The package ‘biovenn’ on R and *chi*-squared test were utilized for analysis of the frequency of genes in the different groups.

**Figure 1:**
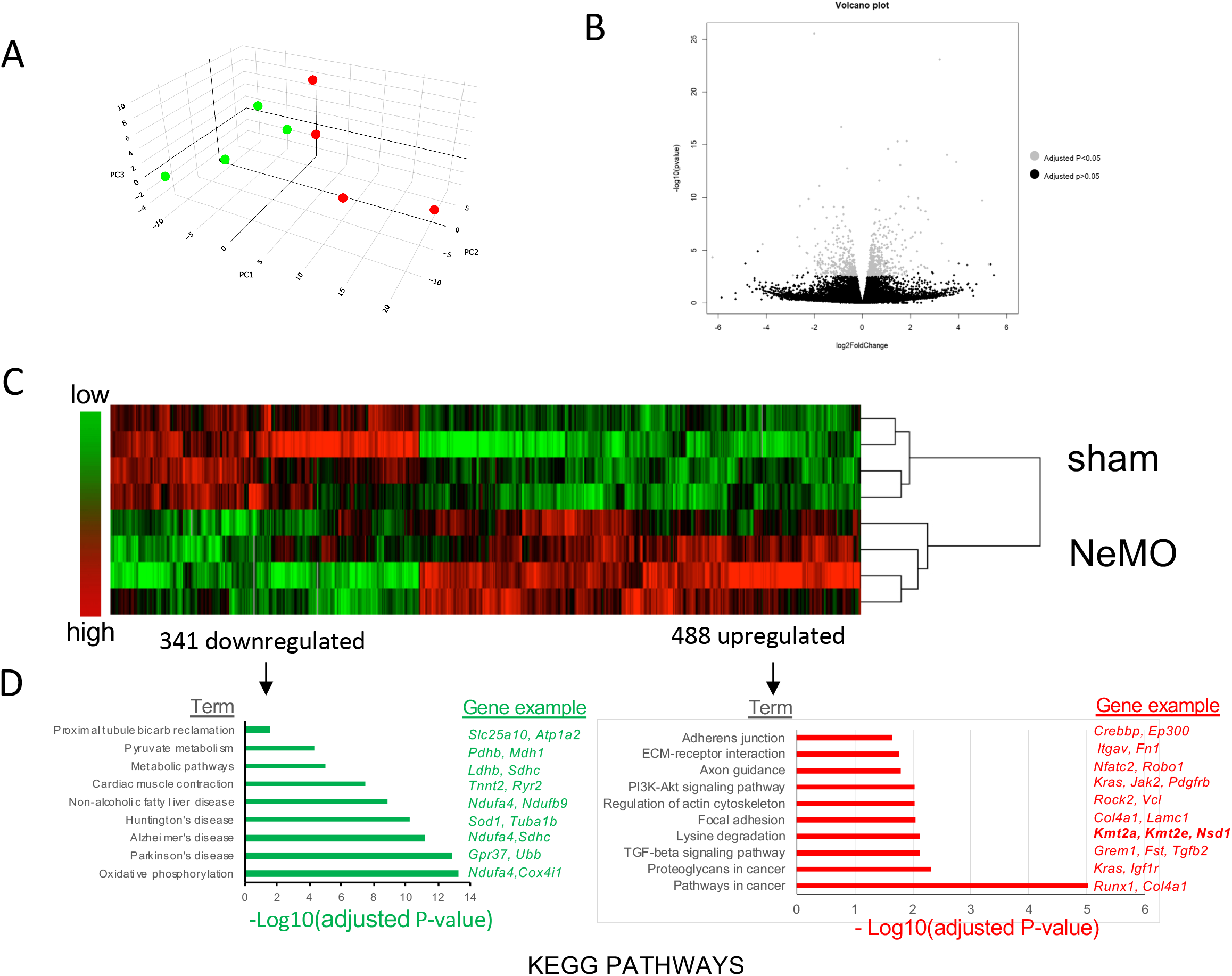
Nerve-sparing mid-urethral partial bladder outlet obstruction (NeMO) induces differential gene expression. (A) Principle component analysis (PCA) of RNAseq reveals separate clustering of NeMO or sham-obstructed (sham) groups. In (B), a Volcano plot of the differentially expressed genes shows the genes that are significant after adjustment for multiple testing (gray dots), adj.p.value p<0.05. (C) A heatmap shows the 831 differentially expressed genes, clustering with 488 upregulated and 343 downregulated in the NeMO bladders vs. sham. N=4 for each group. (D) KEGG pathway analysis of differentially expressed NeMO genes reveals pathways and processes involved in obstructive disease pathogenesis. GO term enrichment analysis revealed that up regulated genes were highly associated with connective tissue formation and remodeling, TGFB, PI3K/AKT, ECM receptor interaction by KEGG pathway analysis. Differentially expressed genes that were downregulated in PBO involve cardiac contraction, both pyruvate <u>and</u> oxidative metabolism.

### Non-invasive Bladder Function

Mice were placed in metabolic cages with a wire grid during the light-cycle (i.e. during daylight) with free access to food and water. After allowing for 1 hour of acclimatization, voiding was assessed non-invasively by continuous recording of voided urine by a weigh scale placed under the grid, for 10 hours. Urine weight increases of > 50μg (50uL by volume) were considered small voids. The largest void recorded was the maximal voided volume and the average voided volume was the mean of all the voided volumes. Recorded weight-traces were analyzed using a software-algorithm that detected weight increase and discriminated fluid from solid material based on weight decrease by evaporation of urine. Thereafter, mice were anesthetized, bladders were removed intact with bladder neck secured closed. Bladders plus neck were first weighed on a fine scale with the bladder neck closed. After removal of bladder neck and emptying the bladder, bladders and neck were re-weighed. The residual urine was determined by the difference in bladder mass containing urine and bladder weight without urine, expressed as a volume. Maximum bladder capacity was the result of the maximum voided volume plus residual volume. Mean or maximum voiding efficiencies were calculated from the mean or maximum voided volumes, respectively, divided by bladder capacity. Other calculated parameters included ratio of residual volume/bladder capacity as a measure of retention, and ratio of number of small/total voids, which serves as a surrogate of bladder hyperactivity.

### Clodronate treatment

Female C57/Bl6 mice of 18-21 g body mass underwent NeMO at the mid-urethral level or sham. Of these mice, 13 NeMO and 13 sham were randomized to 50 mg Clodronate Liposomes (CL, Amsterdam, Netherlands) per kg body mass, 3 times per week intraperitoneally. The remaining 14 NeMO and 12 sham mice received Normal Saline (NS; 0.9% NaCl) injections IP. After 2 weeks, animals underwent micturition recording as above. Micturition physiology measurements could not be effectively measured in three mice, though their residual volumes, bladder and body weights were measured. Three clodronate-treated mice were sacrificed two days early due to weight loss, a subcutaneous tumour and signs of illness, though no UTI was evident.

### Immunofluorescent staining for smooth muscle actin and macrophages

Mid-equatorial bladder cryosections of 7μm were fixed in 4% paraformaldehyde (PFA) 20 min, then permeabilized with a 0.2% Triton-X solution in PBS for 10 min then washed in PBS (26). Sections were blocked in 5% donkey serum, 5% goat serum, 0.3% BSA, 0.3M Glycine solution in PBS for 1 hour before incubation with primary antibodies: rat anti-mouse-F4/80 (DVS Sciences, Sunnyvale, California, USA) to identify macrophages; anti-smooth muscle actin (1:200; Sigma, St. Louis, MO, USA). Secondary antibodies (donkey anti rabbit AlexaFluor® 488, and goat anti rat Alexa 594; Jackson ImmunoResearch Laboratories, West Grove, PA, USA), were incubated at 1:400 dilution in 1% BSA with 5% goat and 5% donkey serum in PBS solution, 4°C overnight. Nuclei were counterstained with Hoechst 33342, or DIC micrographs were taken alongside to define tissue regions. Micrographs were acquired on a Zeiss Axiovert Spinning disk confocal microscope with Volocity (Version 6.3; PerkinElmer Inc., Woodbridge, ON, Canada). Pseudo-colour imaging was utilized to optimize visual contrast. Muscle regions were defined by myosin staining and morphology. Macrophage quantity in smooth muscle marker positive regions were quantified using Volocity.

### Cytokine Array

Harvested bladders were homogenized using a the loose setting of a Dounce homogenizer in PBS + Complete Protease Inhibitor (Sigma). Extracts were centrifuged at first 500g, then 2000g to remove cells and debris. Protein content in supernatant quantitated using the Pierce BCA Protein Assay. Cytokines and chemokines were screened using an array kit (Proteome Profiler Mouse Cytokine Array Kit, R&D Systems, McKinley Place, MN, USA), according to the manufacturer’s protocol; resulting films were scanned and pixel intensity was quantitated using Image J.

### Statistics

Physiologic, immunostaining and cytokine data were analysed by type III ANOVA, Shapiro-Wilk’s normality test and Levene’s test for homogeneity of variance to establish the appropriate post-hoc test (Student’s, Welch’s t-test or Kruskal-Wallis test). P-values less than 0.05 were considered significant. Quantitative results from physiology were summarized graphically using boxplots, the box encompassing the interquartile range (IQR) from the first quartile (Q1) to the third quartile (Q3), the bold transverse bar representing the median. The whiskers mark maximum and minimum values. *X^2^* tests were performed using ‘R’. Pearson’s correlations and heatmaps were performed on ‘R’ using ggcorrplot to correlate physiologic data with expression and cytokine data. Statistical analysis and plotting of functional data was performed using IBM SPSS Statistics 24 (SPSS Inc., Chicago, IL USA) and the ‘R’ ggplot and boxplot() base function.

## RESULTS

### *RNAseq reveals a set of 831 differentially expressed genes in* nerve-sparing mid-urethral *obstruction (NeMO)*

We previously established the NeMO model of obstruction in mice, which showed decreased contractile efficiency coupled with hypertrophy, increased residual volumes and overactivity. To uncover the transcriptional response to partial obstruction in the bladder, we performed the NeMO model in adult female mice (14, 15), non-invasively measured physiology, then isolated the bladder dome for RNA-seq analysis. PCA plots demonstrated that the transcriptional profiles of the sham-operated bladders clustered separately from NeMO, by Principal Component Analysis of the two groups (Fig. 1A). A total of 831 genes demonstrated significantly different gene expression profiles in NeMO vs. sham bladders, that clustered separately using Euclidean distance into 488 upregulated and 343 downregulated genes (Fig.1B,C, Table 1, Supplemental Table S1), highlighting the reproducibility of the NeMO model.

**Table 1:**
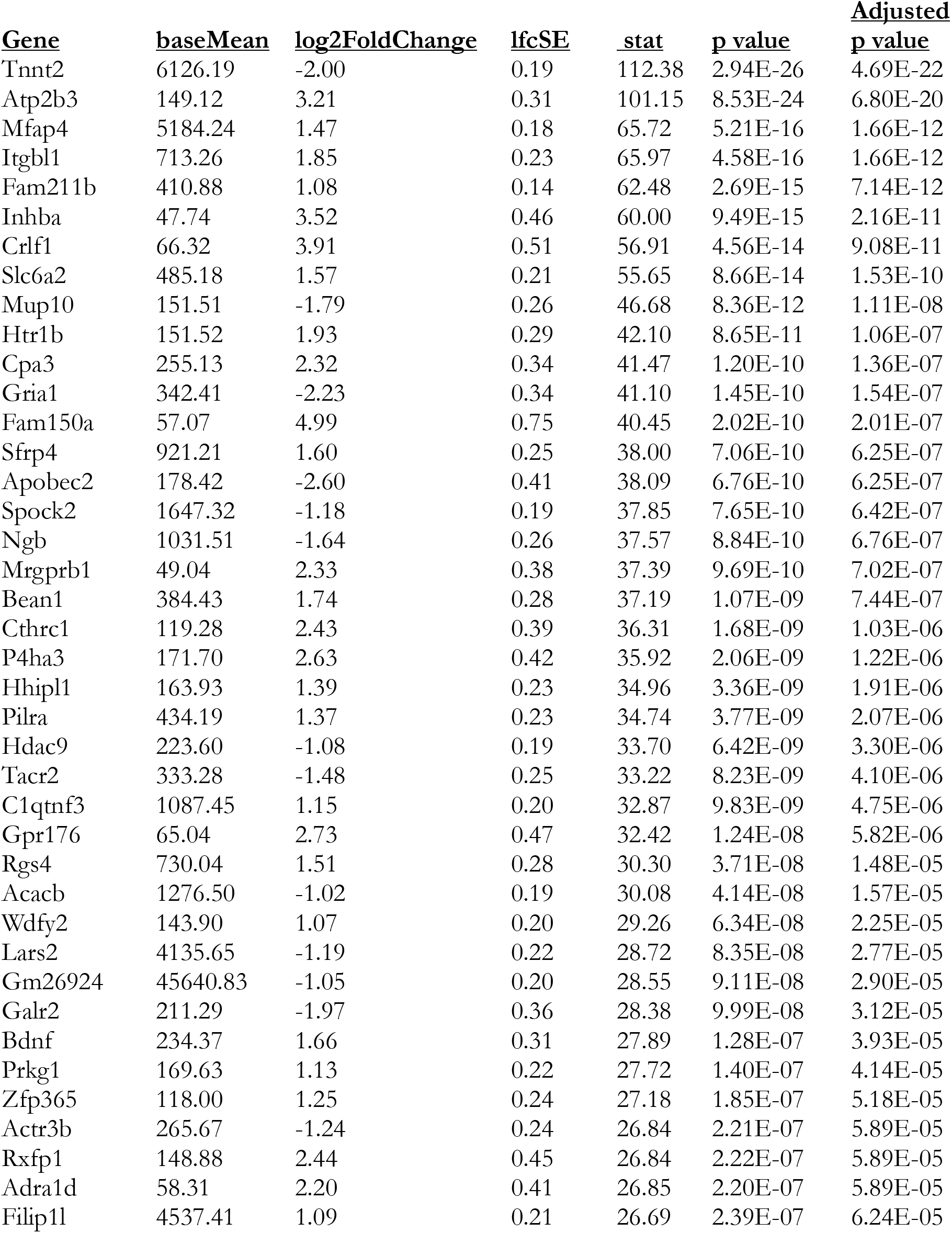
Top 40 differentially expressed genes in NeMO vs. sham by RNAseq analysis. RNA was sequenced by Illumina’s Mouse Ref-8 v2 Beadchip and analysed using Illumina BeadStudio software. Differentially expressed genes between biological groups were identified after adjustment with a false discovery rate of 0.01, adjustment for multiple testing. For this Table, BaseMean (average counts) >10, and logFold changes >1 or < −1 are shown. Complete data for above comparisons can be found in Supplemental Tables 1 and 2, and NCBI GEO. Adjusted p values less than 0.05 were considered significant.

### Gene ontology (GO) term enrichment of revealed distinct pathways

The 341 downregulated and 488 upregulated genes were associated with unique KEGG pathways in NeMO (Fig.1D). Downregulated pathways included oxidative phosphorylation, metabolism pathways and pyruvate metabolism. In addition, Alzheimer’s, Huntington’s and Parkinson’s and non-alcoholic fatty liver disease pathways were enriched in the downregulated set, but the genes in these pathways were shared with oxidative phosphorylation, including many *Nduf*-genes. Intriguingly, the term cardiac muscle contraction was enriched in downregulated genes, including Ryr2, Myl4(myosin light chain4), Tnnt2 (cardiac troponin T2) which also had high mean counts (Table 1, S1). From the upregulated set of genes, ECM and ECM receptor pathways were highly enriched (Fig.1D), comprising TGF beta signaling, Focal Adhesions, ECM-receptor interaction and Adherens junction. Axon guidance also was upregulated due to the presence of genes such as Nfatc2 and Robo1. Interestingly, Lysine degradation was also upregulated, including two lysine methylation enzymes Kmt2a and Kmt2e, and two dioxygenases, Plod and Bbox1. Two Cancer pathways were also enriched, possibly due to their association with ECM and growth pathways. ‘Regulation of actin cytoskeleton’ was also enriched with genes such as Vcl (vinculin), Integrins alpha-2,-6 and -v, and Moesin. The PI3K/AKT pathway was also increased, which is expected as it has been studied in bladder smooth muscle. PDGFRB, which is a highly SMC-selective membrane receptor, was found upregulated in both ‘Regulation of actin cytoskeleton’ and ‘PI3K/AKT signaling pathway’.

### Voiding dysfunction and gross pathology was increased in NeMO mice and associated with Gene expression changes

NeMO mice were significantly increased in hyperactivity, retention ratio (residual volume/bladder capacity), bladder mass and bladder to body mass ratio, and decreased in maximum and mean voided volumes, and voiding efficiencies (Fig.2A). These physiologic parameters were significantly correlated with each other (Fig.2B), but also with at least one gene (either positively or negatively, Fig.2C). Many genes were correlated with gross hypertrophy, efficiency and overactivity, and each had unique genes, thought efficiency and hypertrophy showed a high degree of overlap.

**Figure 2:**
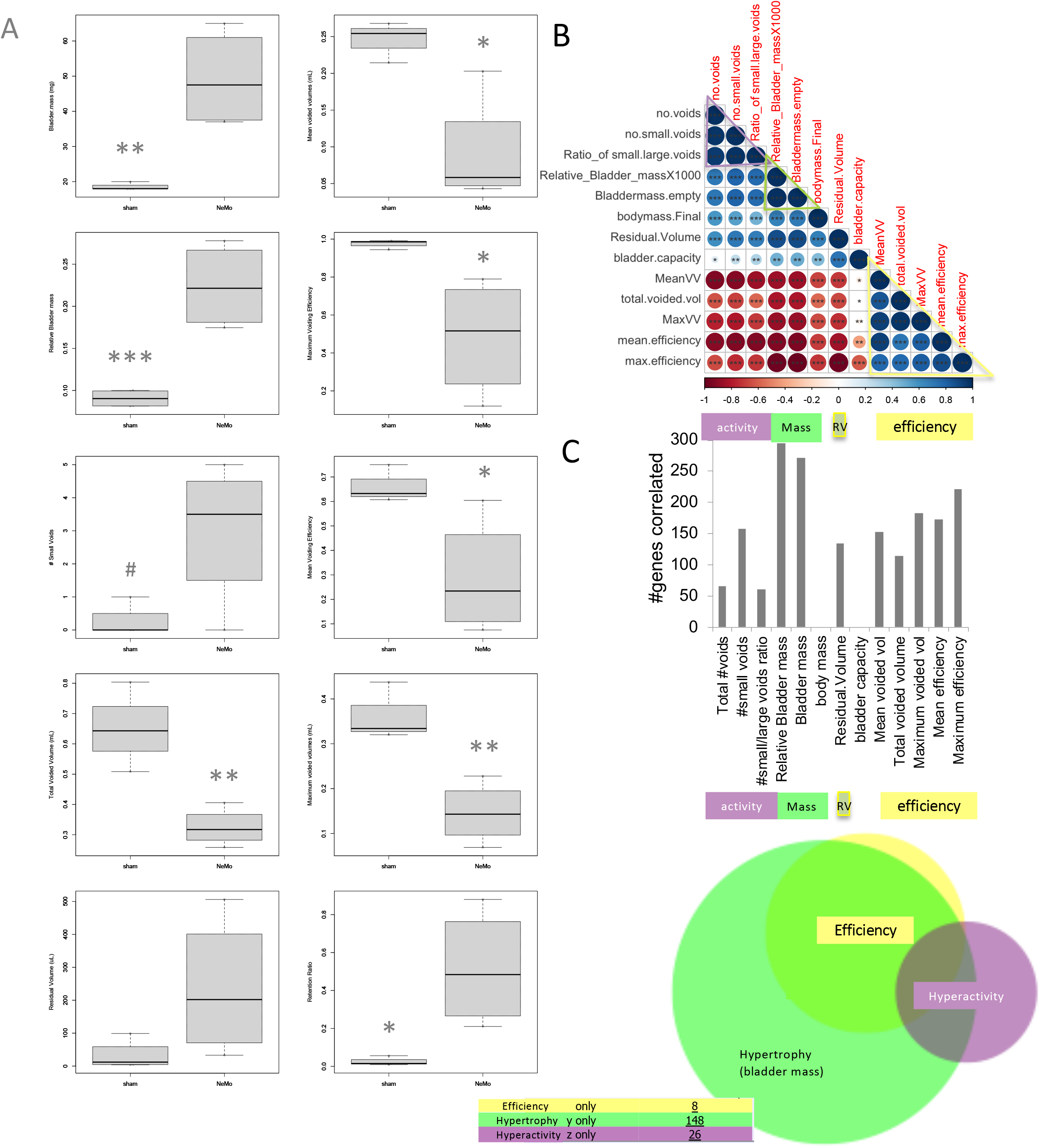
Physiologic changes and their correlations with gene expression. Bladder function of WT mice that underwent sham or NeMO surgery and were placed in metabolic cages for 11 hours to record voiding patterns and volumes. One day later bladders were harvested and residual volumes, body and bladder masses recorded. In A, we see that many physiologic aspects of NeMO were highly significantly different compared to sham. n=4 for these groups. Most physiologic aspects were had very strong correlations either in a positive or negative direction, though bladder capacity and body weight had very weak correlations. Most physiologic parameters were also highly correlated with at least one gene expression profile (C). With bladder mass and bladder to body mass ratio (or gross hypertrophy) have the largest number of gene significant correlations.

### Predicted transcription factor binding sites (TFBS) for up or downregulated genes of NeMO were distinct

To understand how the 831 genes may be regulated in NeMO, upregulated vs. downregulated genes were analysed on Opossum 3.0 for transcription factor binding sites (Fig.3). The TFBS for KLF4, MZF and SP1 were associated with downregulated genes, and 3 related FOX’s, SRY, HOXA5, ARID3A and the cardiac NKX2-5, were associated with upregulated genes(Fig.3A).

**Figure 3:**
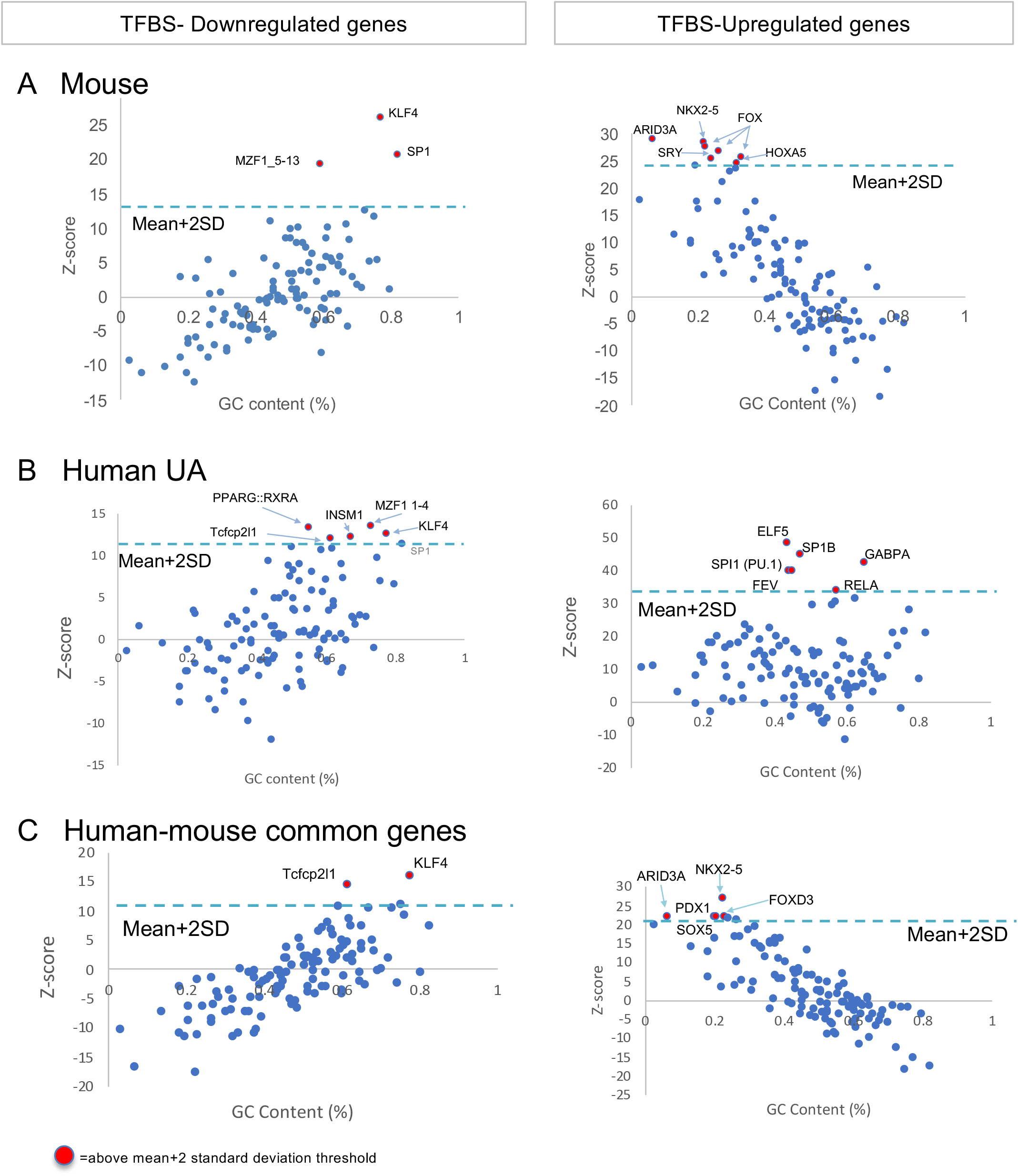
Transcription factor binding site (TFBS) were differentially regulated in downregulated vs upregulated genes in obstruction with common TFBS in mouse and human. We analysed TFBS on Opossum 3.0 in down and upregulated NeMO genes, identifying KLF4, and NKX2-5 sites as the top TFBS, respectively. We also examined human TFBS using published datasets, and found that human underactive bladder obstruction (UA) had similar TFBS for the downregulated genes, including KLF4 and MZF, as in mouse. However, the upregulated TFBSs included many myeloid/immune regulators including SPI1 (PU.1) and RELA. We examined our mouse data for human orthologs in common and found 100 upregulated and 60 downregulated genes. These up and downregulated genes in common had the very similar TFBS to mouse alone, including sites for KLF4 (down), and NKX2-5 and Fox factor (up).

### Patient and NeMO RNA expression patterns showed concordance

While the transcriptional responses to NeMO showed a high reproducibility and consistency within groups at the gene expression level, it is still unclear if this model closely recapitulates human PBO at the transcriptional level. To assess the similarity between our NeMO model and human PBO, we cross-referenced the transcriptome of mouse NeMO that we generated against published transcriptomes of human PBO that resulted in bladder underactivity. Of the 831 genes differentially expressed in mouse NeMO, 796 had human orthologs. Analysis of these publicly available transcriptomes of human PBO(27) using the same bioinformatic parameters of our analysis on mice bladders revealed that 160 (20.1%) of the 796 genes were dysregulated in both human PBO and mouse NeMO (Supplemental Table S2), with 100 upregulated and 60 downregulated. The TFBS analysis of the human UA downregulated genes showed a similar association with KLF4, and MZF, but also other factors (Fig. 3B). The upregulated UA TFBS were associated mainly with inflammatory genes, including the macrophage/monocyte regulator SPI1 (Pu.1) and RELA. Interestingly the common dataset of 160 genes had 100 upregulated and 60 downregulated genes. The common dataset TFBS’s (Fig.3C), resembled the mouse (Fig.3A) with KLF4 for the downregulated genes and NKX2-5 for the upregulated genes.

We next examined mouse and human pathways using another method ‘gprofiler’. Our NeMO mouse model showed that ECM, tissue remodeling and cytokine activity were amongst upregulated processes, and muscle contractile processes and oxidative metabolism were amongst downregulated processes (Fig.4A). The genes upregulated in human PBO were predominantly related to immune cell migration, differentiation, and cytokine secretion (Fig.4B). Downregulated genes were enriched for functions related to amine transport, muscle contraction and oxidative metabolism (Fig.4B). GO term enrichment analysis on the commonly upregulated genes identified cytokine receptor interactions to be the most highly enriched term (data not shown). “TGFB signaling pathway” was the only enriched KEGG pathway (adjusted P-value= 0.014) for human PBO alone (data not shown). This suggests immune activation and TGFB signaling play conserved roles in the response to bladder obstruction and may serve as targets for therapy.

**Figure 4:**
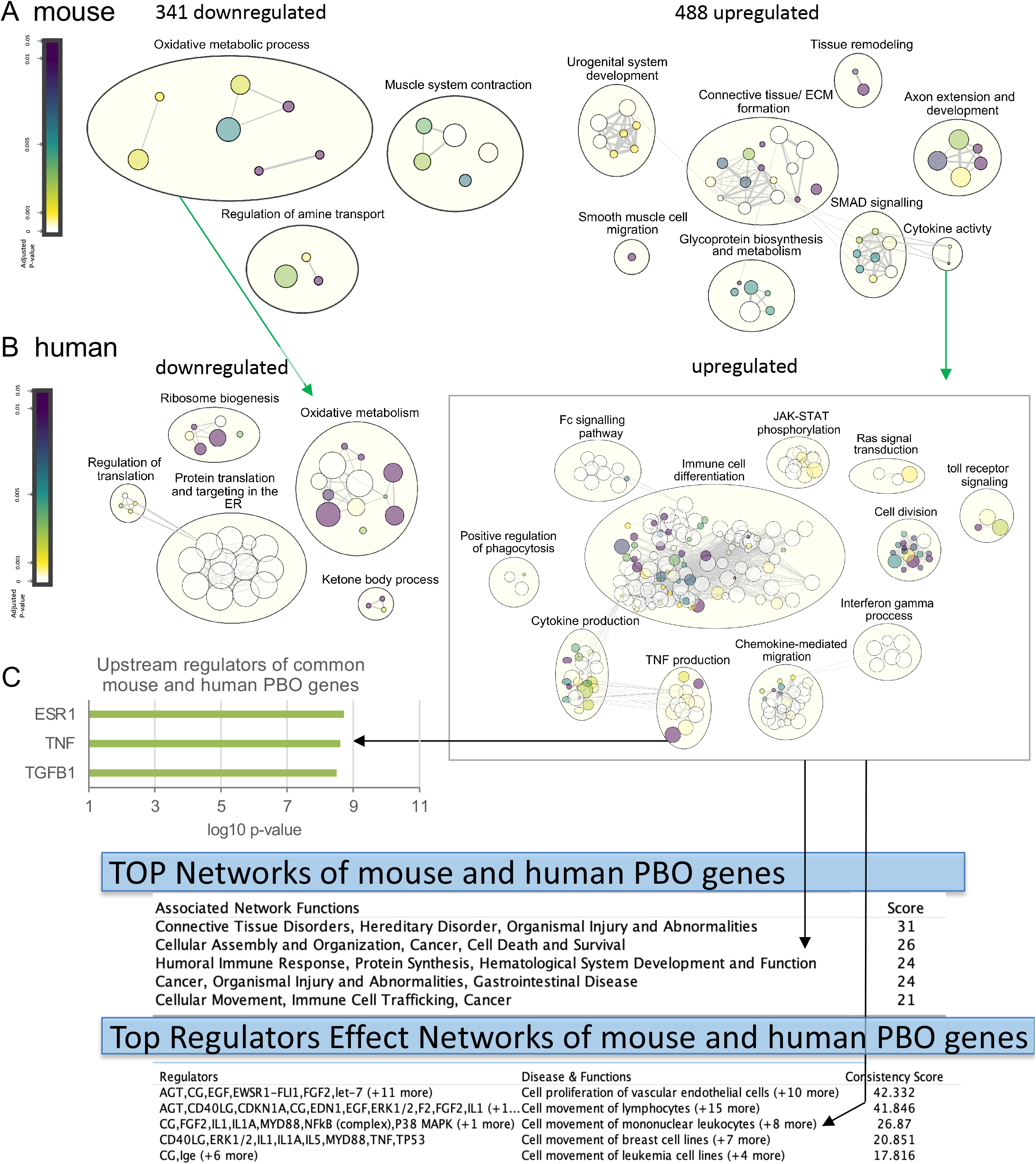
Gene set enrichment analysis in upregulated vs downregulated genes from NeMO and Human obstruction reveals genes and pathways in common. (A) Mouse genes were examined using ‘gprofiler’ and Cytoscape. Mouse pathways were increased similar to KEGG analysis (as in Fig.1), but also including cytokine activity. (B) GO term enrichment was examined in upregulated human PBO genes, which revealed pathways involved in phagocytosis, JAK-STAT signaling, inflammation, TNF and immunity. GO term enrichment for downregulated human PBO genes revealed oxidative metabolism and translation mechanisms, similar to mouse. Both mouse and human PBO genes were associated with terms for cytokines and oxidative phosphorylation (see green arrows). (C) Ingenuity Pathway Analysis (IPA) of differential gene expression in both NeMO and human PBO revealed the Top upstream regulators to be ESR1, TNF and TGFB1 with high significance, (p<10^−8^). Similarly, IPA also revealed potential pathway connections related to connective Tissue, leukocytes and Humoral immune response.

Importantly, Ingenuity pathway analysis (IPA) of the genes conserved between mouse NeMO and human PBO (UA) also showed the upstream regulators of TNF and TGFbeta, p<10^−13^ (Fig.4C) and several networks with both ECM, TNF and immune functions (Fig.S1), similar to published human PBO data(27). In addition, IPA Disease Associations included Immunological Disease with a significance between 3.15×10^−4^ - 2.13 ×10^−11^, and included genes JAK2, ABL2, CXCL5, CD209. The top shared networks comprised: (1) connective tissue disorders, (2) cellular assembly and organization, cancer, cell death and survival, (3) **<u>humoral immune response</u>**, protein synthesis and **<u>hematological system</u>** (Fig.4C). These immune pathways included a mutually dysregulated group of inflammatory genes, *Jak2, Cd80, Cxcl6* (*Cxcl5* in mouse), *Crlf1, Ccrl2, Il13ra1* and *Ighg3*.

### Proof of principle targeting of pathway identified through RNAseq

Analysis of the transcriptome of mouse and human obstructed bladders together suggested that connective tissue disorders, TGFB and activation of the immune response are a conserved response to bladder obstruction. As ECM and TGFB are already well studied topics in bladder obstruction, we decided to explore the cytokine response, as an example of the utility of the RNAseq for uncovering pathologic mechanisms. As cytokines and TNF were implicated bioinformatically, in **<u>both</u>** human and mouse datasets, and macrophages are known to be significant producers of TNF, we investigated whether macrophage/monocytes gene expression profiles might also be associated with our data. A macrophage associated profile (25) was tested for enrichment in NeMO vs sham (Figure 5A). Macrophage-associated genes, such as Arg1 (M2), iNOS and CD80 (M1) were enriched, with macrophage associated genes occurring more frequently than expected in the upregulated NeMO genes, p<0.0005 by *chi*-squared test. Next, as a proof of principle to explore mechanisms and potential treatments based on RNAseq-derived targets in PBO, we depleted the macrophages by clodronate liposome (CL) treatment in sham or NeMO mice. Signs of infection were absent in treated mice. Immunofluorescent staining confirmed an increase of macrophages (F4/80^+^) in NeMO bladders, which was blocked by clodronate treatment (~2.5-fold reduction, p<0.05, Fig.5B,C, Supplemental Fig.S2). CL-induced depletion of macrophages improved specific aspects of bladder function. Mean voiding efficiency of NeMO bladders was significantly restored with CL treatment, p<0.05. Also, CL treatment of NeMO led to a ~50% reduction in residual volumes and hyperactivity, p<0.05, 2-tailed t-test, Fig. 5D). Bladder mass (Fig.5D), and voided volumes were not affected by CL (Fig.S2). To investigate if cytokine secretion was affected by CL, proteins were extracted from fresh bladders and tested by cytokine protein array. IL17,TREM1, CXCL9 were found to be increased in NeMO (Fig.5E). Significant reductions in IL17,TREM1, CXCL9 and TNF were observed with CL treatment (Fig.5E, p<0.05).

**Figure 5:**
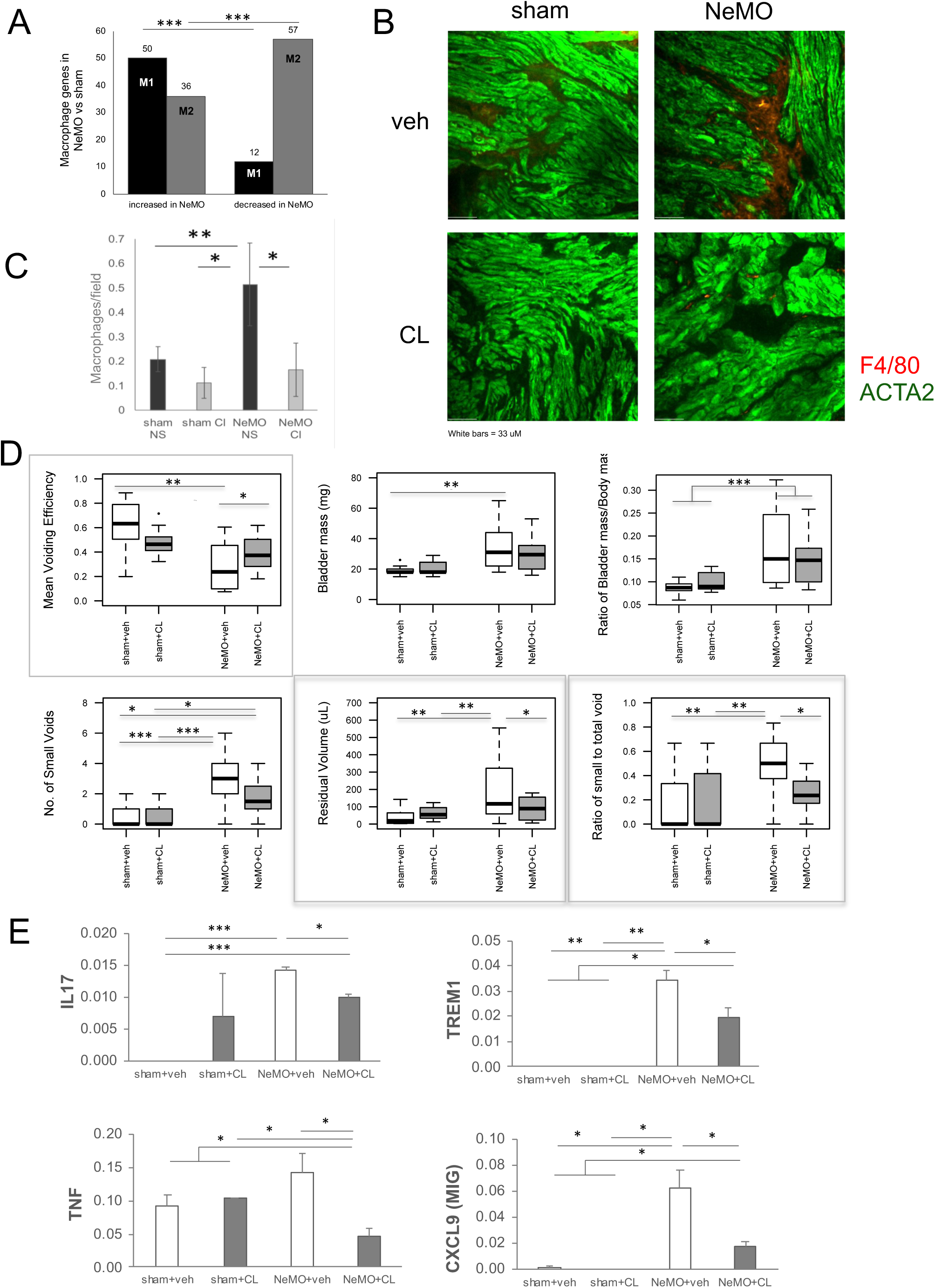
Macrophage-dependent changes in NeMO. (A) Macrophage associated genes (both M1 and M2) occur more frequently than expected enriched in the RNAseq dataset, *chi*-sq<0.0001. (B) Macrophage infiltration is increased during NeMO, but reduced by clodronate treatment (CL). Immunofluorescent staining for F4/80, a marker of macrophages, were visualized in cryosections of the bladder with and without CL. (C) Bladder function, which is lost during nerve-sparing mid-urethral obstruction (NeMO) vs. sham, improves with clodronate (CL). NeMO increased hyperactive voiding (ratio of #small to total voids, Kruskal-Wallis rank sum test, p<0.005), and residual volumes (Welch’s t-test, p<0.005). The NeMO increase in volume and hyperactivity was reduced by CL (p<0.05). NeMO caused a decrease in mean voiding efficiency (Student’s t-test, p<0.05), which CL significantly increased (Student’s t-test, p<0.05). (F) The ratio of the Residual volumes (RV)/bladder capacity is an indication of retention and was significantly increased during NeMO with or without CL (Student’s t-test, p<0.005). (G, H) Compared to sham, NeMO led to >2-fold increase of bladder to body mass ratio and absolute bladder mass (Welch’s, p<0.005), which were similar between NeMO+vehicle vs. NeMO+CL. Sham n=11, sham+CL n= 11, NeMO n=13, NeMO+CL n=12. Statistical analysis was performed using ANOVA with post hoc t-tests as indicated above. 2-tailed (*) and 1-tailed (**) t-test, *p*<0.05, ***, 2-tailed t-test *p*<0.005. **E)** Cytokine expression, altered by NeMO and partially restored by clodronate treatment (CL), correlated with bladder function during nerve-sparing mid-urethral obstruction (NeMO). (E) By secreted protein array, NeMO increased TREM1, CXCL9 and IL17 expression. CL treatment of NeMO reduced TREM1, CXCL9, IL17, TNF expression. We also found discrete Pearson’s correlations (*R*-value) between secreted factors and bladder function in sham, NeMO +/-CL (Supplemental Fig.S3). The secreted factors appeared to correlate with together in 3 groups (I, II and III), by hierarchical clustering. Secreted factors correlated strongly with functional parameters affected by CL, including: small to total voids ratio (hyperactivity) – TNF, residual volumes - IL17; bladder capacity -TNF, TREM1, IL17, CXCL9. *, p<0.05; **, p<0.01, ***, p<0.005. n= 4.

## DISCUSSION

Current treatment for various forms of PBO are purely symptomatic, leaving the principal causes unaddressed. For instance, hyperactivity occurring in most human PBO can be managed with anticholinergics or adrenergic treatments to reduce spontaneous contractions and high frequencies, but the etiology of the hyperactivity is less understood. To non-subjectively uncover the underlying changes in a consistent mouse model of PBO (NeMO), we determined the genome-wide expression changes during mouse PBO. Our NeMO model shows multiple changes in bladder function, including gross hypertrophy, voiding efficiency, residual volumes and hyperactivity (Fig. 2,5), similar to changes seen in human PBO, in particular UA. Other murine models of PBO have the difficulty of high PBO mortality rates as well as variability in both PBO and sham groups, potentially obscuring gene expression differences. There is high variability and high mortality in traditional obstruction as the bladder neck and ureters are minute and susceptible to damage during surgery. By measuring bladder function non-invasively in NeMO(15, 16, 28), we could determine the efficacy of treatments without a high baseline mortality rate or secondary effects of sham surgeries.

### NeMO shows similarity to human PBO

RNAseq analysis of NeMO revealed a unique group of 831 gene changes with enrichment for pathways related to inflammation, ECM and loss of enrichment for oxidative phosphorylation and muscle contraction (Figs.1,4,S2). While these pathways are common to chronic diseases of other organs, the specific collection of gene changes may be unique to the bladder. Importantly, some expression changes were conserved across species, as 160 or 20.5% of the orthologs that were differentially altered genes in the mouse PBO were shared with human PBO (underactive-UA). Several NeMO functional parameters were similarly altered in human UA. RV of obstructed mice was 3-fold of shams, which is consistent with findings in other rodent studies(22, 29, 30). Measurements of total voided volumes and number of micturitions indicated that the bladders had voided in the preceding measurement period, and were not in retention. Indeed, the bladder capacity exceeded that of the residual volumes in all animals (Figure 2,5), although the ratio of RVs/bladder capacity (retention ratio) in the vehicle NeMO was higher than in shams (p<0.05, Figure 2). This is consistent with human underactive bladders that demonstrate a high retention ratio, alongside other features of obstruction(27).

### ECM pathways

Analysis of mouse NeMO RNAseq data using KEGG, gene ontology, and IPA revealed pathways for ECM genes, metabolic, oxidative, inflammation and muscle development (Figure 1,4,S2). Previous work by us(22, 29), and others(10, 11, 31–41) have shown the crucial role of ECM in PBO pathology. Here we determined that matrix gene expression is enriched in NeMO, including FN1, TGFB2, collagens type I, III, IV, and lesser known ECM-related genes, such as Fst, *Itgav* and Col11a1 (Supplemental Table S1), many which were upregulated and identified by KEGG pathway analysis. Interestingly, Col11a1, FN1 and denatured collagen contain RGD or cryptic RGD peptide sequences that can bind alphav beta3 integrin to induce signaling(42, 43). This is of interest as alphavbeta3 integrin-ECM receptor signaling mediates matrix- (24, 44) and stretch-dependent responses in BSMC (45) We also noted that PI3K/AKT and ECM-integrin interact in the IPA networks (Supplemental Figure S1).

### Both Oxidative phosphorylation and pyruvate metabolism are downregulated pathways

Oxidative phosphorylation is downregulated in NeMO bladders, which affects the neurologic disease pathways listed in Fig. 1D through loss expression of key mitochondrial respiration chain genes. Interestingly, downregulated genes are also enriched for the pyruvate metabolism genes, potentially altering the energetic status of the bladder.

### Myogenic program genes were downregulated

While we expected downregulation of myogenic program gene, including SMC markers, some lesser known genes appeared here. The downregulation of muscle and cardiac genes in NeMO (Figure 1,4) included a highly significant transcriptional downregulation of two troponins, TNNT2 and TNNC2 (p<10^−6^, Table 1), RyR2 and many other genes. The troponins have been studied primarily in cardiac and skeletal muscle, but in the bladder they have been associated with development of myelomeningocoele(46) and contractile force generation (47). The contractile proteins downregulation occurred alongside upregulation of the actin-cytoskeletal proteins. This could suggest that pathways regulating mechanical contractile force are being replaced with mechanical tension in obstruction.

In addition, the TFBS analysis associated KLF4 with the downregulated genes. The role of this gene in bladder SMC phenotype is also not clear, although in vascular biology it has a clear role in the regulation of SMC phenotype, through interactions with histone deacetylases (HDAC) and other factors binding that CAARG element of SMC genes (48–50), including many contractile proteins. The KLF4 TFBS was also conserved in the downregulated mouse and human UA genes. Together these changes could underlie reduced efficiency and overactivity seen during PBO.

### Inflammatory genes and pathways were upregulated in the PBO bladder

GO term analysis indicated a significant enrichment of cytokines. Inflammation has been previously associated with hyperactivity in both human studies(51) and experimental models that mimic overactive bladder(52, 53). Here however it seems to be associated with underactive bladder as well. It is likely that similar pathways are activated in other forms of human PBO (44). However, the large variability in available patient PBO and DO RNAseq data precluded their comparison with NeMO. By ingenuity pathway analysis, the master cytokine TNF was supported as a potential upstream regulator of the significantly altered genes in NeMO(Fig.4). TNF protein has been reported to rise in the bladder and serum in myopathic bladder models(54–56). TNF protein is produced in high amounts by macrophages and dendritic cells, and can stimulate the production of other chemo/cytokines (57) Therefore, it is not surprising that TNF homologs or TNF receptors, such as the TNF superfamily genes Tnfaip8l3 and Tnfrsf22 are significantly increased during NeMO (Supplemental Table S1). TNF was also previously identified as the top upstream regulator(27) in <u>human</u> PBO similar to our merged dataset, lending support to the nerve-sparing female mouse obstruction model.

### Depletion of macrophages in bladder obstruction

Several NeMO parameters, residual volumes, ratio of number of small to total voids (hyperactivity), bladder capacity and mean voiding efficiency are reversed by CL (p<0.05, Figure S3), while other parameters (gross hypertrophy and voided volumes, Figure 5D, Supplemental Figure S3) are not. As a result, macrophage-dependent inflammatory mechanisms distinctly affect hyperactivity, mean efficiency, capacity and residual volume separately from other aspects of obstructive dysfunction.

The improvements in bladder function due to CL treatment (Figure 5) were consistent with the reduction in TNF, CXCL9, TREM1 and IL17 <u>proteins</u> (Figure 5E). These chemo/cytokines have also been implicated in causing myofibroblast activation, SMC contraction and SMC secretory functions(56, 58–65). Interestingly, *Crlf1, Ccrl2* and *Il13ra1* (macrophage associated genes) were upregulated in both mouse and human obstruction, suggesting a conserved inflammatory response across species, which might be leveraged to uncover discrete mechanisms of human PBO. Macrophages have additional functions in remodeling the ECM through their phagocytic and enzymatic activities (66–70). TNF induces the secretion of MMPs and elastin-binding protein in smooth muscle cells, potentially linking ECM to cytokines(71). Also, chemokines produced here can activate integrin receptors(72, 73), which could synergize with ECM ligands. However, CL did not alter the gross hypertrophy of the bladder in NeMO bladders, suggesting that other mechanisms may regulate gross hypertrophy.

<u>In summary</u>, PBO in mice is associated with changes in contractile, inflammatory, matrix and AKT pathways, alongside changes in hyperactivity and voiding efficiency. Upstream regulators and many transcription factor binding sites appear to be governed similarly in human and NeMO bladders. Macrophage-dependent cytokines were associated with several pathophysiologic features of obstructive disease, similar to the inflammatory pathways upregulated during human PBO (27). We found that modulating inflammation through clodronate treatment influenced cytokine expression, leading to improvements in discrete aspects of bladder function, and opening new avenues for therapeutic approaches. Other pathways and regulators are implicated here, such as AKT activity, ECM, SMC differentiation and contractile function, dysregulation of oxidative metabolism and smooth muscle/myogenic genes, but remain for further exploration. The specific links between inflammation, fibrosis, AKT and an overactive micturition pattern can also be explored as future work. With this model, we now have performed a crucial step to finding new therapies, with the definition of gene expression changes in a useful model and in common with human PBO.

## Acknowledgements

We would like to acknowledge the Sickkids Imaging Facility, the support of the CIHR operating grants (D. Bägli, P. Delgado-Olguin), Institute of Medical Sciences, University of Toronto (M. Sidler) and Peri-Operative Services, Sickkids (M. Sidler).

## AUTHOR CONTRIBUTIONS

R. Weksberg, P. Delgado-Olguin, K.J. Aitken & D. Bägli designed research; M. Sidler, A. Ahmed, K.J. Aitken, D. Davani and D. Koshkebaghi analyzed data; M. Sidler, A. Ahmed, J-X. Jiang, D. Davani and K.J. Aitken performed the research; M. Sidler, K.J. Aitken, A. Ahmed, P. Delgado-Olguin & D. Bägli wrote the paper; M. Sidler developed software necessary to perform and record experiments.

## Supplementary material

RNAseq data can be found at NCBI GEO.

## Notes

### Competing Interest Statement

The authors have declared no competing interest.

### Summary of Updates

The revision corrected a margin issue in Figure 5.

