## Supplemental Materials for "Human - murine concordance of molecular signatures in nerve-sparing murine partial bladder outlet obstruction (NeMO)"

**Figure S1:** Ingenuity pathway analysis (IPA) of the differentially expressed genes in our mouse NeMO vs. sham bladders were similar to those published in human(44). These included gene networks involving ECM regulation (A-D), inflammation (B,C), ERK-collagen (A), PI3K(B), NFKB-MMPs and -collagens (C) and AKT-integrin (D). IPA also generated a TNF-centred pathway that show the connection to FN1, MMP2 and Col4A1, which are highly upregulated during NeMO (E). (F) **Connected** networks from Ingenuity pathway analysis (IPA) in genes common to partial outlet obstruction in humans, and the Nerve-sparing mid-urethral obstruction (NeMO) model in mice, included PI3K, STAT3, inflammatory and TGFbeta pathways.

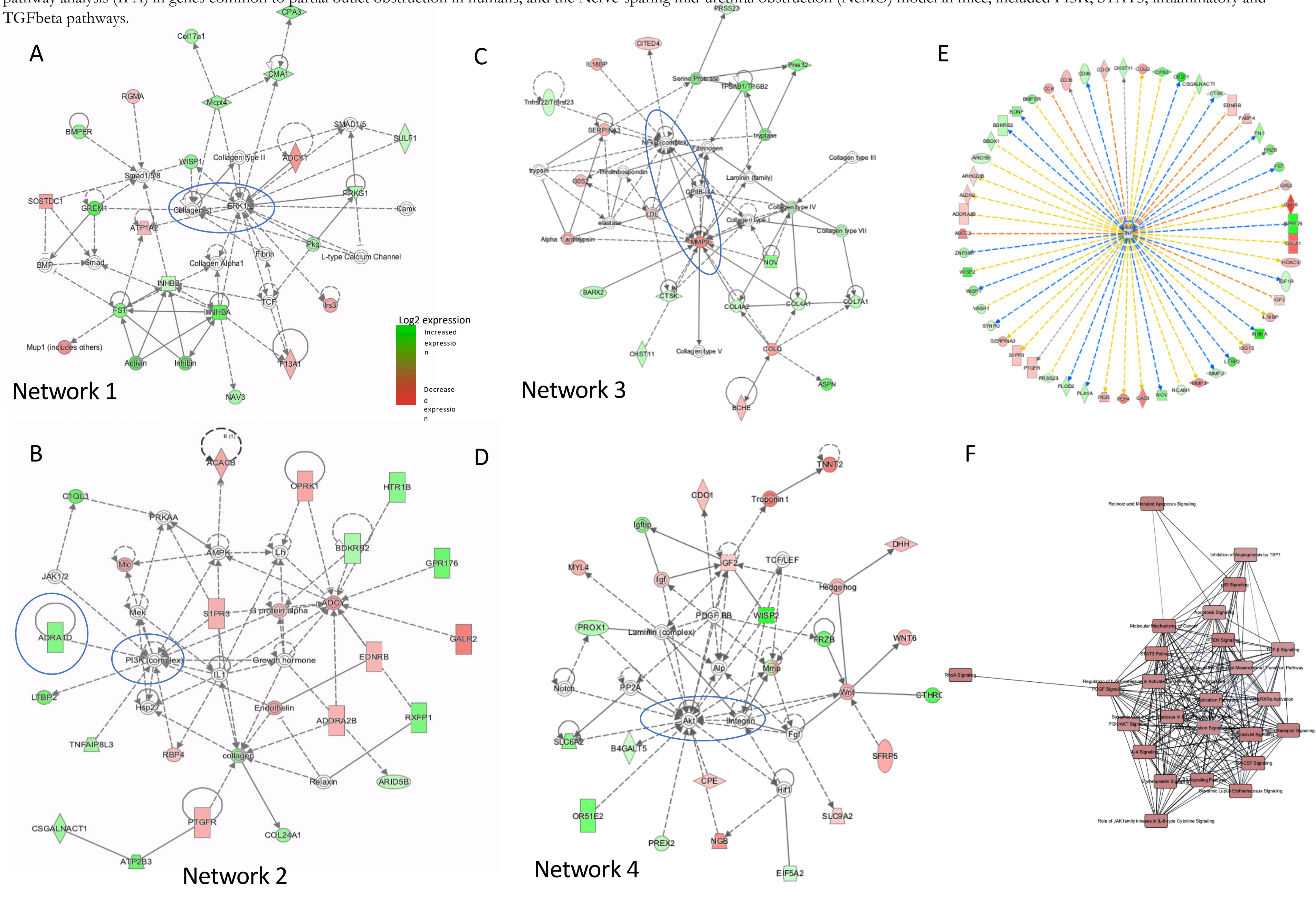

**Supplemental Figure S2:** Clodronate treatment effects on Bladder pathophysiology of NeMO vs. sham and on cell composition. (A) Mean and Maximum voided volumes, Total number of voids and Total voided volume were all significantly different from sham, by ANOVA with post-hoc t-test (*, p<0.05, **, p<0.01, ***, p<0.005). Body mass and change in body mass were not significantly altered. Bladder capacity was decreased by CL (Welch's t-test, p<0.05). *CL leads to macrophage depletion*: : (B-E) Mass cytometry time of flight analysis of pooled bladder and spleen samples of controls 2 weeks of NeMO. Of the 129,000 live cells counted from the spleen, ~12,500 were CD11b-positive macrophages. In control bladders only 3.5% of live cells were macrophages (B), while in NeMO bladders (D), 17% of live cells were macrophages. CL reduced the percentage of CD11b+ macrophages in both sham (C) and NeMO bladders to 7% of live cells (E). pooled n=3 for each group. The bladders of each group were pooled, minced and digested into a single cell suspension using collagenase. Single cell suspensions were quantitated on a CyTOF2® (cytometry by time of flight) mass cytometer (Fluidigm, San Francisco, CA, USA). The following antibodies were used: Rat anti-mouse CD45 to detect myeloid cells, rat anti-mouse CD3 to label T-cells, rat anti-mouse CD11b to label macrophages (all Fluidigm), rat anti-CD19 to label B-cells, and rat anti-mouse CD4 to label T-helper cells (Biolegend, San Diego, California, United States). Data analysis was performed on Cytobank (Cytobank, Santa Clara, CA, USA).

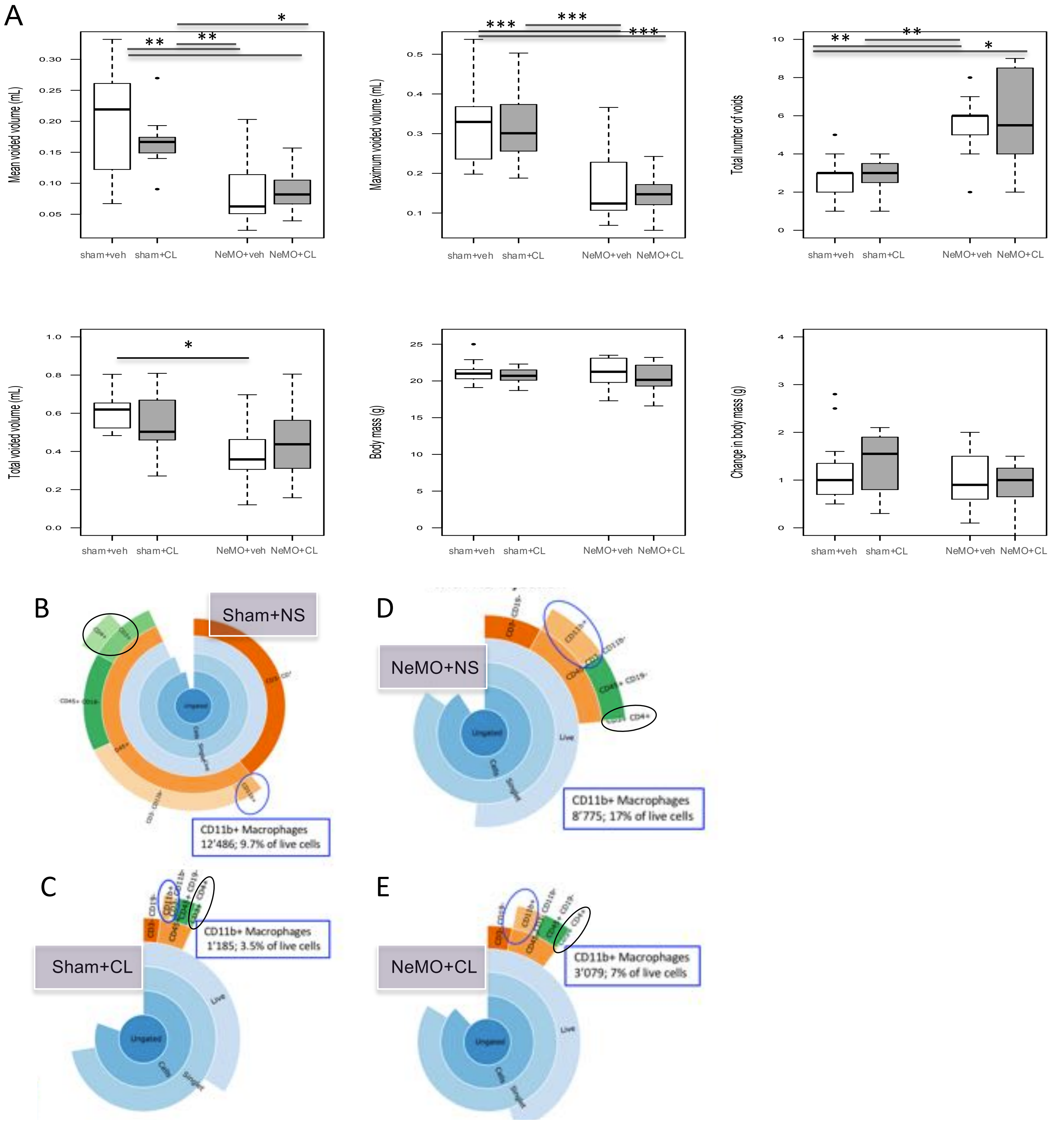

**Supplemental Figure S3**: Cytokine expression, altered by NeMO and partially restored by clodronate treatment (CL), correlated with bladder function during nerve-sparing mid-urethral obstruction (NeMO). Examining only factors altered during NeMO vs. sham or with clodronate treatment (A-F), we found discrete groups of in the Pearson's correlations (*R*-value) between secreted factors and bladder function in sham, NeMO +/- CL. The secreted factors appeared to correlate with together in 3 groups (I, II and III), by hierarchical clustering. Secreted factors correlated strongly with functional parameters affected by CL, including: small to total voids ratio (hyperactivity) – TNF; residual volumes - IL17 and CCL17; bladder capacity - TNF, CCL4, TREM1, IL16, IL17, CXCL9, sICAM1 and IFNG (negatively), CCL1 (positively) and IL7 and CCL2 (weakly positive).

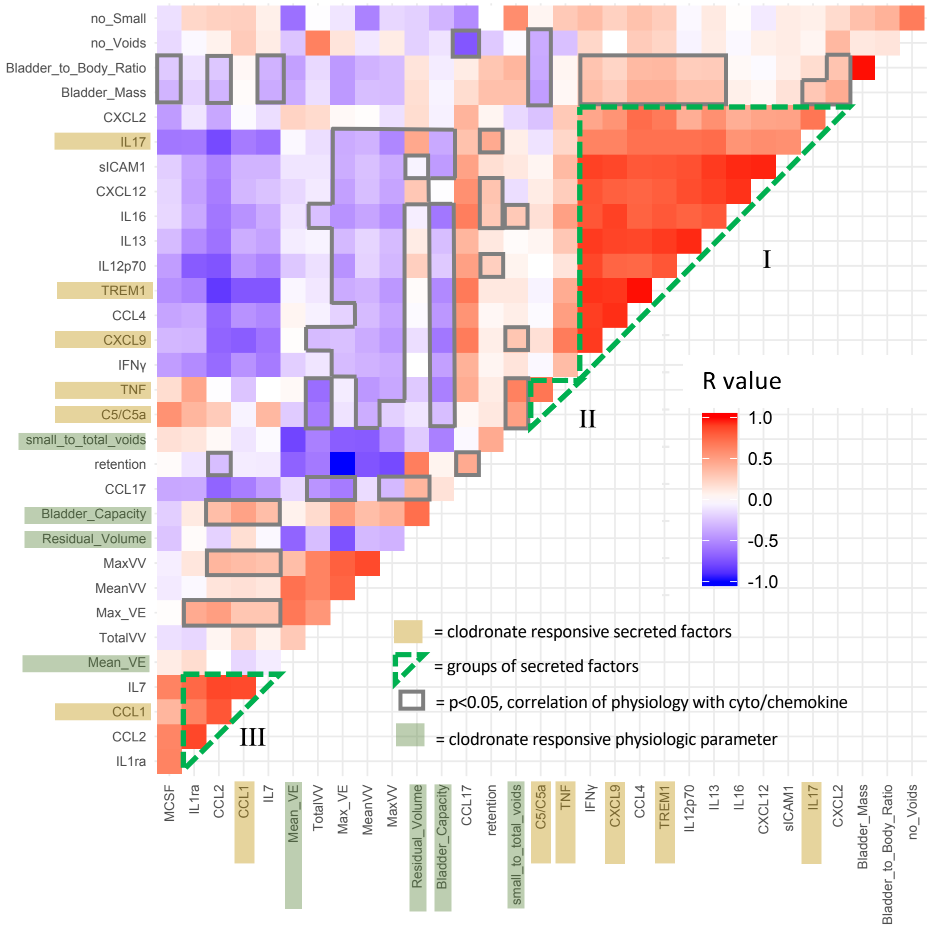

**Table S1**: RNAseq revealed 831 differentially expressed genes between sham (Group 0) and mouse NeMO (Group 2).

| Gene | baseMean | log2FoldChange | lfcSE | stat | pvalue | padj | sham | sham | sham | sham | NeMO | NeMO | NeMO | NeMO |
|---|---|---|---|---|---|---|---|---|---|---|---|---|---|---|
| Tnnt2 | 6126.19 | -2.00 | 0.19 | 112.38 | 2.94E-26 | 4.69E-22 | 7720.38 | 9175.45 | 15035.16 | 6684.90 | 3237.74 | 1764.40 | 2590.99 | 2800.52 |
| Atp2b3 | 149.12 | 3.21 | 0.31 | 101.15 | 8.53E-24 | 6.80E-20 | 43.14 | 34.47 | 20.63 | 31.08 | 182.71 | 199.58 | 232.16 | 449.23 |
| Cpe | 8274.88 | -0.88 | 0.10 | 71.99 | 2.17E-17 | 1.15E-13 | 9175.07 | 10803.32 | 11927.66 | 10843.31 | 5923.12 | 5707.37 | 5625.05 | 6194.18 |
| Mfap4 | 5184.24 | 1.47 | 0.18 | 65.72 | 5.21E-16 | 1.66E-12 | 3054.30 | 3638.77 | 2009.38 | 3194.14 | 4733.39 | 10974.35 | 7116.37 | 6753.18 |
| Itgbl1 | 713.26 | 1.85 | 0.23 | 65.97 | 4.58E-16 | 1.66E-12 | 480.10 | 219.28 | 305.78 | 270.26 | 1016.58 | 1241.55 | 872.82 | 1299.72 |
| Fam211b | 410.88 | 1.08 | 0.14 | 62.48 | 2.69E-15 | 7.14E-12 | 311.96 | 259.50 | 228.12 | 300.39 | 458.89 | 567.83 | 496.22 | 664.16 |
| Inhba | 47.74 | 3.52 | 0.46 | 60.00 | 9.49E-15 | 2.16E-11 | 5.53 | 8.62 | 7.28 | 10.36 | 55.24 | 127.43 | 101.02 | 66.42 |
| Crlf1 | 66.32 | 3.91 | 0.51 | 56.91 | 4.56E-14 | 9.08E-11 | 12.17 | 3.83 | 9.71 | 7.53 | 58.42 | 170.54 | 182.54 | 85.79 |
| Slc6a2 | 485.18 | 1.57 | 0.21 | 55.65 | 8.66E-14 | 1.53E-10 | 338.51 | 230.77 | 206.28 | 233.53 | 520.50 | 808.65 | 855.10 | 688.14 |
| Efemp1 | 4527.01 | -0.63 | 0.09 | 54.26 | 1.76E-13 | 2.81E-10 | 5878.51 | 5806.71 | 5055.00 | 5589.74 | 3171.88 | 3594.40 | 3050.00 | 4069.80 |
| Prss23 | 10031.96 | 0.70 | 0.10 | 48.98 | 2.59E-12 | 3.75E-09 | 8012.42 | 8613.36 | 7189.36 | 7438.24 | 10018.10 | 13427.46 | 12091.01 | 13465.76 |
| Mup10 | 151.51 | -1.79 | 0.26 | 46.68 | 8.36E-12 | 1.11E-08 | 190.27 | 172.36 | 387.07 | 126.18 | 158.28 | 48.72 | 89.50 | 39.66 |
| Htr1b | 151.52 | 1.93 | 0.29 | 42.10 | 8.65E-11 | 1.06E-07 | 69.69 | 55.54 | 59.46 | 67.80 | 184.83 | 320.46 | 301.28 | 153.12 |
| Cpa3 | 255.13 | 2.32 | 0.34 | 41.47 | 1.20E-10 | 1.36E-07 | 99.56 | 129.27 | 55.82 | 85.69 | 203.95 | 314.84 | 591.04 | 560.84 |
| Gria1 | 342.41 | -2.23 | 0.34 | 41.10 | 1.45E-10 | 1.54E-07 | 441.39 | 657.85 | 575.15 | 592.31 | 146.59 | 65.59 | 60.26 | 200.17 |
| Fam150a | 57.07 | 4.99 | 0.75 | 40.45 | 2.02E-10 | 2.01E-07 | 13.27 | 1.92 | 1.21 | 0.94 | 29.74 | 187.40 | 140.89 | 81.17 |
| Sfrp4 | 921.21 | 1.60 | 0.25 | 38.00 | 7.06E-10 | 6.25E-07 | 599.58 | 648.28 | 262.09 | 560.29 | 781.82 | 1977.11 | 1459.43 | 1081.10 |
| Apobec2 | 178.42 | -2.60 | 0.41 | 38.09 | 6.76E-10 | 6.25E-07 | 267.71 | 354.30 | 344.60 | 257.08 | 47.80 | 13.12 | 51.39 | 91.32 |
| Spock2 | 1647.32 | -1.18 | 0.19 | 37.85 | 7.65E-10 | 6.42E-07 | 2099.62 | 2566.29 | 2418.29 | 1921.95 | 1320.38 | 673.72 | 1002.19 | 1176.11 |
| Ngb | 1031.51 | -1.64 | 0.26 | 37.57 | 8.84E-10 | 6.76E-07 | 1160.44 | 2284.77 | 1323.81 | 1557.52 | 523.69 | 330.77 | 356.22 | 714.89 |
| Adora2b | 859.16 | -0.80 | 0.13 | 37.55 | 8.90E-10 | 6.76E-07 | 1016.63 | 1230.48 | 1141.80 | 914.36 | 722.33 | 573.45 | 688.51 | 585.75 |
| Mrgprb1 | 49.04 | 2.33 | 0.38 | 37.39 | 9.69E-10 | 7.02E-07 | 27.66 | 13.41 | 9.71 | 21.66 | 77.54 | 76.84 | 57.60 | 107.93 |
| Bean1 | 384.43 | 1.74 | 0.28 | 37.19 | 1.07E-09 | 7.44E-07 | 273.24 | 160.87 | 140.75 | 160.08 | 435.52 | 406.67 | 714.21 | 784.07 |
| Phldb2 | 6494.83 | 0.56 | 0.09 | 36.80 | 1.31E-09 | 8.69E-07 | 5919.44 | 5951.31 | 4347.59 | 5444.73 | 7020.42 | 8210.15 | 7274.99 | 7790.00 |
| Btbd7 | 987.84 | 0.72 | 0.12 | 36.39 | 1.61E-09 | 1.03E-06 | 775.47 | 878.09 | 700.13 | 676.12 | 1186.54 | 1202.19 | 1168.78 | 1315.40 |
| Cthrc1 | 119.28 | 2.43 | 0.39 | 36.31 | 1.68E-09 | 1.03E-06 | 37.61 | 49.79 | 33.98 | 32.02 | 157.21 | 333.58 | 221.53 | 88.55 |
| P4ha3 | 171.70 | 2.63 | 0.42 | 35.92 | 2.06E-09 | 1.22E-06 | 76.33 | 57.45 | 38.83 | 33.90 | 122.16 | 537.85 | 342.93 | 164.19 |
| Hhipl1 | 163.93 | 1.39 | 0.23 | 34.96 | 3.36E-09 | 1.91E-06 | 92.92 | 93.84 | 93.43 | 75.33 | 193.33 | 264.24 | 309.25 | 189.10 |
| Pilra | 434.19 | 1.37 | 0.23 | 34.74 | 3.77E-09 | 2.07E-06 | 311.96 | 325.57 | 199.00 | 193.04 | 411.09 | 549.09 | 550.28 | 933.51 |
| Ifi27l2a | 2287.05 | -0.79 | 0.14 | 34.67 | 3.92E-09 | 2.08E-06 | 2521.10 | 2595.98 | 3481.23 | 2875.86 | 1563.63 | 1458.94 | 1909.57 | 1890.08 |
| Hdac9 | 223.60 | -1.08 | 0.19 | 33.70 | 6.42E-09 | 3.30E-06 | 220.14 | 299.72 | 357.95 | 322.05 | 163.59 | 143.36 | 142.66 | 139.29 |
| Tacr2 | 333.28 | -1.48 | 0.25 | 33.22 | 8.23E-09 | 4.10E-06 | 432.54 | 557.31 | 446.53 | 538.63 | 195.45 | 104.95 | 142.66 | 248.14 |
| C1qtnf3 | 1087.45 | 1.15 | 0.20 | 32.87 | 9.83E-09 | 4.75E-06 | 600.68 | 718.18 | 467.16 | 1159.19 | 897.60 | 1832.81 | 1164.35 | 1859.64 |
| Gpr176 | 65.04 | 2.73 | 0.47 | 32.42 | 1.24E-08 | 5.82E-06 | 21.02 | 19.15 | 10.92 | 26.37 | 32.93 | 187.40 | 129.37 | 93.17 |
| Cyp2f2 | 5857.44 | 0.98 | 0.17 | 32.32 | 1.30E-08 | 5.94E-06 | 3282.18 | 4135.75 | 4358.51 | 3745.02 | 6958.81 | 9456.38 | 9235.07 | 5687.76 |
| Aldh2 | 10244.05 | -0.85 | 0.15 | 31.70 | 1.80E-08 | 7.97E-06 | 10070.01 | 12382.36 | 17511.69 | 12088.20 | 7470.82 | 6111.23 | 7804.00 | 8514.11 |
| Clip4 | 965.95 | 0.70 | 0.12 | 31.54 | 1.96E-08 | 8.43E-06 | 707.99 | 868.52 | 611.55 | 817.37 | 1180.16 | 1195.63 | 1155.49 | 1190.87 |
| Tnrc6a | 4226.80 | 0.52 | 0.09 | 31.10 | 2.45E-08 | 1.03E-05 | 3867.38 | 3686.65 | 3106.29 | 3378.71 | 4958.59 | 4876.24 | 4896.66 | 5043.90 |
| Atl3 | 7411.30 | 0.50 | 0.09 | 30.68 | 3.05E-08 | 1.24E-05 | 6378.52 | 6653.21 | 6022.07 | 5950.40 | 7496.31 | 9672.83 | 7846.53 | 9270.51 |
| Rgs4 | 730.04 | 1.51 | 0.28 | 30.30 | 3.71E-08 | 1.48E-05 | 341.82 | 339.94 | 441.68 | 404.92 | 523.69 | 1536.71 | 1307.90 | 943.66 |
| Tmem2 | 1026.01 | 0.75 | 0.14 | 30.14 | 4.02E-08 | 1.56E-05 | 740.07 | 795.74 | 734.10 | 807.95 | 1180.16 | 1495.48 | 1373.48 | 1081.10 |
| Acacb | 1276.50 | -1.02 | 0.19 | 30.08 | 4.14E-08 | 1.57E-05 | 1445.84 | 1250.59 | 2640.34 | 1301.39 | 810.50 | 608.12 | 1273.34 | 881.85 |
| Slc9a2 | 680.56 | -0.70 | 0.13 | 29.60 | 5.32E-08 | 1.97E-05 | 784.32 | 826.38 | 883.35 | 812.66 | 674.53 | 452.58 | 525.47 | 485.20 |
| Chst11 | 984.57 | 0.94 | 0.18 | 29.34 | 6.06E-08 | 2.19E-05 | 716.84 | 638.70 | 735.32 | 645.98 | 945.40 | 1756.91 | 1206.89 | 1230.53 |
| Wdfy2 | 143.90 | 1.07 | 0.20 | 29.26 | 6.34E-08 | 2.25E-05 | 99.56 | 100.55 | 93.43 | 78.16 | 195.45 | 218.33 | 192.29 | 173.42 |
| Lars2 | 4135.65 | -1.19 | 0.22 | 28.72 | 8.35E-08 | 2.77E-05 | 7246.91 | 3010.60 | 7091.07 | 5366.57 | 2515.41 | 2598.35 | 2942.78 | 2313.48 |
| Sult1a1 | 530.07 | -0.96 | 0.18 | 28.76 | 8.21E-08 | 2.77E-05 | 714.62 | 649.23 | 802.05 | 646.93 | 254.94 | 357.94 | 374.83 | 440.00 |
| Foxn3 | 334.17 | 0.89 | 0.16 | 28.75 | 8.24E-08 | 2.77E-05 | 272.13 | 256.63 | 220.84 | 195.87 | 473.76 | 453.52 | 384.57 | 416.02 |
| Kalrn | 1015.19 | 0.71 | 0.13 | 28.63 | 8.74E-08 | 2.84E-05 | 743.39 | 857.03 | 751.09 | 777.82 | 958.15 | 1355.86 | 1318.54 | 1359.68 |
| Gm26924 | 45640.83 | -1.05 | 0.20 | 28.55 | 9.11E-08 | 2.90E-05 | 49894.28 | 37339.54 | 86627.79 | 67835.92 | 26901.53 | 29284.61 | 39355.83 | 27887.18 |
| Galr2 | 211.29 | -1.97 | 0.36 | 28.38 | 9.99E-08 | 3.12E-05 | 219.03 | 490.28 | 309.42 | 340.88 | 64.80 | 40.29 | 77.09 | 148.51 |
| Bdnf | 234.37 | 1.66 | 0.31 | 27.89 | 1.28E-07 | 3.93E-05 | 162.62 | 83.31 | 123.77 | 60.27 | 295.31 | 271.74 | 522.81 | 355.14 |
| Tspan7 | 3362.82 | -0.52 | 0.10 | 27.74 | 1.39E-07 | 4.14E-05 | 3510.07 | 4125.22 | 4325.75 | 3705.47 | 3089.03 | 2539.32 | 2785.94 | 2821.74 |
| Prkg1 | 169.63 | 1.13 | 0.22 | 27.72 | 1.40E-07 | 4.14E-05 | 116.15 | 131.19 | 84.94 | 117.71 | 158.28 | 317.65 | 217.10 | 214.01 |
| Fat4 | 1526.26 | 0.65 | 0.12 | 27.65 | 1.45E-07 | 4.21E-05 | 1150.48 | 1270.70 | 1113.90 | 1363.54 | 1413.86 | 2272.27 | 1684.50 | 1940.81 |
| Dhrs7 | 1873.44 | -0.50 | 0.10 | 27.32 | 1.73E-07 | 4.92E-05 | 2033.25 | 2313.49 | 2245.99 | 2145.12 | 1525.39 | 1457.06 | 1667.66 | 1599.51 |
| Zfp365 | 118.00 | 1.25 | 0.24 | 27.18 | 1.85E-07 | 5.18E-05 | 109.52 | 77.56 | 40.04 | 81.93 | 128.53 | 178.03 | 135.58 | 192.79 |
| Actr3b | 265.67 | -1.24 | 0.24 | 26.84 | 2.21E-07 | 5.89E-05 | 268.81 | 532.41 | 359.16 | 353.13 | 137.03 | 161.17 | 112.54 | 201.09 |
| Rxfp1 | 148.88 | 2.44 | 0.45 | 26.84 | 2.22E-07 | 5.89E-05 | 34.29 | 114.91 | 29.12 | 32.96 | 156.15 | 456.33 | 221.53 | 145.75 |
| Adra1d | 58.31 | 2.20 | 0.41 | 26.85 | 2.20E-07 | 5.89E-05 | 23.23 | 20.11 | 16.99 | 23.54 | 48.86 | 94.64 | 153.30 | 85.79 |
| Filip1l | 4537.41 | 1.09 | 0.21 | 26.69 | 2.39E-07 | 6.24E-05 | 3709.19 | 2810.47 | 2721.64 | 2780.75 | 3113.46 | 7826.91 | 6496.98 | 6839.89 |
| Ccdc88a | 1035.18 | 0.50 | 0.10 | 26.56 | 2.55E-07 | 6.56E-05 | 941.40 | 884.80 | 759.58 | 965.21 | 1019.76 | 1359.61 | 972.07 | 1379.05 |
| Ptpn14 | 3837.52 | 0.51 | 0.10 | 26.35 | 2.85E-07 | 7.22E-05 | 3599.67 | 3020.18 | 2850.26 | 3309.96 | 4296.81 | 4496.75 | 4637.92 | 4488.59 |
| Lgi3 | 83.36 | 1.43 | 0.28 | 26.11 | 3.22E-07 | 8.03E-05 | 30.97 | 47.88 | 53.39 | 49.91 | 89.23 | 134.93 | 124.06 | 136.52 |
| Herc2 | 4745.37 | 0.45 | 0.09 | 26.01 | 3.40E-07 | 8.34E-05 | 4216.95 | 4227.68 | 3885.29 | 3698.87 | 5397.30 | 5459.06 | 5731.38 | 5346.46 |
| Itih5 | 14815.15 | 0.75 | 0.15 | 25.93 | 3.55E-07 | 8.52E-05 | 12460.57 | 12173.61 | 9599.15 | 11741.67 | 14541.17 | 15879.64 | 16551.71 | 25573.70 |
| Nav1 | 5538.04 | 0.54 | 0.11 | 25.91 | 3.58E-07 | 8.52E-05 | 4792.19 | 4852.97 | 4011.48 | 4903.27 | 5080.75 | 6914.25 | 6648.51 | 7100.94 |
| Ldlrad4 | 467.33 | 0.90 | 0.18 | 25.73 | 3.93E-07 | 9.21E-05 | 346.25 | 402.18 | 332.47 | 251.43 | 495.01 | 725.25 | 545.85 | 640.17 |
| Utrn | 5924.11 | 0.39 | 0.08 | 25.53 | 4.36E-07 | 0.0001008 | 5221.40 | 5160.35 | 5129.01 | 5149.04 | 6215.24 | 6710.92 | 6618.38 | 7188.57 |
| Dnajc15 | 519.85 | -0.59 | 0.12 | 25.41 | 4.63E-07 | 0.0001053 | 592.94 | 579.33 | 701.34 | 552.76 | 505.63 | 344.82 | 492.68 | 389.27 |
| Mgl2 | 761.22 | -1.05 | 0.21 | 25.30 | 4.90E-07 | 0.0001086 | 765.51 | 1160.58 | 953.73 | 1300.44 | 284.68 | 657.79 | 373.94 | 593.13 |
| Coro6 | 666.02 | 0.98 | 0.19 | 25.30 | 4.91E-07 | 0.0001086 | 533.20 | 527.62 | 308.20 | 515.09 | 795.63 | 685.90 | 888.77 | 1073.72 |
| Tpsb2 | 195.36 | 2.10 | 0.40 | 25.15 | 5.31E-07 | 0.000116 | 76.33 | 86.18 | 58.24 | 82.87 | 100.91 | 290.48 | 543.19 | 324.70 |
| Cox6c | 5050.65 | -0.80 | 0.16 | 25.06 | 5.55E-07 | 0.0001196 | 6210.37 | 6692.47 | 7006.13 | 5478.63 | 4348.86 | 2739.84 | 3518.76 | 4410.18 |
| Akap5 | 461.47 | 0.87 | 0.17 | 24.97 | 5.83E-07 | 0.0001239 | 347.36 | 428.99 | 213.56 | 416.22 | 533.25 | 713.07 | 478.50 | 560.84 |
| Timd4 | 95.19 | -2.71 | 0.52 | 24.85 | 6.18E-07 | 0.0001297 | 128.32 | 158.00 | 99.50 | 287.21 | 6.37 | 7.50 | 24.81 | 49.81 |
| Prex2 | 1921.02 | 1.02 | 0.21 | 24.57 | 7.17E-07 | 0.0001484 | 1281.01 | 1562.76 | 1260.72 | 1119.64 | 1649.68 | 3912.05 | 2303.89 | 2278.42 |
| Bmper | 161.78 | 2.02 | 0.40 | 24.48 | 7.52E-07 | 0.0001537 | 57.52 | 80.44 | 44.90 | 97.93 | 111.54 | 442.27 | 309.25 | 150.36 |
| Arhgdib | 1802.18 | -0.56 | 0.11 | 24.42 | 7.75E-07 | 0.0001559 | 2287.68 | 2111.45 | 2032.43 | 2270.36 | 1141.92 | 1545.14 | 1506.39 | 1522.02 |
| Kcnma1 | 1220.34 | 0.56 | 0.11 | 24.40 | 7.82E-07 | 0.0001559 | 918.17 | 1181.64 | 882.14 | 1035.84 | 1349.06 | 1565.76 | 1371.70 | 1458.38 |
| Rgs6 | 206.95 | -0.98 | 0.20 | 24.18 | 8.76E-07 | 0.0001724 | 269.92 | 333.23 | 252.39 | 261.78 | 113.66 | 126.50 | 108.99 | 189.10 |
| Dlgap2 | 158.02 | 1.40 | 0.29 | 24.12 | 9.03E-07 | 0.0001755 | 126.11 | 114.91 | 67.95 | 64.98 | 114.72 | 254.87 | 216.21 | 304.40 |
| Csgalnact1 | 306.33 | 1.38 | 0.28 | 24.05 | 9.36E-07 | 0.0001798 | 144.92 | 182.90 | 158.95 | 225.06 | 228.38 | 738.37 | 404.95 | 367.13 |
| Kitl | 2106.99 | 0.45 | 0.09 | 23.85 | 1.04E-06 | 0.0001976 | 1901.61 | 1929.51 | 1559.21 | 1983.15 | 2133.00 | 2792.31 | 1859.95 | 2697.21 |
| Fabp4 | 14837.09 | -1.07 | 0.22 | 23.67 | 1.15E-06 | 0.0002148 | 20604.64 | 13302.58 | 28581.48 | 16446.24 | 8578.75 | 6484.16 | 12744.08 | 11954.81 |
| Cma1 | 190.73 | 1.78 | 0.36 | 23.62 | 1.17E-06 | 0.0002176 | 101.77 | 78.52 | 76.44 | 95.11 | 117.91 | 246.44 | 456.35 | 353.29 |
| Upk2 | 16303.71 | -0.42 | 0.09 | 23.48 | 1.26E-06 | 0.0002311 | 17753.88 | 16140.82 | 21546.22 | 16401.04 | 17874.51 | 12584.15 | 16466.64 | 11662.40 |
| Ppa1 | 2455.92 | -0.47 | 0.10 | 23.25 | 1.42E-06 | 0.0002552 | 2715.79 | 2441.81 | 3379.30 | 2714.83 | 2226.48 | 1903.08 | 2165.66 | 2100.39 |
| Arid5b | 1257.25 | 0.65 | 0.13 | 23.25 | 1.42E-06 | 0.0002552 | 1004.46 | 895.33 | 1087.20 | 874.81 | 1335.25 | 1533.90 | 1871.47 | 1455.61 |
| Clstn3 | 79.23 | -1.53 | 0.31 | 23.19 | 1.47E-06 | 0.0002605 | 101.77 | 70.86 | 183.22 | 83.81 | 49.93 | 19.68 | 88.61 | 35.98 |

| | | | | | | | | | | | | | | |
|---|---|---|---|---|---|---|---|---|---|---|---|---|---|---|
| Man1a2 | 3961.08 | 0.46 | 0.10 | 23.05 | 1.58E-06 | 0.0002745 | 3703.66 | 3453.96 | 3101.43 | 3325.03 | 4343.55 | 5215.44 | 4161.19 | 4384.35 |
| 0610009B22R | 482.06 | -0.59 | 0.12 | 23.04 | 1.58E-06 | 0.0002745 | 560.86 | 552.52 | 633.39 | 537.69 | 459.96 | 372.00 | 375.71 | 364.36 |
| Dbi | 3158.70 | -0.59 | 0.12 | 22.88 | 1.72E-06 | 0.0002883 | 3471.35 | 3385.97 | 4980.98 | 2970.96 | 2518.60 | 2310.69 | 3229.00 | 2402.03 |
| Bche | 2131.06 | -0.92 | 0.19 | 22.87 | 1.73E-06 | 0.0002883 | 2185.91 | 4118.51 | 1892.89 | 3362.70 | 1273.64 | 1543.27 | 1075.74 | 1595.82 |
| Col17a1 | 1906.88 | 1.17 | 0.24 | 22.88 | 1.72E-06 | 0.0002883 | 1319.73 | 1239.10 | 1099.33 | 1039.60 | 2608.89 | 3752.76 | 2781.51 | 1414.10 |
| Mgat5 | 918.49 | 0.65 | 0.14 | 22.87 | 1.74E-06 | 0.0002883 | 849.58 | 722.01 | 706.19 | 611.14 | 999.58 | 1209.69 | 1150.17 | 1099.55 |
| C530008M17I | 132.60 | -1.04 | 0.22 | 22.66 | 1.94E-06 | 0.0003181 | 150.45 | 197.26 | 166.23 | 210.93 | 74.36 | 86.21 | 65.57 | 109.77 |
| Adhfe1 | 381.55 | -0.80 | 0.17 | 22.27 | 2.36E-06 | 0.0003843 | 360.63 | 392.60 | 600.63 | 508.50 | 327.17 | 212.70 | 386.35 | 263.82 |
| C1ql3 | 34.62 | 3.31 | 0.67 | 22.25 | 2.39E-06 | 0.0003845 | 16.59 | 7.66 | 2.43 | 3.77 | 31.87 | 115.25 | 73.55 | 25.83 |
| Ctsk | 2407.46 | 0.82 | 0.17 | 22.09 | 2.61E-06 | 0.0004113 | 1643.86 | 2013.77 | 1355.36 | 2330.63 | 2629.07 | 4260.62 | 2451.88 | 2574.53 |
| Lyz1 | 146.31 | -1.84 | 0.39 | 22.10 | 2.59E-06 | 0.0004113 | 134.96 | 222.16 | 275.44 | 284.38 | 31.87 | 38.42 | 75.32 | 107.93 |
| Dnaic1 | 499.86 | -1.41 | 0.30 | 22.05 | 2.65E-06 | 0.0004143 | 661.53 | 880.01 | 706.19 | 691.18 | 230.51 | 149.92 | 186.97 | 492.58 |
| Rims4 | 21.40 | -4.15 | 0.87 | 21.97 | 2.77E-06 | 0.000428 | 32.08 | 10.53 | 94.64 | 7.53 | 9.56 | 0.00 | 16.84 | 0.00 |
| Cit | 802.33 | 1.21 | 0.25 | 21.94 | 2.82E-06 | 0.0004321 | 724.58 | 391.65 | 428.33 | 417.16 | 918.85 | 760.86 | 1310.56 | 1466.68 |
| Fat1 | 8547.13 | 0.46 | 0.10 | 21.88 | 2.90E-06 | 0.0004401 | 7558.87 | 7316.80 | 6974.58 | 7034.26 | 9030.20 | 10303.44 | 10870.84 | 9288.04 |
| Bbox1 | 455.11 | 1.03 | 0.22 | 21.85 | 2.95E-06 | 0.0004401 | 313.06 | 300.68 | 279.08 | 285.33 | 812.62 | 716.82 | 522.81 | 410.49 |
| Ppp1r1a | 361.67 | -1.20 | 0.25 | 21.85 | 2.94E-06 | 0.0004401 | 378.33 | 513.26 | 582.43 | 510.38 | 237.94 | 117.13 | 260.52 | 293.34 |
| Fbxl20 | 3358.97 | 0.40 | 0.08 | 21.81 | 3.01E-06 | 0.0004436 | 3046.56 | 2950.28 | 2640.34 | 3121.63 | 3717.88 | 4091.96 | 3618.89 | 3684.22 |
| Tecr | 4316.90 | -0.39 | 0.08 | 21.79 | 3.04E-06 | 0.0004453 | 4636.21 | 4392.38 | 5787.89 | 4089.67 | 4550.69 | 3213.97 | 4462.47 | 3401.96 |
| Gatad2b | 642.70 | 0.53 | 0.11 | 21.71 | 3.17E-06 | 0.0004587 | 552.01 | 594.65 | 464.73 | 521.68 | 715.96 | 774.91 | 759.40 | 758.24 |
| Igsf9b | 179.65 | 1.64 | 0.34 | 21.63 | 3.30E-06 | 0.0004741 | 177.00 | 86.18 | 49.75 | 77.22 | 155.09 | 334.52 | 307.48 | 249.98 |
| Ppfibp1 | 5398.62 | 0.42 | 0.09 | 21.54 | 3.46E-06 | 0.0004877 | 4949.27 | 4716.04 | 4220.18 | 4805.33 | 5600.19 | 6716.54 | 6286.09 | 5895.31 |
| Itgav | 3857.61 | 0.41 | 0.09 | 21.56 | 3.44E-06 | 0.0004877 | 3640.60 | 3513.33 | 3085.66 | 3331.62 | 3764.62 | 4778.79 | 3675.60 | 5070.65 |
| Adamts6 | 469.59 | 0.92 | 0.20 | 21.46 | 3.60E-06 | 0.0005039 | 399.35 | 341.85 | 293.64 | 310.75 | 467.39 | 885.48 | 556.48 | 501.81 |
| B3galnt2 | 2141.96 | 0.37 | 0.08 | 21.41 | 3.71E-06 | 0.000514 | 1981.26 | 2004.20 | 1720.59 | 1913.47 | 2182.93 | 2473.73 | 2271.11 | 2588.36 |
| Tulp4 | 1848.63 | 0.51 | 0.11 | 21.26 | 4.00E-06 | 0.0005501 | 1743.42 | 1506.26 | 1509.46 | 1343.76 | 2165.93 | 2279.76 | 2160.34 | 2080.10 |
| Igf2bp1 | 45.20 | -1.94 | 0.42 | 21.23 | 4.07E-06 | 0.0005544 | 63.06 | 69.90 | 88.58 | 64.98 | 12.75 | 31.86 | 21.27 | 9.22 |
| Serpina3b | 28.14 | -2.60 | 0.55 | 21.07 | 4.44E-06 | 0.0005996 | 23.23 | 42.13 | 72.80 | 38.61 | 27.62 | 1.87 | 13.29 | 5.53 |
| Fam102b | 2545.25 | 0.55 | 0.12 | 20.95 | 4.70E-06 | 0.0006299 | 1902.72 | 2263.70 | 2094.32 | 2155.48 | 2441.05 | 3154.94 | 2797.46 | 3552.31 |
| G0s2 | 620.45 | -1.02 | 0.23 | 20.85 | 4.97E-06 | 0.0006604 | 596.26 | 592.74 | 1488.83 | 547.11 | 357.98 | 309.22 | 684.08 | 387.42 |
| Cd109 | 539.09 | 0.71 | 0.16 | 20.77 | 5.18E-06 | 0.0006828 | 479.00 | 507.51 | 354.31 | 368.19 | 535.37 | 837.69 | 448.37 | 782.23 |
| Plod2 | 2336.45 | 1.07 | 0.24 | 20.69 | 5.41E-06 | 0.000706 | 1545.40 | 2173.69 | 1212.18 | 1439.81 | 1821.76 | 5784.21 | 2093.89 | 2620.65 |
| Trim46 | 267.97 | 1.06 | 0.23 | 20.66 | 5.49E-06 | 0.000706 | 194.70 | 159.91 | 179.58 | 152.55 | 263.44 | 406.67 | 493.57 | 293.34 |
| Rab3b | 45.17 | 1.83 | 0.40 | 20.66 | 5.49E-06 | 0.000706 | 17.70 | 14.36 | 15.77 | 34.84 | 58.42 | 58.10 | 72.66 | 89.48 |
| Krt20 | 3932.86 | -0.74 | 0.16 | 20.63 | 5.57E-06 | 0.00071 | 5820.98 | 4119.47 | 5611.94 | 3528.43 | 3879.34 | 2380.96 | 3364.57 | 2757.17 |
| Mir143hg | 1250.81 | 0.81 | 0.17 | 20.53 | 5.88E-06 | 0.0007436 | 1098.49 | 1005.45 | 580.00 | 1226.99 | 1345.87 | 1366.17 | 1439.93 | 1943.58 |
| Upk1a | 48977.64 | -0.47 | 0.10 | 20.48 | 6.04E-06 | 0.0007533 | 56637.86 | 48445.45 | 66857.97 | 46438.38 | 53982.58 | 33031.74 | 49431.82 | 36995.34 |
| Apoc1 | 604.05 | -0.88 | 0.19 | 20.47 | 6.05E-06 | 0.0007533 | 750.02 | 776.59 | 843.31 | 714.73 | 488.64 | 270.80 | 462.55 | 525.79 |
| Atp1a3 | 185.03 | -1.52 | 0.34 | 20.42 | 6.23E-06 | 0.0007702 | 136.07 | 147.47 | 611.55 | 133.72 | 78.61 | 44.98 | 260.52 | 67.34 |
| Maob | 952.14 | -0.90 | 0.20 | 20.39 | 6.32E-06 | 0.0007754 | 943.61 | 1723.63 | 1153.94 | 1194.04 | 660.72 | 595.94 | 532.55 | 812.67 |
| Dock7 | 4823.70 | 0.36 | 0.08 | 20.36 | 6.41E-06 | 0.000776 | 4409.43 | 4409.62 | 3969.01 | 4264.82 | 5274.08 | 5786.08 | 5277.69 | 5198.87 |
| Sntb2 | 723.81 | 0.72 | 0.16 | 20.36 | 6.43E-06 | 0.000776 | 715.73 | 602.31 | 453.81 | 489.67 | 706.40 | 954.82 | 926.87 | 940.89 |
| Crim1 | 3217.34 | 0.37 | 0.08 | 20.32 | 6.54E-06 | 0.0007784 | 3073.11 | 2817.18 | 2567.54 | 3014.28 | 3119.83 | 3862.39 | 3400.90 | 3883.47 |
| Zfp568 | 771.08 | 0.52 | 0.12 | 20.33 | 6.51E-06 | 0.0007784 | 690.29 | 586.03 | 595.78 | 669.53 | 856.17 | 932.33 | 965.86 | 872.63 |
| Tgfb2 | 1225.42 | 0.63 | 0.14 | 20.29 | 6.66E-06 | 0.0007869 | 1073.04 | 854.15 | 982.85 | 987.81 | 1138.73 | 1677.26 | 1523.23 | 1566.30 |
| Dock5 | 1970.41 | 0.40 | 0.09 | 20.25 | 6.81E-06 | 0.0007953 | 1851.83 | 1720.76 | 1641.72 | 1627.20 | 2209.48 | 2202.93 | 2193.13 | 2316.24 |
| Zbed6 | 314.99 | 1.00 | 0.22 | 20.24 | 6.84E-06 | 0.0007953 | 279.88 | 220.24 | 167.45 | 194.93 | 312.30 | 432.90 | 520.15 | 392.04 |
| Car5b | 981.45 | -0.76 | 0.17 | 20.08 | 7.41E-06 | 0.0008561 | 1056.45 | 925.97 | 1724.23 | 1106.46 | 672.41 | 595.01 | 1013.71 | 757.32 |
| Enpp1 | 2785.32 | 0.37 | 0.08 | 20.02 | 7.66E-06 | 0.0008786 | 2406.05 | 2226.35 | 2662.18 | 2324.98 | 3201.63 | 3151.19 | 3201.53 | 3108.62 |
| Man2a1 | 3513.07 | 0.51 | 0.12 | 19.98 | 7.83E-06 | 0.0008915 | 2896.11 | 3070.93 | 2743.48 | 3023.70 | 3498.00 | 4810.65 | 4263.98 | 3797.68 |
| Slc12a8 | 391.10 | -0.64 | 0.14 | 19.90 | 8.17E-06 | 0.0009238 | 403.77 | 447.19 | 531.47 | 458.59 | 406.84 | 237.07 | 358.88 | 285.03 |
| Stard9 | 1288.96 | 0.63 | 0.14 | 19.83 | 8.47E-06 | 0.0009505 | 1188.09 | 1237.18 | 802.05 | 1005.70 | 1234.34 | 1891.84 | 1491.33 | 1461.14 |
| Mcpt4 | 180.53 | 1.70 | 0.37 | 19.79 | 8.66E-06 | 0.0009657 | 86.29 | 81.39 | 70.38 | 116.77 | 126.41 | 165.85 | 379.26 | 417.86 |
| Lrp6 | 4561.19 | 0.42 | 0.10 | 19.71 | 9.01E-06 | 0.0009972 | 3950.35 | 4152.03 | 3893.78 | 3854.25 | 4474.20 | 5930.38 | 4769.06 | 5465.45 |
| 1110001J03Ri | 427.28 | -0.70 | 0.16 | 19.67 | 9.20E-06 | 0.0010045 | 495.59 | 533.37 | 639.46 | 403.98 | 355.85 | 257.68 | 384.57 | 347.76 |
| Tnnc2 | 125.27 | -1.44 | 0.31 | 19.68 | 9.15E-06 | 0.0010045 | 134.96 | 226.94 | 189.29 | 163.85 | 54.17 | 44.98 | 117.85 | 70.11 |
| Slc30a4 | 1365.00 | 0.39 | 0.09 | 19.56 | 9.73E-06 | 0.0010547 | 1241.19 | 1277.40 | 1047.16 | 1290.09 | 1380.93 | 1726.92 | 1374.36 | 1581.98 |
| Cep170 | 1083.09 | 0.48 | 0.11 | 19.51 | 1.00E-05 | 0.0010785 | 949.15 | 908.74 | 867.58 | 986.87 | 1055.88 | 1471.12 | 1027.01 | 1398.42 |
| Gstm1 | 158497.27 | -0.52 | 0.12 | 19.48 | 1.02E-05 | 0.0010867 | 188686.29 | 170112.57 | 210476.42 | 161808.79 | 149317.55 | 101090.13 | 148083.69 | 138402.75 |
| Akap13 | 5511.15 | 0.31 | 0.07 | 19.43 | 1.04E-05 | 0.0011064 | 5135.12 | 5074.17 | 4603.61 | 5089.72 | 5670.30 | 6180.57 | 5964.43 | 6371.29 |
| Gxylt2 | 174.81 | 1.24 | 0.27 | 19.36 | 1.08E-05 | 0.0011403 | 168.15 | 127.36 | 70.38 | 80.04 | 208.20 | 309.22 | 230.39 | 204.78 |
| Tnfrsf22 | 277.26 | 0.72 | 0.16 | 19.28 | 1.13E-05 | 0.0011807 | 200.23 | 221.20 | 184.44 | 252.37 | 304.87 | 399.17 | 338.50 | 317.32 |
| Adam12 | 474.26 | 1.34 | 0.31 | 19.16 | 1.20E-05 | 0.0012503 | 265.50 | 356.22 | 234.18 | 271.20 | 335.67 | 1349.31 | 526.35 | 455.68 |
| Ccdc136 | 1164.14 | 0.65 | 0.15 | 19.13 | 1.22E-05 | 0.0012655 | 1095.17 | 976.72 | 787.49 | 902.12 | 1166.35 | 1265.91 | 1223.72 | 1895.61 |
| Gm12866 | 23.71 | 2.28 | 0.53 | 19.00 | 1.31E-05 | 0.0013439 | 8.85 | 5.75 | 12.13 | 1.88 | 52.05 | 34.67 | 53.17 | 21.22 |
| Tmem261 | 587.17 | -0.58 | 0.13 | 18.95 | 1.35E-05 | 0.0013744 | 710.20 | 719.14 | 787.49 | 585.72 | 432.34 | 475.07 | 506.86 | 480.59 |
| Psca | 20232.39 | -0.56 | 0.13 | 18.91 | 1.37E-05 | 0.0013882 | 26397.96 | 19388.91 | 27456.66 | 19420.03 | 21034.71 | 14317.63 | 20097.93 | 13745.26 |
| Postn | 20481.03 | 0.64 | 0.15 | 18.88 | 1.39E-05 | 0.0013963 | 14951.80 | 20244.02 | 14457.58 | 16585.61 | 18010.48 | 32190.30 | 23162.11 | 24246.31 |
| Igfbp5 | 11323.43 | -0.55 | 0.12 | 18.88 | 1.39E-05 | 0.0013963 | 13531.40 | 13776.58 | 12271.05 | 14117.49 | 10936.94 | 7808.17 | 8651.12 | 9494.66 |
| Adig | 263.23 | -1.29 | 0.29 | 18.74 | 1.50E-05 | 0.0014939 | 240.05 | 244.18 | 618.83 | 277.79 | 236.88 | 46.85 | 284.44 | 156.81 |
| Sema3c | 7915.79 | 0.48 | 0.11 | 18.73 | 1.51E-05 | 0.001494 | 7464.84 | 7500.66 | 5786.67 | 6426.89 | 8112.42 | 8615.88 | 8533.27 | 10885.70 |
| Robo1 | 1206.89 | 0.41 | 0.09 | 18.69 | 1.54E-05 | 0.0015054 | 1163.75 | 1133.76 | 882.14 | 1103.64 | 1224.78 | 1577.94 | 1167.01 | 1402.11 |
| Eif5a2 | 309.38 | 0.69 | 0.16 | 18.70 | 1.53E-05 | 0.0015054 | 250.01 | 245.14 | 203.85 | 285.33 | 363.29 | 416.04 | 264.95 | 446.46 |
| Ccar1 | 2661.48 | 0.34 | 0.08 | 18.63 | 1.58E-05 | 0.0015348 | 2459.15 | 2622.79 | 2043.35 | 2541.56 | 2621.64 | 3097.78 | 2629.98 | 3275.58 |
| Sertad4 | 743.15 | 0.79 | 0.18 | 18.63 | 1.59E-05 | 0.0015348 | 658.21 | 485.49 | 509.63 | 576.30 | 721.27 | 1270.60 | 931.30 | 792.38 |
| mmu-mir-623( | 2454.40 | -1.75 | 0.40 | 18.56 | 1.64E-05 | 0.001577 | 5240.21 | 1560.84 | 4748.01 | 3390.95 | 1115.36 | 1112.24 | 1461.20 | 1006.38 |
| Heatr5a | 2296.79 | 0.47 | 0.11 | 18.53 | 1.67E-05 | 0.0015944 | 2014.44 | 2003.24 | 1822.52 | 1914.41 | 2404.94 | 2890.70 | 2886.96 | 2437.08 |
| Dgat2 | 6603.05 | -0.79 | 0.19 | 18.49 | 1.71E-05 | 0.0016194 | 6212.59 | 5293.45 | 15433.15 | 5036.04 | 5124.30 | 3381.70 | 8122.11 | 4221.08 |
| Frem2 | 1880.85 | 0.52 | 0.12 | 18.48 | 1.72E-05 | 0.0016224 | 1488.99 | 1911.31 | 1448.79 | 1484.07 | 1848.32 | 2576.80 | 1910.46 | 2378.05 |
| Ccrl2 | 224.74 | -0.76 | 0.17 | 18.34 | 1.85E-05 | 0.0017252 | 246.69 | 304.51 | 254.81 | 325.82 | 176.33 | 148.05 | 169.25 | 172.50 |
| Slitrk3 | 33.84 | -2.32 | 0.52 | 18.34 | 1.84E-05 | 0.0017252 | 21.02 | 91.93 | 46.11 | 67.80 | 12.75 | 9.37 | 8.86 | 12.91 |
| Cxcl12 | 10650.00 | -0.56 | 0.13 | 18.31 | 1.88E-05 | 0.0017283 | 10473.78 | 15154.53 | 11511.46 | 14145.74 | 8711.53 | 8818.27 | 7821.72 | 8563.00 |
| Diap2 | 1671.03 | 0.47 | 0.11 | 18.31 | 1.87E-05 | 0.0017283 | 1544.30 | 1636.49 | 1191.55 | 1400.26 | 1805.83 | 1872.16 | 1687.16 | 2230.46 |
| Pigr | 645.99 | -1.11 | 0.25 | 18.30 | 1.89E-05 | 0.0017309 | 568.60 | 845.54 | 730.46 | 1420.04 | 407.90 | 452.58 | 399.64 | 343.15 |
| 4732418A04R | 35.20 | -2.28 | 0.53 | 18.12 | 2.07E-05 | 0.0018863 | 46.46 | 95.76 | 47.32 | 47.08 | 7.44 | 4.69 | 8.86 | 23.98 |
| Pkd2 | 5164.92 | 0.35 | 0.08 | 18.02 | 2.18E-05 | 0.0019785 | 4923.83 | 5258.02 | 3761.52 | 4806.28 | 5102.00 | 6277.08 | 5185.53 | 6005.08 |
| Gm2a | 4215.02 | -0.44 | 0.11 | 17.58 | 2.75E-05 | 0.0024797 | 4314.30 | 4941.07 | 4875.41 | 5409.89 | 3180.38 | 3808.04 | 3516.10 | 3675.00 |
| Map4k5 | 3179.94 | 0.33 | 0.08 | 17.56 | 2.78E-05 | 0.0024889 | 2917.13 | 3069.02 | 2556.62 | 2969.08 | 3365.21 | 3766.81 | 3148.36 | 3647.32 |
| Ky | 25.52 | -1.80 | 0.43 | 17.55 | 2.80E-05 | 0.0024915 | 38.72 | 41.18 | 41.26 | 35.78 | 11.68 | 13.12 | 15.06 | 7.38 |
| Cst3 | 9642.18 | -0.36 | 0.09 | 17.46 | 2.93E-05 | 0.0025829 | 10796.80 | 10563.93 | 11091.63 | 10845.20 | 7751.25 | 8272.93 | 8984.30 | 8831.43 |
| Gria3 | 715.18 | 0.81 | 0.19 | 17.46 | 2.93E-05 | 0.0025829 | 532.10 | 643.49 | 396.78 | 631.86 | 598.05 | 1307.14 | 842.69 | 769.31 |
| Mybpc2 | 43.31 | -1.67 | 0.40 | 17.42 | 3.00E-05 | 0.0026282 | 30.97 | 67.99 | 101.93 | 53.68 | 21.25 | 13.12 | 36.33 | 21.22 |
| Inhbb | 464.65 | 0.62 | 0.15 | 17.40 | 3.03E-05 | 0.0026353 | 377.22 | 402.18 | 383.43 | 311.69 | 564.06 | 646.54 | 498.88 | 533.17 |

| | | | | | | | | | | | | | |
|---|---|---|---|---|---|---|---|---|---|---|---|---|---|
| Fn1 | 87598.58 | 1.45 | 0.35 | 17.34 | 3.12E-05 | 0.0027025 | 50118.85 | 42261.46 | 51361.72 | 45878.09 | 68853.04 | 139060.89 | 153033.51 | 150221.04 |
| Ahnak | 41320.62 | 0.37 | 0.09 | 17.32 | 3.16E-05 | 0.0027244 | 40127.38 | 37891.10 | 34030.83 | 34162.79 | 43863.56 | 45792.98 | 43503.73 | 51192.60 |
| Dkkl1 | 124.92 | -0.81 | 0.20 | 17.29 | 3.20E-05 | 0.0027294 | 149.34 | 168.53 | 157.74 | 164.79 | 75.42 | 92.76 | 94.81 | 95.93 |
| Brinp2 | 122.88 | -0.92 | 0.22 | 17.30 | 3.19E-05 | 0.0027294 | 168.15 | 151.30 | 175.94 | 152.55 | 67.98 | 110.57 | 74.43 | 82.10 |
| S1pr3 | 311.60 | -0.81 | 0.20 | 17.28 | 3.23E-05 | 0.0027361 | 304.21 | 414.63 | 498.70 | 370.08 | 192.27 | 242.69 | 208.24 | 261.97 |
| Alg10b | 803.40 | 0.62 | 0.15 | 17.22 | 3.33E-05 | 0.0028091 | 710.20 | 749.78 | 563.01 | 586.66 | 913.54 | 1220.93 | 797.50 | 885.54 |
| B4galt5 | 1035.39 | 0.69 | 0.16 | 17.19 | 3.38E-05 | 0.0028316 | 824.14 | 797.66 | 655.23 | 1039.60 | 919.91 | 1167.52 | 1213.09 | 1665.92 |
| Edem3 | 2156.22 | 0.40 | 0.10 | 17.15 | 3.45E-05 | 0.0028819 | 2084.14 | 2073.14 | 1675.70 | 1756.21 | 2211.61 | 2644.26 | 2279.97 | 2524.72 |
| Whsc1 | 1189.63 | 0.45 | 0.11 | 17.08 | 3.58E-05 | 0.0029685 | 1024.37 | 1101.21 | 982.85 | 994.40 | 1241.77 | 1593.87 | 1138.66 | 1439.93 |
| Colec12 | 3153.54 | 0.61 | 0.15 | 17.04 | 3.66E-05 | 0.0030261 | 2666.01 | 3010.60 | 1863.77 | 3094.32 | 2352.89 | 5151.72 | 3306.09 | 3782.92 |
| Sdhc | 5644.28 | -0.40 | 0.10 | 17.00 | 3.75E-05 | 0.0030624 | 6161.70 | 6305.61 | 7138.39 | 5861.89 | 4821.56 | 4423.66 | 5189.96 | 5251.45 |
| Eepd1 | 1167.15 | -0.55 | 0.13 | 17.00 | 3.74E-05 | 0.0030624 | 1257.78 | 1220.90 | 1583.48 | 1396.49 | 1055.88 | 755.24 | 1006.62 | 1060.80 |
| Nlgn2 | 2573.04 | 0.47 | 0.11 | 16.97 | 3.79E-05 | 0.0030662 | 2132.81 | 2399.67 | 1880.76 | 2441.75 | 2486.73 | 3532.56 | 2877.21 | 2832.81 |
| Pds5a | 1548.66 | 0.40 | 0.10 | 16.96 | 3.81E-05 | 0.0030662 | 1507.79 | 1374.12 | 1227.95 | 1310.80 | 1716.60 | 1915.26 | 1589.69 | 1747.10 |
| Col24a1 | 232.34 | 1.59 | 0.39 | 16.97 | 3.80E-05 | 0.0030662 | 112.84 | 155.13 | 88.58 | 143.13 | 202.89 | 791.78 | 202.92 | 161.43 |
| Nup98 | 1177.83 | 0.54 | 0.13 | 16.92 | 3.91E-05 | 0.0031305 | 1028.79 | 1128.02 | 911.26 | 801.36 | 1343.75 | 1372.73 | 1411.58 | 1425.17 |
| Ash1l | 4010.30 | 0.37 | 0.09 | 16.90 | 3.93E-05 | 0.0031351 | 3709.19 | 3836.03 | 3226.41 | 3387.18 | 4512.45 | 4388.99 | 4279.93 | 4742.26 |
| Luzp1 | 3649.53 | 0.41 | 0.10 | 16.82 | 4.12E-05 | 0.0032327 | 3357.41 | 3009.65 | 3190.01 | 2909.76 | 4153.40 | 4467.70 | 4364.11 | 3744.18 |
| Rnf19a | 1971.52 | 0.36 | 0.09 | 16.81 | 4.14E-05 | 0.0032327 | 1738.99 | 1635.53 | 1708.46 | 1805.18 | 2238.16 | 2090.49 | 2286.17 | 2269.20 |
| Itsn2 | 1914.46 | 0.40 | 0.10 | 16.82 | 4.10E-05 | 0.0032327 | 1798.73 | 1852.90 | 1378.41 | 1718.55 | 2108.57 | 2181.38 | 2029.20 | 2247.98 |
| Dync2h1 | 610.40 | 0.48 | 0.12 | 16.83 | 4.08E-05 | 0.0032327 | 554.22 | 542.94 | 472.01 | 484.96 | 745.70 | 699.95 | 658.38 | 725.04 |
| Qpct | 489.41 | -0.56 | 0.14 | 16.71 | 4.36E-05 | 0.003394 | 553.11 | 612.85 | 587.28 | 592.31 | 421.71 | 399.17 | 319.89 | 428.93 |
| Sprr1a | 30625.68 | -0.53 | 0.13 | 16.68 | 4.43E-05 | 0.0033958 | 37362.91 | 30938.18 | 37736.53 | 33506.45 | 34124.84 | 22732.98 | 29178.82 | 19424.72 |
| Galnt18 | 1198.54 | 0.79 | 0.20 | 16.68 | 4.42E-05 | 0.0033958 | 913.75 | 744.99 | 922.18 | 984.99 | 1127.05 | 2127.97 | 1505.51 | 1261.90 |
| Mogat1 | 32.53 | -1.99 | 0.49 | 16.69 | 4.40E-05 | 0.0033958 | 33.19 | 22.02 | 95.86 | 49.91 | 9.56 | 10.31 | 29.24 | 10.15 |
| Myl4 | 696.50 | -1.18 | 0.29 | 16.64 | 4.53E-05 | 0.0034534 | 1154.90 | 1104.08 | 993.77 | 613.03 | 413.22 | 227.70 | 429.76 | 635.56 |
| Rb1cc1 | 2373.00 | 0.43 | 0.11 | 16.62 | 4.57E-05 | 0.0034566 | 2121.75 | 2279.98 | 1798.25 | 2066.96 | 2763.98 | 2789.50 | 2211.74 | 2951.80 |
| Sectm1a | 14.14 | -6.23 | 1.43 | 16.62 | 4.58E-05 | 0.0034566 | 6.64 | 11.49 | 7.28 | 86.63 | 1.06 | 0.00 | 0.00 | 0.00 |
| Msn | 9664.71 | 0.70 | 0.17 | 16.60 | 4.62E-05 | 0.0034731 | 7282.31 | 6959.63 | 7700.19 | 7671.77 | 7848.98 | 12852.13 | 14579.22 | 12423.41 |
| Synpo | 1965.27 | 0.65 | 0.16 | 16.55 | 4.75E-05 | 0.003555 | 1630.58 | 1355.92 | 1684.19 | 1466.18 | 1780.33 | 2739.84 | 2745.18 | 2319.93 |
| Tgfbr1 | 3390.06 | 0.49 | 0.12 | 16.52 | 4.82E-05 | 0.0035877 | 3258.95 | 3150.41 | 2568.75 | 2598.06 | 3675.39 | 4056.35 | 3072.15 | 4740.41 |
| Ube2i | 2370.73 | 0.46 | 0.11 | 16.49 | 4.88E-05 | 0.0036209 | 2149.40 | 2192.84 | 1641.72 | 2146.06 | 2870.20 | 2730.47 | 2579.48 | 2655.70 |
| Fstl1 | 15974.64 | 0.83 | 0.21 | 16.47 | 4.95E-05 | 0.0036481 | 11675.15 | 12261.70 | 11197.19 | 12052.42 | 12572.81 | 30449.32 | 21082.40 | 16506.12 |
| Cytl1 | 205.14 | -1.26 | 0.31 | 16.46 | 4.97E-05 | 0.0036481 | 314.17 | 291.10 | 311.84 | 226.94 | 144.47 | 53.41 | 127.60 | 171.57 |
| Fam13b | 1893.66 | 0.34 | 0.08 | 16.40 | 5.13E-05 | 0.0037511 | 1774.39 | 1805.02 | 1528.88 | 1688.41 | 1971.54 | 2115.79 | 1982.24 | 2283.04 |
| Arhgef12 | 7201.81 | 0.33 | 0.08 | 16.36 | 5.23E-05 | 0.003795 | 6557.73 | 6751.84 | 6238.05 | 5975.83 | 8315.31 | 7886.88 | 7766.78 | 8122.08 |
| St6gal2 | 98.87 | -1.40 | 0.34 | 16.36 | 5.24E-05 | 0.003795 | 112.84 | 283.44 | 63.10 | 147.84 | 29.74 | 57.16 | 22.15 | 74.72 |
| Glg1 | 7859.81 | 0.35 | 0.09 | 16.31 | 5.38E-05 | 0.0038618 | 6965.93 | 7433.63 | 6862.95 | 6435.36 | 8358.86 | 9370.18 | 8889.49 | 8562.08 |
| Ankib1 | 1649.47 | 0.38 | 0.09 | 16.31 | 5.37E-05 | 0.0038618 | 1532.13 | 1510.09 | 1323.81 | 1469.00 | 1860.00 | 2006.15 | 1610.95 | 1883.62 |
| Acss2 | 2949.29 | -0.81 | 0.20 | 16.26 | 5.51E-05 | 0.0039411 | 3300.99 | 2500.22 | 6555.96 | 1844.73 | 2391.13 | 1554.51 | 3741.17 | 1705.59 |
| Rgma | 947.83 | -0.63 | 0.16 | 16.21 | 5.68E-05 | 0.0040425 | 848.48 | 1130.89 | 1338.37 | 1255.24 | 823.25 | 726.19 | 675.22 | 785.00 |
| Slc7a10 | 197.39 | -1.36 | 0.34 | 16.10 | 6.00E-05 | 0.0042528 | 155.98 | 188.64 | 411.34 | 344.65 | 115.79 | 52.47 | 172.79 | 137.44 |
| Grem2 | 1396.82 | 0.69 | 0.18 | 16.06 | 6.13E-05 | 0.0043205 | 866.18 | 1024.60 | 1258.29 | 1116.82 | 1581.69 | 2264.77 | 1582.60 | 1479.59 |
| Dlk1 | 74.98 | -1.76 | 0.44 | 16.04 | 6.21E-05 | 0.0043604 | 103.99 | 126.40 | 148.03 | 85.69 | 31.87 | 18.74 | 16.84 | 68.26 |
| Chd2 | 2674.08 | 0.33 | 0.08 | 16.02 | 6.26E-05 | 0.0043763 | 2604.06 | 2391.06 | 2094.32 | 2570.76 | 2738.49 | 3195.23 | 2783.28 | 3015.45 |
| Kras | 1967.24 | 0.38 | 0.09 | 16.00 | 6.33E-05 | 0.0044055 | 1982.36 | 1767.68 | 1420.88 | 1865.45 | 2079.89 | 2301.31 | 1988.44 | 2331.93 |
| Rbm24 | 1464.79 | -0.90 | 0.23 | 15.98 | 6.42E-05 | 0.004448 | 1648.28 | 2257.00 | 1788.54 | 2050.01 | 1014.45 | 782.41 | 637.12 | 1540.47 |
| Tmtc3 | 1448.47 | 0.48 | 0.12 | 15.95 | 6.49E-05 | 0.00448 | 1375.04 | 1364.54 | 971.93 | 1288.20 | 1725.10 | 1753.16 | 1344.23 | 1765.55 |
| F3 | 1414.53 | 0.46 | 0.12 | 15.93 | 6.56E-05 | 0.0045103 | 1396.06 | 1091.63 | 1139.38 | 1233.59 | 1531.77 | 1797.20 | 1383.22 | 1743.41 |
| Fbln7 | 135.09 | 1.19 | 0.30 | 15.91 | 6.64E-05 | 0.0045403 | 69.69 | 56.50 | 89.79 | 98.88 | 196.52 | 225.82 | 237.48 | 106.08 |
| Cdo1 | 2991.35 | -0.96 | 0.24 | 15.85 | 6.85E-05 | 0.0046679 | 2927.08 | 2156.45 | 7390.78 | 2776.98 | 1753.78 | 1304.33 | 3620.66 | 2000.77 |
| 9930013L23R | 373.26 | 1.24 | 0.31 | 15.79 | 7.07E-05 | 0.004794 | 245.58 | 189.60 | 220.84 | 238.24 | 268.75 | 729.00 | 741.68 | 352.37 |
| Cldn10 | 288.82 | -0.89 | 0.23 | 15.61 | 7.79E-05 | 0.0052629 | 417.05 | 360.05 | 414.98 | 317.34 | 190.14 | 156.48 | 163.04 | 291.49 |
| Sp6 | 730.13 | -0.52 | 0.13 | 15.58 | 7.90E-05 | 0.0053118 | 766.62 | 851.28 | 935.53 | 814.54 | 764.82 | 505.99 | 599.90 | 602.35 |
| Map1b | 3389.11 | 0.39 | 0.10 | 15.57 | 7.94E-05 | 0.0053173 | 3125.10 | 2921.55 | 2880.60 | 2996.39 | 3117.71 | 4157.55 | 3658.76 | 4255.21 |
| Slc25a10 | 3503.74 | -0.39 | 0.10 | 15.55 | 8.02E-05 | 0.0053509 | 3541.04 | 3124.56 | 5272.19 | 3274.18 | 3473.56 | 2631.15 | 4101.82 | 2611.42 |
| Mybl1 | 161.02 | 0.70 | 0.18 | 15.51 | 8.22E-05 | 0.0054623 | 143.81 | 138.85 | 87.36 | 156.32 | 149.78 | 214.58 | 137.35 | 260.13 |
| Prrg4 | 366.68 | 0.78 | 0.19 | 15.50 | 8.26E-05 | 0.0054648 | 298.68 | 288.23 | 249.96 | 226.00 | 583.18 | 497.56 | 443.94 | 345.91 |
| Hadh | 4405.79 | -0.36 | 0.09 | 15.46 | 8.41E-05 | 0.0055098 | 4668.29 | 4385.68 | 5846.13 | 4612.29 | 4171.46 | 3544.74 | 4052.20 | 3965.57 |
| Abl2 | 1470.40 | 0.42 | 0.11 | 15.47 | 8.37E-05 | 0.0055098 | 1391.64 | 1258.25 | 1107.83 | 1301.39 | 1760.15 | 1666.95 | 1674.75 | 1602.28 |
| Myo18b | 453.54 | 0.82 | 0.21 | 15.46 | 8.43E-05 | 0.0055098 | 344.04 | 338.02 | 317.91 | 336.18 | 524.75 | 408.54 | 536.98 | 821.89 |
| Mdh1 | 7132.63 | -0.32 | 0.08 | 15.44 | 8.50E-05 | 0.0055291 | 7515.73 | 7681.64 | 8976.68 | 7105.83 | 6708.12 | 5861.98 | 6775.22 | 6435.86 |
| Dst | 14814.37 | 0.40 | 0.10 | 15.38 | 8.79E-05 | 0.005647 | 14293.59 | 12185.10 | 11870.63 | 12669.21 | 16925.92 | 16937.53 | 18160.89 | 15472.07 |
| Cox6b1 | 4057.53 | -0.40 | 0.10 | 15.39 | 8.77E-05 | 0.005647 | 4461.42 | 4823.29 | 5058.64 | 3853.31 | 3640.34 | 3284.25 | 3891.81 | 3447.15 |
| Cacna1g | 196.46 | 0.75 | 0.19 | 15.39 | 8.75E-05 | 0.005647 | 147.13 | 180.98 | 143.18 | 133.72 | 175.27 | 300.78 | 205.58 | 285.03 |
| Bptf | 3652.18 | 0.39 | 0.10 | 15.36 | 8.89E-05 | 0.0056933 | 3618.48 | 3257.66 | 2855.11 | 3032.17 | 4084.36 | 4335.58 | 4082.32 | 3951.73 |
| Chchd10 | 1527.11 | -0.66 | 0.17 | 15.35 | 8.94E-05 | 0.0056986 | 1744.52 | 1887.37 | 2276.33 | 1387.08 | 1495.65 | 849.87 | 1244.99 | 1331.08 |
| Spock3 | 1203.17 | -1.74 | 0.43 | 15.32 | 9.06E-05 | 0.0057528 | 1436.99 | 2699.39 | 1526.45 | 1834.37 | 661.78 | 319.52 | 330.52 | 816.36 |
| Ralgps1 | 830.33 | 0.49 | 0.12 | 15.30 | 9.18E-05 | 0.0058046 | 787.64 | 736.37 | 566.65 | 739.21 | 849.80 | 928.58 | 1002.19 | 1032.21 |
| Zfp445 | 1042.01 | 0.67 | 0.17 | 15.28 | 9.25E-05 | 0.0058308 | 922.60 | 1063.86 | 615.19 | 730.73 | 1377.74 | 1321.19 | 1067.77 | 1236.99 |
| Ncam1 | 1146.89 | 0.58 | 0.15 | 15.25 | 9.41E-05 | 0.0059028 | 790.95 | 1054.29 | 894.27 | 1034.89 | 985.77 | 1591.99 | 1230.81 | 1592.13 |
| Gabpb2 | 2884.86 | 0.39 | 0.10 | 15.22 | 9.59E-05 | 0.005995 | 2653.85 | 2679.28 | 2245.99 | 2544.39 | 3274.92 | 3654.37 | 2998.61 | 3027.44 |
| Gpr68 | 152.96 | 1.03 | 0.27 | 15.14 | 9.96E-05 | 0.0062034 | 111.73 | 75.65 | 115.27 | 108.29 | 128.53 | 289.54 | 160.39 | 234.30 |
| Flrt2 | 1313.11 | 0.66 | 0.17 | 15.12 | 0.0001007 | 0.0062461 | 1279.91 | 1085.89 | 826.32 | 1095.16 | 1046.32 | 2264.77 | 1441.71 | 1464.83 |
| Rbp4 | 1032.07 | -0.63 | 0.16 | 15.10 | 0.0001019 | 0.0062967 | 1101.80 | 869.47 | 1633.23 | 1253.36 | 797.75 | 737.43 | 1143.97 | 719.50 |
| Stk17b | 2121.40 | 0.60 | 0.16 | 15.09 | 0.0001028 | 0.0063242 | 1646.07 | 1694.90 | 1811.60 | 1698.77 | 1963.04 | 2707.98 | 2003.50 | 3445.31 |
| Mapt | 204.33 | -0.96 | 0.24 | 15.05 | 0.0001046 | 0.0064001 | 259.96 | 305.47 | 232.97 | 295.68 | 132.78 | 89.02 | 122.28 | 196.48 |
| Gdpd2 | 34.08 | 2.02 | 0.53 | 15.05 | 0.0001048 | 0.0064001 | 5.53 | 19.15 | 15.77 | 15.07 | 32.93 | 101.20 | 45.19 | 37.82 |
| Irs3 | 70.09 | -1.27 | 0.33 | 15.03 | 0.0001056 | 0.0064148 | 82.97 | 45.96 | 173.52 | 80.04 | 40.37 | 35.61 | 70.00 | 32.29 |
| Alb | 26.59 | -1.99 | 0.51 | 15.03 | 0.0001058 | 0.0064148 | 24.34 | 52.67 | 57.03 | 32.02 | 14.87 | 3.75 | 13.29 | 14.76 |
| Col4a1 | 39024.02 | 0.83 | 0.22 | 15.01 | 0.0001069 | 0.0064561 | 27461.05 | 22224.28 | 34428.82 | 28827.30 | 29951.25 | 65594.04 | 54875.22 | 48830.23 |
| Senp5 | 624.28 | 0.47 | 0.12 | 14.98 | 0.0001085 | 0.0065015 | 582.98 | 538.16 | 516.91 | 481.19 | 636.29 | 775.85 | 723.95 | 738.87 |
| Bhlhe22 | 23.85 | 3.61 | 0.92 | 14.99 | 0.0001082 | 0.0065015 | 4.42 | 4.79 | 1.21 | 8.48 | 9.56 | 130.25 | 27.47 | 4.61 |
| Dip2b | 2116.45 | 0.49 | 0.12 | 14.97 | 0.000109 | 0.0065053 | 1933.69 | 1692.03 | 1560.42 | 1896.52 | 2683.25 | 2610.53 | 2427.06 | 2128.07 |
| Bdkrb2 | 136.90 | 0.92 | 0.24 | 14.94 | 0.0001109 | 0.0065966 | 107.30 | 75.65 | 100.71 | 99.82 | 125.35 | 208.02 | 199.38 | 178.95 |
| Naip5 | 821.44 | 0.75 | 0.19 | 14.91 | 0.0001128 | 0.0066822 | 756.66 | 564.97 | 486.57 | 777.82 | 806.25 | 1508.60 | 829.40 | 841.26 |
| Trim71 | 31.64 | -1.65 | 0.43 | 14.89 | 0.0001141 | 0.0067373 | 43.14 | 63.20 | 42.47 | 46.14 | 13.81 | 21.55 | 5.32 | 17.53 |
| Mgst1 | 11078.75 | -0.37 | 0.10 | 14.88 | 0.0001146 | 0.0067384 | 11803.47 | 11531.08 | 14798.54 | 10816.00 | 11323.60 | 8371.31 | 10137.13 | 9848.88 |
| Cox8a | 2737.62 | -0.41 | 0.11 | 14.85 | 0.0001164 | 0.0067971 | 3052.09 | 2955.07 | 3626.83 | 2451.16 | 2776.73 | 1987.41 | 2870.12 | 2181.57 |
| Pik3c2a | 1471.63 | 0.44 | 0.11 | 14.85 | 0.0001164 | 0.0067971 | 1286.55 | 1481.36 | 970.71 | 1459.59 | 1478.65 | 1863.73 | 1546.27 | 1686.22 |
| Abcc2 | 47.20 | -1.85 | 0.47 | 14.83 | 0.0001176 | 0.0068402 | 77.44 | 109.16 | 50.96 | 64.98 | 18.06 | 9.37 | 7.98 | 39.66 |
| Slc25a1 | 3967.74 | -0.78 | 0.21 | 14.82 | 0.0001183 | 0.0068599 | 4192.61 | 3131.26 | 8951.20 | 2947.42 | 2805.41 | 2254.46 | 5001.22 | 2458.30 |
| mt-Rnr1 | 10007.20 | -0.76 | 0.20 | 14.79 | 0.0001205 | 0.0069325 | 10511.40 | 11453.51 | 15976.75 | 12455.45 | 7384.77 | 6529.14 | 5409.72 | 10336.85 |

| | | | | | | | | | | | | | | |
|---|---|---|---|---|---|---|---|---|---|---|---|---|---|---|
| Ugcg | 1927.22 | 0.43 | 0.11 | 14.79 | 0.0001204 | 0.0069325 | 1733.46 | 1660.43 | 1362.64 | 2052.84 | 1925.86 | 2625.52 | 1820.08 | 2236.91 |
| Col7a1 | 518.89 | 1.05 | 0.27 | 14.77 | 0.0001213 | 0.0069544 | 380.54 | 387.82 | 291.21 | 355.01 | 543.87 | 1229.37 | 555.59 | 407.72 |
| Frrs1 | 594.51 | 0.61 | 0.16 | 14.75 | 0.0001229 | 0.0070202 | 594.05 | 409.84 | 441.68 | 439.76 | 713.83 | 765.54 | 755.85 | 635.56 |
| Eogt | 1549.09 | 0.43 | 0.11 | 14.70 | 0.0001258 | 0.0071615 | 1514.43 | 1370.28 | 1138.16 | 1393.67 | 1526.46 | 2082.99 | 1692.48 | 1674.23 |
| Herc4 | 2361.25 | 0.38 | 0.10 | 14.65 | 0.0001294 | 0.0073163 | 2310.91 | 2211.03 | 1801.89 | 2124.40 | 2350.76 | 2582.42 | 2389.85 | 3118.77 |
| Hipk2 | 779.86 | 0.73 | 0.18 | 14.65 | 0.0001293 | 0.0073163 | 700.24 | 597.52 | 440.46 | 611.14 | 956.03 | 932.33 | 1205.11 | 796.06 |
| Zzef1 | 2653.23 | 0.42 | 0.11 | 14.61 | 0.0001321 | 0.0074401 | 2368.44 | 2488.73 | 2040.93 | 2222.34 | 3277.05 | 3105.28 | 2964.05 | 2759.02 |
| Edn3 | 206.47 | -1.21 | 0.32 | 14.61 | 0.0001326 | 0.0074401 | 288.73 | 233.65 | 340.96 | 289.09 | 160.40 | 84.33 | 67.34 | 187.26 |
| Large | 1832.98 | 0.46 | 0.12 | 14.58 | 0.0001341 | 0.0075027 | 1562.00 | 1583.82 | 1544.65 | 1499.14 | 1772.90 | 2044.57 | 2401.37 | 2255.36 |
| Pbx1 | 7943.47 | 0.30 | 0.08 | 14.56 | 0.0001361 | 0.0075411 | 7210.41 | 8012.00 | 6483.16 | 7384.56 | 8191.02 | 9348.62 | 7774.76 | 9143.22 |
| Ccdc93 | 1198.18 | 0.45 | 0.12 | 14.57 | 0.0001354 | 0.0075411 | 1130.57 | 1184.52 | 845.74 | 996.29 | 1389.43 | 1457.06 | 1193.59 | 1388.27 |
| H2-Q10 | 45.83 | -1.85 | 0.49 | 14.55 | 0.0001363 | 0.0075411 | 54.21 | 30.64 | 160.17 | 28.25 | 17.00 | 6.56 | 51.39 | 18.45 |
| Etfb | 3321.65 | -0.37 | 0.10 | 14.53 | 0.0001383 | 0.0075954 | 3888.40 | 3556.42 | 4018.76 | 3308.08 | 2932.88 | 2712.67 | 3384.07 | 2771.93 |
| Cdcp1 | 2084.80 | 0.34 | 0.09 | 14.52 | 0.0001389 | 0.0075954 | 1798.73 | 1763.85 | 1917.16 | 1725.14 | 2650.32 | 2266.65 | 2504.16 | 2052.43 |
| Kcnk3 | 1930.88 | -0.68 | 0.18 | 14.51 | 0.0001396 | 0.0075954 | 1929.26 | 2329.77 | 3106.29 | 2066.02 | 1286.39 | 1240.61 | 1619.81 | 1868.86 |
| Trim7 | 816.56 | 0.45 | 0.12 | 14.50 | 0.0001401 | 0.0075954 | 761.09 | 667.43 | 623.68 | 707.19 | 942.22 | 949.20 | 1015.49 | 866.17 |
| Sostdc1 | 161.96 | -1.20 | 0.30 | 14.51 | 0.0001397 | 0.0075954 | 146.02 | 255.67 | 192.93 | 306.04 | 126.41 | 63.72 | 87.73 | 117.15 |
| Asgr2 | 38.47 | -1.67 | 0.43 | 14.50 | 0.0001401 | 0.0075954 | 56.42 | 48.84 | 41.26 | 93.23 | 8.50 | 24.36 | 20.38 | 14.76 |
| Eln | 4691.83 | 1.53 | 0.40 | 14.48 | 0.0001415 | 0.0076477 | 2545.43 | 1973.55 | 2455.91 | 2916.35 | 3665.83 | 10789.76 | 8309.97 | 4877.86 |
| Aak1 | 444.73 | 0.68 | 0.18 | 14.47 | 0.0001421 | 0.0076541 | 388.29 | 331.32 | 349.46 | 264.61 | 593.80 | 579.08 | 637.12 | 414.18 |
| Ern2 | 247.15 | 1.10 | 0.29 | 14.46 | 0.000143 | 0.007674 | 199.12 | 137.89 | 138.33 | 173.27 | 357.98 | 512.55 | 256.97 | 201.09 |
| Hpse2 | 344.79 | -0.81 | 0.21 | 14.43 | 0.0001455 | 0.0077284 | 405.99 | 527.62 | 356.74 | 511.33 | 226.26 | 201.46 | 170.13 | 358.83 |
| B3galt2 | 262.11 | -1.21 | 0.32 | 14.44 | 0.000145 | 0.0077284 | 205.76 | 278.65 | 401.63 | 572.53 | 160.40 | 113.38 | 154.18 | 210.32 |
| Uprt | 124.69 | 0.94 | 0.24 | 14.44 | 0.0001449 | 0.0077284 | 97.35 | 116.82 | 67.95 | 65.92 | 171.02 | 160.23 | 158.61 | 159.58 |
| Ndufa4 | 5579.47 | -0.51 | 0.13 | 14.42 | 0.0001462 | 0.0077415 | 6584.28 | 7195.19 | 6816.84 | 5448.49 | 4862.99 | 3823.97 | 4685.77 | 5218.24 |
| Golim4 | 5654.55 | 0.37 | 0.10 | 14.38 | 0.0001491 | 0.007868 | 4950.38 | 5798.10 | 4322.11 | 5087.84 | 5988.98 | 6248.97 | 6061.01 | 6779.00 |
| Cfp | 1140.95 | -0.61 | 0.16 | 14.35 | 0.0001516 | 0.0079772 | 1170.39 | 1615.42 | 1227.95 | 1602.72 | 710.65 | 807.71 | 871.05 | 1121.69 |
| Mpc2 | 1757.90 | -0.51 | 0.14 | 14.34 | 0.0001525 | 0.007995 | 2023.29 | 1797.36 | 2732.56 | 1490.66 | 1573.19 | 1247.17 | 1756.28 | 1442.69 |
| Upk3b | 10553.59 | -0.55 | 0.14 | 14.31 | 0.0001547 | 0.0080839 | 14152.00 | 10485.41 | 13864.23 | 9780.17 | 11523.31 | 6719.35 | 10133.59 | 7770.63 |
| Igf1r | 785.09 | 0.67 | 0.17 | 14.27 | 0.0001581 | 0.0082076 | 792.06 | 614.76 | 554.52 | 457.65 | 1038.88 | 907.97 | 992.45 | 922.44 |
| Clca5 | 545.23 | 1.02 | 0.27 | 14.28 | 0.0001577 | 0.0082076 | 443.60 | 259.50 | 424.69 | 266.49 | 549.18 | 895.79 | 1042.96 | 479.67 |
| Bgn | 38154.48 | 0.72 | 0.19 | 14.24 | 0.0001605 | 0.0082104 | 29512.00 | 33696.94 | 22971.96 | 35368.13 | 24380.80 | 61029.82 | 49018.89 | 49257.32 |
| Rmdn3 | 1400.49 | -0.33 | 0.09 | 14.25 | 0.0001602 | 0.0082104 | 1521.07 | 1444.02 | 1638.08 | 1600.84 | 1305.51 | 1199.38 | 1263.60 | 1231.46 |
| F2r | 776.51 | 0.74 | 0.20 | 14.26 | 0.0001591 | 0.0082104 | 582.98 | 600.40 | 501.13 | 730.73 | 585.30 | 1243.42 | 1044.73 | 923.36 |
| Itgb1bp2 | 134.36 | -0.72 | 0.19 | 14.24 | 0.0001607 | 0.0082104 | 178.10 | 168.53 | 158.95 | 159.14 | 111.54 | 89.95 | 107.22 | 101.47 |
| Tcap | 113.86 | -1.80 | 0.46 | 14.26 | 0.0001595 | 0.0082104 | 97.35 | 264.29 | 232.97 | 90.40 | 69.05 | 14.06 | 72.66 | 70.11 |
| Kcnip1 | 37.12 | -1.67 | 0.45 | 14.20 | 0.0001643 | 0.0083689 | 49.78 | 74.69 | 49.75 | 54.62 | 10.62 | 34.67 | 6.20 | 16.60 |
| Htra4 | 68.60 | 1.25 | 0.32 | 14.16 | 0.0001675 | 0.0085035 | 48.67 | 39.26 | 23.05 | 60.27 | 87.10 | 97.45 | 107.22 | 85.79 |
| Nin | 1759.87 | 0.41 | 0.11 | 14.14 | 0.0001701 | 0.00861 | 1659.34 | 1624.04 | 1209.75 | 1761.86 | 1956.67 | 2124.22 | 1714.63 | 2028.44 |
| Slc36a2 | 326.25 | -1.13 | 0.31 | 14.09 | 0.0001739 | 0.0087745 | 306.43 | 266.20 | 845.74 | 308.87 | 159.34 | 110.57 | 396.98 | 215.85 |
| Bcl2l11 | 1445.45 | 0.40 | 0.11 | 14.09 | 0.0001745 | 0.0087749 | 1248.93 | 1308.04 | 1147.87 | 1342.82 | 1478.65 | 1774.71 | 1712.86 | 1549.70 |
| Gstz1 | 2098.20 | -0.47 | 0.12 | 14.06 | 0.0001768 | 0.0088332 | 2121.75 | 2125.81 | 2901.22 | 2387.13 | 1812.20 | 1439.26 | 2105.40 | 1892.84 |
| Slc39a14 | 1437.91 | 0.37 | 0.10 | 14.07 | 0.0001764 | 0.0088332 | 1320.84 | 1338.68 | 1113.90 | 1373.89 | 1318.25 | 1742.85 | 1578.17 | 1716.66 |
| Pmvk | 570.56 | -0.61 | 0.16 | 14.05 | 0.0001777 | 0.0088531 | 639.40 | 613.80 | 899.12 | 500.03 | 452.52 | 382.30 | 677.88 | 399.42 |
| Rbm33 | 3317.56 | 0.40 | 0.11 | 14.03 | 0.0001797 | 0.008895 | 2950.31 | 3264.36 | 2424.36 | 3011.46 | 3911.21 | 3840.83 | 3462.04 | 3675.92 |
| Suv420h1 | 2983.88 | 0.37 | 0.10 | 14.03 | 0.0001796 | 0.008895 | 2623.98 | 2921.55 | 2332.14 | 2786.40 | 3311.04 | 3689.04 | 2795.69 | 3411.18 |
| Cd80 | 72.64 | 0.98 | 0.26 | 14.01 | 0.0001815 | 0.0089563 | 63.06 | 53.62 | 36.40 | 55.56 | 62.67 | 112.44 | 91.27 | 106.08 |
| Dopey2 | 1613.87 | 0.36 | 0.10 | 14.00 | 0.0001831 | 0.0089845 | 1579.70 | 1416.25 | 1218.25 | 1562.23 | 1727.22 | 1824.37 | 1682.73 | 1900.22 |
| Synj1 | 1465.79 | 0.32 | 0.09 | 14.00 | 0.0001832 | 0.0089845 | 1358.45 | 1408.59 | 1197.62 | 1367.30 | 1469.09 | 1800.95 | 1439.93 | 1684.37 |
| Uqcr11 | 1708.61 | -0.53 | 0.14 | 13.96 | 0.0001869 | 0.0091256 | 2027.72 | 2084.63 | 2318.79 | 1517.03 | 1377.74 | 1223.74 | 1653.49 | 1465.76 |
| Ttyh1 | 152.94 | -0.72 | 0.20 | 13.96 | 0.0001872 | 0.0091256 | 146.02 | 206.84 | 218.41 | 183.63 | 121.10 | 117.13 | 117.85 | 112.54 |
| Ighg3 | 27.46 | 4.02 | 1.03 | 13.94 | 0.0001889 | 0.0091781 | 3.32 | 0.00 | 6.07 | 1.88 | 133.84 | 4.69 | 6.20 | 63.65 |
| Rapgef2 | 775.44 | 0.43 | 0.12 | 13.93 | 0.0001895 | 0.0091803 | 714.62 | 727.75 | 587.28 | 687.42 | 719.14 | 1035.40 | 844.47 | 887.39 |
| Golgb1 | 5726.02 | 0.38 | 0.10 | 13.90 | 0.0001931 | 0.0093239 | 5448.18 | 4970.75 | 4800.18 | 4625.48 | 6437.25 | 6844.91 | 6870.04 | 5811.37 |
| Atxn3 | 1626.75 | 0.37 | 0.10 | 13.89 | 0.0001936 | 0.0093239 | 1482.35 | 1610.64 | 1269.21 | 1432.28 | 1804.77 | 1963.99 | 1532.09 | 1918.67 |
| Zcchc6 | 3085.70 | 0.32 | 0.09 | 13.88 | 0.0001949 | 0.0093597 | 2888.37 | 2762.59 | 2432.85 | 3090.56 | 3178.26 | 3645.94 | 3255.58 | 3431.47 |
| Mmp9 | 708.41 | -1.85 | 0.50 | 13.87 | 0.0001959 | 0.009376 | 775.47 | 1065.78 | 1442.72 | 1149.78 | 249.63 | 138.68 | 286.21 | 559.00 |
| Cdkn1c | 486.90 | -0.57 | 0.15 | 13.86 | 0.0001973 | 0.0094147 | 491.17 | 596.57 | 749.88 | 403.98 | 458.89 | 348.57 | 510.40 | 335.77 |
| Naip6 | 849.49 | 0.77 | 0.21 | 13.84 | 0.000199 | 0.0094689 | 900.47 | 523.79 | 493.85 | 726.97 | 857.24 | 1522.65 | 879.91 | 891.08 |
| Plekhh2 | 1059.31 | 0.72 | 0.20 | 13.83 | 0.0002004 | 0.0095093 | 933.66 | 739.25 | 883.35 | 688.36 | 867.86 | 1663.21 | 1378.79 | 1320.01 |
| Stox2 | 1825.03 | 0.37 | 0.10 | 13.82 | 0.0002017 | 0.0095407 | 1604.03 | 1589.57 | 1545.86 | 1664.87 | 2061.83 | 2256.34 | 2007.93 | 1869.78 |
| Ankrd17 | 4057.16 | 0.30 | 0.08 | 13.79 | 0.0002042 | 0.0096156 | 3773.35 | 3868.59 | 3351.39 | 3692.28 | 4467.83 | 4654.17 | 4299.42 | 4350.22 |
| Cd209b | 107.63 | -1.63 | 0.44 | 13.79 | 0.0002045 | 0.0096156 | 119.47 | 206.84 | 179.58 | 146.90 | 24.43 | 22.49 | 69.12 | 92.24 |
| C1qtnf7 | 1003.19 | 0.73 | 0.19 | 13.77 | 0.0002062 | 0.0096676 | 827.46 | 962.36 | 598.20 | 791.00 | 868.92 | 1787.83 | 1195.37 | 994.39 |
| Nr2c2 | 2267.81 | 0.33 | 0.09 | 13.75 | 0.0002084 | 0.0097133 | 2156.04 | 2062.61 | 1888.04 | 1987.86 | 2597.21 | 2604.91 | 2402.25 | 2443.54 |
| Mum1l1 | 464.31 | 0.45 | 0.12 | 13.76 | 0.0002081 | 0.0097133 | 440.28 | 476.87 | 296.07 | 441.64 | 427.03 | 657.79 | 409.38 | 565.46 |
| Whsc1l1 | 3542.54 | 0.28 | 0.08 | 13.74 | 0.0002095 | 0.0097156 | 3225.77 | 3204.03 | 3141.48 | 3285.48 | 3830.48 | 4099.45 | 3763.32 | 3790.30 |
| Pdpr | 958.08 | 0.54 | 0.15 | 13.74 | 0.0002097 | 0.0097156 | 823.03 | 891.50 | 799.63 | 605.49 | 1207.78 | 1196.57 | 1042.96 | 1097.70 |
| Olfm1 | 1573.08 | -0.43 | 0.12 | 13.72 | 0.0002123 | 0.0098079 | 1598.50 | 1782.04 | 2003.31 | 1852.26 | 1270.45 | 1326.82 | 1246.76 | 1504.50 |
| Sulf1 | 3189.42 | 0.68 | 0.18 | 13.70 | 0.0002148 | 0.0098683 | 2789.91 | 2572.99 | 2538.42 | 2046.25 | 2802.22 | 5160.16 | 4187.77 | 3417.64 |
| Fxyd6 | 1541.22 | 0.87 | 0.24 | 13.70 | 0.0002145 | 0.0098683 | 1084.11 | 1080.14 | 1254.65 | 952.97 | 1285.32 | 2921.62 | 2287.06 | 1463.91 |
| Larp4b | 1817.52 | 0.39 | 0.10 | 13.69 | 0.0002155 | 0.0098706 | 1569.74 | 1672.88 | 1396.62 | 1760.92 | 2145.75 | 2109.23 | 1806.78 | 2078.26 |
| Add2 | 246.45 | -1.09 | 0.29 | 13.68 | 0.0002169 | 0.0099054 | 329.66 | 378.24 | 265.73 | 381.38 | 182.71 | 80.58 | 134.69 | 218.62 |
| Slc6a8 | 5424.96 | 0.33 | 0.09 | 13.65 | 0.0002201 | 0.0099088 | 5044.41 | 4843.40 | 4784.41 | 4599.11 | 5843.45 | 5622.11 | 6131.90 | 6530.87 |
| Grem1 | 376.97 | 5.26 | 1.24 | 13.66 | 0.0002188 | 0.0099088 | 21.02 | 40.22 | 10.92 | 9.42 | 41.43 | 2403.45 | 440.40 | 48.89 |
| Tshr | 186.24 | -1.53 | 0.42 | 13.66 | 0.0002192 | 0.0099088 | 159.30 | 168.53 | 524.19 | 217.53 | 71.17 | 28.11 | 199.38 | 121.76 |
| Hmgcll1 | 172.82 | 0.74 | 0.20 | 13.67 | 0.0002182 | 0.0099088 | 148.23 | 142.68 | 86.15 | 172.33 | 208.20 | 235.19 | 167.48 | 222.31 |
| Klb | 80.32 | -1.41 | 0.38 | 13.65 | 0.0002199 | 0.0099088 | 75.22 | 69.90 | 211.13 | 80.98 | 41.43 | 15.93 | 103.68 | 44.28 |
| Lipe | 2190.97 | -0.91 | 0.25 | 13.64 | 0.0002211 | 0.0099297 | 1758.91 | 1855.77 | 4455.58 | 2984.15 | 1315.07 | 987.62 | 2535.17 | 1635.48 |
| Nhsl2 | 1893.41 | 0.50 | 0.14 | 13.60 | 0.0002265 | 0.0101438 | 1531.02 | 2092.29 | 1223.10 | 1741.15 | 1575.32 | 2708.92 | 1958.31 | 2317.17 |
| Glrx | 824.18 | -0.44 | 0.12 | 13.58 | 0.000228 | 0.0101803 | 870.60 | 906.82 | 928.25 | 1060.32 | 761.63 | 646.54 | 728.39 | 690.91 |
| Pde9a | 612.84 | -0.64 | 0.17 | 13.57 | 0.0002301 | 0.0102184 | 730.11 | 842.66 | 629.75 | 866.34 | 415.34 | 443.21 | 308.37 | 666.92 |
| Vash1 | 135.60 | 0.96 | 0.26 | 13.57 | 0.0002297 | 0.0102184 | 90.71 | 86.18 | 88.58 | 104.53 | 129.59 | 213.64 | 229.50 | 142.06 |
| Fmn1 | 475.21 | 0.55 | 0.15 | 13.55 | 0.000232 | 0.0102733 | 417.05 | 376.33 | 361.59 | 422.81 | 501.38 | 721.50 | 490.91 | 510.11 |
| Dnal1 | 1003.41 | 0.41 | 0.11 | 13.53 | 0.0002353 | 0.0103888 | 934.76 | 976.72 | 708.62 | 944.49 | 1105.80 | 1276.22 | 908.27 | 1172.42 |
| Fnip2 | 841.79 | 0.62 | 0.17 | 13.51 | 0.0002379 | 0.0104373 | 797.59 | 681.79 | 567.87 | 613.97 | 1216.28 | 1084.13 | 927.76 | 844.95 |
| Pappa | 340.83 | 0.75 | 0.20 | 13.50 | 0.0002383 | 0.0104373 | 272.13 | 292.06 | 209.92 | 294.74 | 303.80 | 641.86 | 360.65 | 351.45 |
| Gan | 226.82 | 0.89 | 0.24 | 13.51 | 0.0002375 | 0.0104373 | 191.38 | 206.84 | 121.34 | 124.30 | 316.55 | 322.33 | 303.05 | 228.76 |
| 4931406P16R | 2296.54 | 0.46 | 0.13 | 13.49 | 0.0002395 | 0.0104523 | 1786.56 | 1808.85 | 2089.46 | 2055.66 | 2365.63 | 3100.59 | 2725.68 | 2439.85 |
| Shroom4 | 784.20 | 0.50 | 0.14 | 13.48 | 0.0002407 | 0.0104523 | 710.20 | 709.56 | 594.56 | 630.92 | 859.36 | 1113.18 | 854.21 | 801.60 |
| Dyrk2 | 720.00 | 0.40 | 0.11 | 13.49 | 0.0002404 | 0.0104523 | 633.87 | 639.66 | 607.91 | 597.96 | 794.56 | 758.05 | 871.05 | 856.95 |
| Bicc1 | 3677.39 | 0.56 | 0.15 | 13.47 | 0.0002425 | 0.0105057 | 3310.95 | 3189.67 | 2539.63 | 3324.09 | 2952.00 | 5872.29 | 4195.75 | 4034.75 |
| Ankrd63 | 39.33 | -1.74 | 0.48 | 13.44 | 0.0002465 | 0.0106484 | 23.23 | 45.96 | 103.14 | 67.80 | 17.00 | 12.18 | 19.49 | 25.83 |

| | | | | | | | | | | | | | | |
|---|---|---|---|---|---|---|---|---|---|---|---|---|---|---|
| Ednrb | 983.48 | -0.69 | 0.19 | 13.40 | 0.0002517 | 0.0108443 | 1445.84 | 1623.08 | 946.45 | 998.17 | 583.18 | 646.54 | 625.60 | 999.00 |
| Rai14 | 3643.70 | 0.34 | 0.09 | 13.38 | 0.0002539 | 0.0109102 | 3242.36 | 3364.91 | 3369.59 | 2866.44 | 3903.78 | 4249.37 | 4101.82 | 4051.35 |
| Fcor | 51.92 | -1.23 | 0.34 | 13.36 | 0.0002564 | 0.0109859 | 69.69 | 58.41 | 110.42 | 46.14 | 29.74 | 21.55 | 43.42 | 35.98 |
| Enpp2 | 2831.77 | -0.52 | 0.14 | 13.36 | 0.0002574 | 0.0109989 | 3617.37 | 3829.33 | 2789.59 | 3447.45 | 2195.67 | 2115.79 | 1671.21 | 2987.78 |
| Serpina3c | 221.03 | -1.15 | 0.32 | 13.34 | 0.0002593 | 0.0110168 | 195.80 | 189.60 | 546.03 | 269.32 | 137.03 | 72.15 | 170.13 | 188.18 |
| Gpr37 | 23.11 | -1.86 | 0.52 | 13.33 | 0.0002606 | 0.0110168 | 21.02 | 41.18 | 49.75 | 32.96 | 9.56 | 14.06 | 5.32 | 11.07 |
| Cxcl5 | 15.86 | 4.36 | 1.17 | 13.35 | 0.0002587 | 0.0110168 | 0.00 | 7.66 | 0.00 | 0.94 | 13.81 | 91.83 | 7.09 | 5.53 |
| Mrgprx2 | 14.31 | 2.45 | 0.67 | 13.34 | 0.0002604 | 0.0110168 | 7.74 | 4.79 | 1.21 | 7.53 | 12.75 | 18.74 | 23.93 | 37.82 |
| Kmt2e | 4815.48 | 0.34 | 0.09 | 13.25 | 0.000273 | 0.011482 | 4428.24 | 4905.64 | 3533.40 | 4707.40 | 4912.91 | 5337.25 | 4690.20 | 6008.77 |
| Spg11 | 1430.43 | 0.39 | 0.11 | 13.25 | 0.0002726 | 0.011482 | 1298.71 | 1252.50 | 1309.25 | 1049.96 | 1730.41 | 1623.85 | 1618.93 | 1559.84 |
| Spag9 | 4615.61 | 0.31 | 0.09 | 13.23 | 0.0002756 | 0.0115605 | 4471.38 | 4615.49 | 3705.70 | 4103.79 | 4715.34 | 5549.95 | 4272.84 | 5490.36 |
| Tox2 | 363.12 | -0.64 | 0.18 | 13.20 | 0.0002798 | 0.0117065 | 402.67 | 443.36 | 484.14 | 444.47 | 271.94 | 234.25 | 256.09 | 368.05 |
| Me1 | 8175.27 | -0.67 | 0.19 | 13.17 | 0.000284 | 0.0118296 | 9266.89 | 6803.55 | 16522.78 | 6144.39 | 6469.11 | 4863.12 | 9327.23 | 6005.08 |
| Rasa1 | 2520.09 | 0.37 | 0.10 | 13.17 | 0.0002842 | 0.0118296 | 2306.49 | 2584.49 | 1822.52 | 2345.70 | 2739.55 | 3111.84 | 2393.39 | 2856.79 |
| Ncoa6 | 2125.73 | 0.35 | 0.10 | 13.16 | 0.0002863 | 0.0118826 | 2068.65 | 2050.16 | 1669.63 | 1807.06 | 2265.78 | 2511.21 | 2296.81 | 2336.54 |
| Gm11816 | 71.50 | -1.29 | 0.36 | 13.14 | 0.0002886 | 0.0119469 | 90.71 | 110.12 | 92.22 | 119.59 | 37.18 | 27.17 | 23.04 | 71.95 |
| Vps13d | 2764.01 | 0.32 | 0.09 | 13.12 | 0.0002918 | 0.0120502 | 2557.60 | 2482.98 | 2345.49 | 2405.96 | 3103.90 | 2916.94 | 3311.40 | 2987.78 |
| Eif5a | 7023.67 | -0.32 | 0.09 | 13.10 | 0.0002959 | 0.0121875 | 7725.91 | 7111.88 | 9112.58 | 6767.77 | 6600.83 | 6029.71 | 6742.43 | 6098.24 |
| Lrch2 | 1371.50 | 0.44 | 0.12 | 13.07 | 0.0003006 | 0.0123483 | 1206.90 | 1471.79 | 825.11 | 1454.88 | 1334.19 | 1816.88 | 1127.14 | 1735.11 |
| Rock2 | 7553.44 | 0.50 | 0.14 | 13.04 | 0.0003041 | 0.0124625 | 7239.17 | 6135.16 | 6349.69 | 5519.12 | 7530.30 | 7964.65 | 9046.33 | 10643.10 |
| Pcdh7 | 2172.58 | 0.57 | 0.16 | 13.01 | 0.0003096 | 0.0126542 | 1995.64 | 1792.57 | 1696.32 | 1632.85 | 2046.96 | 3340.47 | 2660.11 | 2215.70 |
| Kmt2a | 4742.08 | 0.40 | 0.11 | 12.99 | 0.0003134 | 0.0127776 | 4614.08 | 4583.89 | 3412.06 | 4048.23 | 5540.71 | 5487.17 | 5053.50 | 5197.02 |
| Syngap1 | 908.15 | 0.39 | 0.11 | 12.98 | 0.0003143 | 0.0127794 | 856.22 | 851.28 | 711.05 | 782.53 | 908.23 | 1096.31 | 1041.18 | 1018.37 |
| Bmpr1b | 267.27 | 0.91 | 0.25 | 12.97 | 0.0003173 | 0.0128678 | 234.52 | 210.67 | 140.75 | 208.11 | 240.07 | 598.75 | 252.54 | 252.75 |
| Pitpnm2 | 2707.76 | -0.53 | 0.15 | 12.92 | 0.0003247 | 0.013137 | 2567.56 | 3581.32 | 3657.17 | 2915.41 | 2338.02 | 1888.09 | 2139.96 | 2574.53 |
| Dstyk | 2717.45 | 0.28 | 0.08 | 12.90 | 0.0003284 | 0.0132209 | 2584.15 | 2437.02 | 2412.23 | 2452.10 | 2928.63 | 2969.41 | 2883.41 | 3072.64 |
| Ryr2 | 202.55 | -1.21 | 0.33 | 12.90 | 0.0003283 | 0.0132209 | 198.02 | 259.50 | 288.79 | 383.26 | 155.09 | 63.72 | 92.16 | 179.88 |
| Acat1 | 3630.48 | -0.35 | 0.10 | 12.88 | 0.0003317 | 0.0132846 | 3900.57 | 3930.83 | 4494.41 | 3879.67 | 3306.79 | 3111.84 | 2980.00 | 3439.78 |
| Peg10 | 1586.67 | 0.42 | 0.12 | 12.88 | 0.0003317 | 0.0132846 | 1424.82 | 1185.47 | 1308.04 | 1590.48 | 1680.48 | 1961.18 | 1625.13 | 1917.75 |
| Reep6 | 1519.22 | -0.40 | 0.11 | 12.85 | 0.000337 | 0.0134304 | 1654.92 | 1528.28 | 2022.73 | 1464.29 | 1634.81 | 1021.35 | 1549.81 | 1277.58 |
| Zfp385b | 94.11 | -0.96 | 0.27 | 12.85 | 0.0003368 | 0.0134304 | 98.45 | 160.87 | 107.99 | 138.43 | 62.67 | 58.10 | 46.08 | 80.25 |
| Gm26512 | 46.41 | 1.37 | 0.37 | 12.84 | 0.0003389 | 0.0134714 | 32.08 | 21.07 | 18.20 | 33.90 | 84.98 | 49.66 | 56.71 | 74.72 |
| Sfrp5 | 150.56 | -1.22 | 0.33 | 12.82 | 0.0003432 | 0.0136077 | 140.49 | 246.10 | 254.81 | 158.20 | 101.98 | 38.42 | 163.04 | 101.47 |
| Cox4i1 | 10976.21 | -0.33 | 0.09 | 12.79 | 0.000348 | 0.0137626 | 12224.95 | 11137.51 | 13549.96 | 10959.14 | 10861.52 | 8329.15 | 10915.14 | 9832.28 |
| Pla1a | 648.44 | 1.03 | 0.29 | 12.78 | 0.0003496 | 0.0137954 | 396.03 | 338.98 | 464.73 | 504.73 | 619.29 | 1309.01 | 1070.42 | 484.28 |
| Oprk1 | 119.26 | -1.05 | 0.28 | 12.78 | 0.0003508 | 0.0138055 | 188.06 | 221.20 | 126.19 | 112.06 | 71.17 | 74.96 | 92.16 | 68.26 |
| Skil | 1754.30 | 0.50 | 0.14 | 12.76 | 0.0003538 | 0.013857 | 1610.67 | 1393.27 | 1509.46 | 1405.91 | 1772.90 | 2082.99 | 1634.88 | 2624.34 |
| Nptx2 | 492.13 | 0.93 | 0.26 | 12.76 | 0.0003538 | 0.013857 | 321.91 | 225.03 | 498.70 | 272.14 | 451.46 | 765.54 | 919.79 | 482.44 |
| Prrc2c | 4509.39 | 0.39 | 0.11 | 12.75 | 0.0003557 | 0.0138974 | 4262.30 | 4332.05 | 3424.20 | 3797.75 | 5121.12 | 5536.84 | 4970.21 | 4630.64 |
| Pde10a | 214.86 | 1.04 | 0.29 | 12.74 | 0.0003588 | 0.0139849 | 188.06 | 195.34 | 121.34 | 84.75 | 178.46 | 411.35 | 272.04 | 267.51 |
| Hs2st1 | 1479.69 | 0.47 | 0.13 | 12.73 | 0.00036 | 0.0139952 | 1334.11 | 1317.62 | 1113.90 | 1345.64 | 1389.43 | 2261.96 | 1499.30 | 1575.53 |
| Cpt1c | 696.90 | 0.43 | 0.12 | 12.72 | 0.0003621 | 0.0140444 | 613.96 | 619.55 | 553.31 | 626.21 | 688.34 | 899.54 | 817.88 | 756.40 |
| Ino80 | 1556.04 | 0.39 | 0.11 | 12.71 | 0.0003633 | 0.0140569 | 1410.44 | 1463.17 | 1158.79 | 1422.86 | 1846.19 | 1751.29 | 1707.54 | 1688.06 |
| Wisp2 | 355.38 | 2.58 | 0.71 | 12.71 | 0.0003645 | 0.0140683 | 110.62 | 71.82 | 146.82 | 79.10 | 192.27 | 1277.15 | 688.51 | 276.73 |
| Nktr | 5503.36 | 0.38 | 0.10 | 12.67 | 0.0003709 | 0.014258 | 4910.55 | 5299.20 | 4154.66 | 5102.90 | 6366.08 | 6773.70 | 5716.32 | 5703.44 |
| Tpr | 4678.04 | 0.30 | 0.08 | 12.66 | 0.000373 | 0.014258 | 4522.27 | 4567.62 | 3546.75 | 4560.50 | 4835.37 | 5387.85 | 4789.44 | 5214.55 |
| Atp1a2 | 4198.74 | -0.71 | 0.20 | 12.67 | 0.0003714 | 0.014258 | 3551.00 | 6432.01 | 5201.82 | 5733.82 | 3487.37 | 2728.59 | 2563.53 | 3891.77 |
| Itga2 | 339.23 | 0.73 | 0.20 | 12.67 | 0.0003722 | 0.014258 | 241.16 | 328.45 | 254.81 | 184.57 | 453.58 | 470.38 | 436.85 | 344.07 |
| Rnf5 | 1156.73 | -0.37 | 0.10 | 12.60 | 0.0003848 | 0.014673 | 1318.63 | 1229.52 | 1458.50 | 1118.70 | 1138.73 | 901.41 | 1081.06 | 1007.30 |
| Kcna1 | 467.12 | -0.96 | 0.27 | 12.59 | 0.0003872 | 0.014732 | 361.74 | 845.54 | 522.97 | 788.18 | 281.50 | 257.68 | 237.48 | 441.85 |
| Chd1 | 1603.28 | 0.32 | 0.09 | 12.58 | 0.0003895 | 0.0147813 | 1511.11 | 1523.50 | 1333.52 | 1425.69 | 1608.25 | 1764.40 | 1699.57 | 1960.18 |
| Pla2g16 | 2157.18 | -0.49 | 0.14 | 12.56 | 0.0003934 | 0.0148968 | 2140.55 | 2090.38 | 2708.29 | 2954.01 | 1891.87 | 1631.35 | 2116.92 | 1724.04 |
| Cilp | 1026.69 | 2.32 | 0.64 | 12.55 | 0.0003952 | 0.0149258 | 439.17 | 263.33 | 377.37 | 318.28 | 559.81 | 3450.10 | 1994.64 | 810.82 |
| Slc22a18 | 736.27 | -0.39 | 0.11 | 12.54 | 0.0003977 | 0.0149815 | 808.65 | 699.03 | 1016.82 | 710.02 | 737.20 | 598.75 | 772.69 | 547.01 |
| Gabra1 | 37.05 | -1.75 | 0.45 | 12.54 | 0.0003985 | 0.0149815 | 77.44 | 63.20 | 24.27 | 71.57 | 22.31 | 13.12 | 8.86 | 15.68 |
| Lpgat1 | 4157.54 | 0.31 | 0.09 | 12.48 | 0.0004106 | 0.0152921 | 3937.07 | 3697.18 | 3670.51 | 3665.92 | 4493.33 | 5078.63 | 4259.55 | 4458.15 |
| Pomp | 3045.49 | -0.29 | 0.08 | 12.48 | 0.0004105 | 0.0152921 | 3434.84 | 3226.06 | 3401.14 | 3198.85 | 2903.13 | 2611.47 | 3008.35 | 2580.06 |
| Tmf1 | 1314.21 | 0.38 | 0.11 | 12.49 | 0.0004096 | 0.0152921 | 1297.61 | 1144.30 | 931.89 | 1336.23 | 1358.62 | 1487.05 | 1333.60 | 1624.42 |
| Dlgap1 | 94.48 | -0.95 | 0.27 | 12.49 | 0.00041 | 0.0152921 | 120.58 | 126.40 | 88.58 | 178.92 | 49.93 | 54.35 | 53.17 | 83.94 |
| Omp | 148.51 | -0.75 | 0.21 | 12.48 | 0.0004121 | 0.0153134 | 119.47 | 176.19 | 248.75 | 173.27 | 138.09 | 83.39 | 139.12 | 109.77 |
| 2810474O19R | 1806.93 | 0.41 | 0.11 | 12.46 | 0.0004155 | 0.0153745 | 1756.69 | 1858.65 | 1174.56 | 1693.12 | 1874.87 | 2324.74 | 1777.54 | 1995.24 |
| Timp1 | 326.81 | 1.27 | 0.36 | 12.46 | 0.0004157 | 0.0153745 | 302.00 | 159.91 | 185.65 | 146.90 | 182.71 | 768.35 | 516.60 | 352.37 |
| Thsd7a | 442.70 | 0.52 | 0.15 | 12.44 | 0.0004193 | 0.0154727 | 450.24 | 353.34 | 305.78 | 387.03 | 506.69 | 594.07 | 452.80 | 491.66 |
| Iah1 | 1659.49 | -0.37 | 0.10 | 12.44 | 0.0004207 | 0.0154882 | 1949.18 | 1725.54 | 1909.88 | 1736.44 | 1701.73 | 1319.32 | 1631.33 | 1302.48 |
| Prss32 | 112.06 | 1.70 | 0.49 | 12.43 | 0.0004221 | 0.0155023 | 29.87 | 50.75 | 83.72 | 47.08 | 142.34 | 352.32 | 126.71 | 63.65 |
| Ak5 | 137.82 | 1.38 | 0.39 | 12.41 | 0.0004266 | 0.0156311 | 107.30 | 94.80 | 55.82 | 71.57 | 141.28 | 360.75 | 184.31 | 86.71 |
| Pdgfrb | 2693.03 | 0.53 | 0.15 | 12.39 | 0.0004313 | 0.0157681 | 1794.30 | 2212.95 | 2209.59 | 2701.65 | 2584.46 | 3713.40 | 3197.10 | 3130.76 |
| C3 | 23308.72 | -0.63 | 0.18 | 12.33 | 0.0004446 | 0.0162181 | 36984.58 | 20867.40 | 28913.95 | 25099.23 | 17197.86 | 16928.16 | 23016.79 | 17461.77 |
| Colq | 48.29 | -1.34 | 0.39 | 12.32 | 0.0004472 | 0.0162743 | 48.67 | 86.18 | 82.51 | 61.21 | 15.93 | 28.11 | 22.15 | 41.51 |
| Fndc3b | 4401.38 | 0.39 | 0.11 | 12.31 | 0.0004493 | 0.0163152 | 4098.58 | 4092.66 | 3331.98 | 4098.14 | 4164.03 | 5952.87 | 4853.24 | 4619.57 |
| Slit3 | 4827.73 | 0.53 | 0.15 | 12.27 | 0.0004603 | 0.016652 | 4329.78 | 3546.84 | 4220.18 | 3767.62 | 4125.79 | 6067.19 | 6416.35 | 6148.06 |
| Nov | 703.17 | 1.60 | 0.46 | 12.27 | 0.0004607 | 0.016652 | 258.86 | 518.05 | 328.83 | 336.18 | 697.90 | 2135.46 | 615.85 | 734.26 |
| Pkig | 1410.96 | -0.49 | 0.14 | 12.24 | 0.0004668 | 0.0168318 | 1590.76 | 1587.65 | 2071.26 | 1291.03 | 1120.68 | 1189.08 | 1190.05 | 1247.14 |
| Tnfaip8l3 | 171.10 | 1.26 | 0.37 | 12.24 | 0.0004678 | 0.0168318 | 101.77 | 99.59 | 105.57 | 106.41 | 148.72 | 495.68 | 175.45 | 135.60 |
| Vcl | 17700.99 | 0.43 | 0.12 | 12.23 | 0.0004707 | 0.0169003 | 15566.87 | 16392.67 | 14843.44 | 14670.25 | 15662.90 | 19981.90 | 19897.67 | 24592.22 |
| Chd8 | 4240.03 | 0.30 | 0.09 | 12.22 | 0.0004737 | 0.0169483 | 3952.56 | 4003.61 | 3708.13 | 3653.67 | 4625.04 | 5032.72 | 4442.09 | 4502.42 |
| Ndufa13 | 2740.74 | -0.38 | 0.11 | 12.21 | 0.0004742 | 0.0169483 | 3008.94 | 3031.67 | 3603.78 | 2467.17 | 2555.78 | 2103.60 | 2825.81 | 2329.16 |
| Erbb2ip | 5860.32 | 0.41 | 0.12 | 12.19 | 0.0004795 | 0.0170877 | 6141.79 | 5213.02 | 3971.44 | 5504.99 | 6515.85 | 7380.89 | 5747.33 | 6407.26 |
| Ubb | 3456.47 | -0.30 | 0.09 | 12.18 | 0.0004833 | 0.0170877 | 4004.55 | 3533.44 | 4074.57 | 3484.17 | 3263.24 | 2909.44 | 3353.94 | 3028.37 |
| Ndufv3 | 2095.55 | -0.33 | 0.09 | 12.18 | 0.0004835 | 0.0170877 | 2309.81 | 2272.32 | 2459.55 | 2177.14 | 2049.08 | 1615.42 | 1969.83 | 1911.29 |
| 1500011K16R | 614.34 | -0.51 | 0.15 | 12.18 | 0.0004824 | 0.0170877 | 815.29 | 643.49 | 779.00 | 581.95 | 556.62 | 461.01 | 630.91 | 446.46 |
| Ankfn1 | 124.72 | 1.16 | 0.32 | 12.18 | 0.0004825 | 0.0170877 | 79.65 | 89.05 | 61.88 | 56.50 | 293.18 | 149.92 | 155.07 | 112.54 |
| Kazald1 | 475.76 | -0.50 | 0.14 | 12.16 | 0.0004873 | 0.0171726 | 496.70 | 514.22 | 651.59 | 469.89 | 567.24 | 357.94 | 439.51 | 309.02 |
| Tmem200a | 244.08 | 1.12 | 0.32 | 12.16 | 0.000488 | 0.0171726 | 133.85 | 203.96 | 110.42 | 215.64 | 130.66 | 483.50 | 374.83 | 299.79 |
| Siglech | 44.81 | -1.14 | 0.33 | 12.15 | 0.0004906 | 0.0172265 | 58.63 | 59.37 | 75.23 | 52.73 | 30.81 | 28.11 | 20.38 | 33.21 |
| Gbe1 | 1656.88 | -0.45 | 0.13 | 12.14 | 0.0004946 | 0.0172892 | 1798.73 | 1747.57 | 2386.74 | 1595.19 | 1517.96 | 1201.26 | 1452.34 | 1555.23 |
| Ap4e1 | 718.00 | 0.55 | 0.16 | 12.14 | 0.0004942 | 0.0172892 | 694.71 | 696.15 | 464.73 | 536.75 | 914.60 | 810.52 | 736.36 | 890.15 |
| Prox1 | 208.40 | 1.02 | 0.30 | 12.12 | 0.0004977 | 0.0173597 | 121.69 | 93.84 | 205.06 | 124.30 | 223.07 | 410.41 | 293.30 | 195.56 |
| Paqr9 | 64.41 | -1.74 | 0.51 | 12.11 | 0.0005016 | 0.0174582 | 45.36 | 28.73 | 280.29 | 30.13 | 32.93 | 3.75 | 68.23 | 25.83 |
| Ltbp2 | 599.23 | 2.62 | 0.74 | 12.09 | 0.0005067 | 0.0175968 | 152.66 | 148.42 | 219.62 | 149.73 | 338.86 | 2414.69 | 1011.94 | 357.91 |
| Tjp1 | 8952.68 | 0.36 | 0.10 | 12.09 | 0.0005082 | 0.0176095 | 8392.97 | 8913.08 | 7193.00 | 7191.52 | 10227.36 | 10548.94 | 9469.01 | 9685.61 |
| Fcer1a | 22.45 | 2.71 | 0.72 | 12.07 | 0.0005121 | 0.0177075 | 11.06 | 7.66 | 1.21 | 6.59 | 11.68 | 32.80 | 73.55 | 35.05 |
| Slc2a5 | 64.46 | -2.32 | 0.66 | 12.06 | 0.0005143 | 0.0177433 | 57.52 | 21.07 | 287.57 | 11.30 | 12.75 | 4.69 | 114.31 | 6.46 |

| | | | | | | | | | | | | | |
|---|---|---|---|---|---|---|---|---|---|---|---|---|---|
| Dhx33 | 1570.59 | 0.35 | 0.10 | 12.04 | 0.0005205 | 0.0178823 | 1463.54 | 1502.43 | 1208.54 | 1424.74 | 1782.46 | 1874.97 | 1638.42 | 1669.61 |
| Gm11266 | 66.42 | -0.92 | 0.26 | 12.04 | 0.0005199 | 0.0178823 | 82.97 | 104.38 | 66.74 | 101.70 | 42.49 | 51.54 | 36.33 | 45.20 |
| Mrpl28 | 1021.88 | -0.38 | 0.11 | 12.02 | 0.0005257 | 0.0180213 | 1109.55 | 1122.27 | 1361.43 | 968.98 | 928.41 | 837.69 | 944.60 | 902.15 |
| Mical2 | 3216.87 | 0.58 | 0.17 | 12.01 | 0.0005299 | 0.0181081 | 2887.26 | 2181.35 | 3119.63 | 2060.37 | 3086.90 | 3830.53 | 4356.13 | 4212.78 |
| Igfals | 83.57 | -1.42 | 0.42 | 12.01 | 0.0005305 | 0.0181081 | 101.77 | 66.07 | 243.89 | 62.15 | 38.24 | 17.80 | 83.29 | 55.35 |
| Clu | 7408.15 | -0.56 | 0.16 | 11.95 | 0.0005464 | 0.0186115 | 11324.47 | 7007.51 | 9408.65 | 7642.58 | 5286.83 | 5881.66 | 5951.14 | 6762.40 |
| C1galt1 | 777.95 | 0.50 | 0.14 | 11.94 | 0.000548 | 0.0186265 | 668.16 | 643.49 | 661.30 | 589.48 | 1106.87 | 923.90 | 754.97 | 875.39 |
| Sphk2 | 1558.91 | -0.32 | 0.09 | 11.93 | 0.0005511 | 0.0186912 | 1466.86 | 1569.46 | 2096.74 | 1618.73 | 1607.19 | 1250.92 | 1531.20 | 1330.16 |
| Vwa1 | 1701.66 | 0.67 | 0.20 | 11.93 | 0.0005524 | 0.0186958 | 1181.45 | 1375.07 | 1379.63 | 1339.05 | 1383.05 | 2615.22 | 2527.19 | 1812.59 |
| Pqlc3 | 799.16 | 0.46 | 0.13 | 11.92 | 0.0005565 | 0.0187944 | 881.67 | 725.84 | 538.75 | 683.65 | 679.84 | 1206.88 | 684.08 | 992.54 |
| Ceacam19 | 107.04 | -0.87 | 0.25 | 11.91 | 0.0005597 | 0.0188627 | 164.83 | 114.91 | 158.95 | 106.41 | 98.79 | 76.84 | 67.34 | 68.26 |
| Cep192 | 783.52 | 0.40 | 0.11 | 11.88 | 0.0005665 | 0.01905 | 761.09 | 724.88 | 563.01 | 734.50 | 778.63 | 1010.10 | 843.58 | 852.33 |
| Emc6 | 1072.63 | -0.39 | 0.11 | 11.87 | 0.0005693 | 0.0191047 | 1266.63 | 1071.52 | 1397.83 | 1081.98 | 977.27 | 912.66 | 930.42 | 942.73 |
| Cox7a2 | 2813.45 | -0.47 | 0.14 | 11.86 | 0.0005728 | 0.0191805 | 3380.64 | 3420.45 | 3560.10 | 2615.96 | 2426.18 | 1991.16 | 2429.72 | 2683.38 |
| Atp5g3 | 7209.73 | -0.32 | 0.09 | 11.85 | 0.0005755 | 0.0191904 | 7941.62 | 7935.39 | 8840.78 | 6892.07 | 6704.93 | 5925.70 | 6858.52 | 6578.84 |
| Apaf1 | 659.55 | 0.51 | 0.15 | 11.86 | 0.0005744 | 0.0191904 | 547.58 | 541.03 | 492.64 | 654.46 | 642.66 | 1004.48 | 682.31 | 711.20 |
| Laptm4b | 4254.52 | -0.33 | 0.10 | 11.83 | 0.0005821 | 0.019366 | 4383.99 | 4653.80 | 5376.55 | 4245.98 | 4364.79 | 3459.47 | 3745.60 | 3805.98 |
| Cend1 | 142.78 | 0.77 | 0.23 | 11.83 | 0.0005832 | 0.019366 | 120.58 | 94.80 | 114.06 | 101.70 | 128.53 | 181.78 | 168.36 | 232.45 |
| Cox5a | 2975.08 | -0.35 | 0.10 | 11.79 | 0.0005949 | 0.0197142 | 3423.78 | 3145.62 | 3777.29 | 2821.24 | 2733.17 | 2504.65 | 2734.55 | 2660.31 |
| Yes1 | 1588.58 | 0.52 | 0.15 | 11.77 | 0.0006022 | 0.0199161 | 1438.10 | 1317.62 | 1168.50 | 1424.74 | 1931.17 | 2346.29 | 1498.42 | 1583.83 |
| Ptch1 | 1584.77 | 0.59 | 0.17 | 11.76 | 0.0006046 | 0.0199528 | 1404.91 | 1518.71 | 1113.90 | 1125.29 | 1705.98 | 2379.09 | 1940.59 | 1489.74 |
| Lpin2 | 2149.11 | 0.47 | 0.13 | 11.74 | 0.0006131 | 0.0200665 | 1827.49 | 1534.99 | 1741.22 | 2018.94 | 2414.50 | 2451.24 | 3011.90 | 2192.64 |
| Slc5a1 | 297.88 | -0.93 | 0.28 | 11.74 | 0.0006129 | 0.0200665 | 342.93 | 441.44 | 410.13 | 382.32 | 223.07 | 148.05 | 120.51 | 314.55 |
| Gm4980 | 176.88 | -0.79 | 0.23 | 11.74 | 0.0006117 | 0.0200665 | 231.20 | 239.39 | 269.37 | 156.32 | 98.79 | 139.62 | 137.35 | 142.98 |
| Nox4 | 157.06 | 1.53 | 0.44 | 11.75 | 0.0006095 | 0.0200665 | 99.56 | 58.41 | 53.39 | 147.84 | 84.98 | 495.68 | 197.60 | 118.99 |
| Kif13b | 1443.89 | 0.45 | 0.13 | 11.71 | 0.00062 | 0.020253 | 1331.90 | 1463.17 | 998.62 | 1162.96 | 1717.66 | 1625.73 | 1632.22 | 1618.88 |
| Ifitm3 | 3944.85 | -0.49 | 0.14 | 11.70 | 0.0006243 | 0.020351 | 4145.04 | 4378.97 | 4676.42 | 5343.97 | 2533.47 | 3702.16 | 3560.40 | 3218.39 |
| Pxmp4 | 1307.92 | -0.31 | 0.09 | 11.68 | 0.0006315 | 0.0205437 | 1413.76 | 1369.33 | 1527.66 | 1425.69 | 1158.92 | 1130.04 | 1283.09 | 1154.89 |
| Slco2b1 | 1254.52 | -0.47 | 0.14 | 11.66 | 0.0006396 | 0.0207628 | 1383.89 | 1425.82 | 1382.05 | 1709.13 | 828.56 | 1001.67 | 1105.87 | 1199.17 |
| Gyg | 1477.61 | -0.33 | 0.10 | 11.64 | 0.0006439 | 0.0208613 | 1623.95 | 1624.04 | 1698.75 | 1668.64 | 1191.85 | 1354.93 | 1230.81 | 1427.94 |
| 2410089E03R | 817.07 | 0.38 | 0.11 | 11.63 | 0.0006502 | 0.0210215 | 709.09 | 770.85 | 616.40 | 785.35 | 879.54 | 943.58 | 928.65 | 903.07 |
| Nop10 | 1151.15 | -0.34 | 0.10 | 11.62 | 0.0006534 | 0.0210827 | 1317.52 | 1134.72 | 1453.65 | 1150.72 | 1081.37 | 966.07 | 1144.86 | 960.26 |
| Fam179b | 1394.22 | 0.31 | 0.09 | 11.61 | 0.0006559 | 0.0211203 | 1333.01 | 1322.41 | 1099.33 | 1312.69 | 1522.21 | 1559.20 | 1381.45 | 1623.49 |
| Fkbp4 | 5933.28 | -0.35 | 0.10 | 11.59 | 0.0006621 | 0.0212777 | 5992.45 | 5925.45 | 8663.63 | 5351.50 | 6081.39 | 4677.59 | 5666.69 | 5107.54 |
| Cd209g | 84.32 | -1.68 | 0.50 | 11.56 | 0.0006724 | 0.0215644 | 105.09 | 140.76 | 158.95 | 111.12 | 10.62 | 20.61 | 43.42 | 83.94 |
| Ep400 | 3369.53 | 0.28 | 0.08 | 11.54 | 0.0006799 | 0.0217623 | 3146.12 | 3146.58 | 2853.90 | 3060.42 | 3775.24 | 3699.35 | 3728.76 | 3545.86 |
| Rdh10 | 1195.57 | 0.46 | 0.13 | 11.52 | 0.0006873 | 0.0219547 | 1061.98 | 962.36 | 901.55 | 1114.94 | 1371.37 | 1492.67 | 1500.19 | 1159.51 |
| Bend5 | 364.47 | -0.55 | 0.16 | 11.51 | 0.0006911 | 0.022032 | 423.69 | 506.56 | 430.75 | 376.67 | 319.74 | 281.11 | 251.66 | 325.62 |
| Nid1 | 10116.88 | 0.61 | 0.18 | 11.49 | 0.0006982 | 0.0221406 | 7238.06 | 9093.10 | 7412.62 | 9311.22 | 7814.99 | 16749.19 | 12187.60 | 11128.30 |
| Pxdn | 4296.41 | 0.51 | 0.15 | 11.50 | 0.0006961 | 0.0221406 | 3216.92 | 3642.60 | 3431.48 | 4176.30 | 3775.24 | 6539.45 | 4870.08 | 4719.20 |
| Pcnx | 3195.69 | 0.28 | 0.08 | 11.49 | 0.0006987 | 0.0221406 | 2942.57 | 3146.58 | 2761.68 | 2866.44 | 3152.76 | 3654.37 | 3371.66 | 3669.46 |
| Frzb | 435.98 | 2.11 | 0.61 | 11.47 | 0.0007084 | 0.0224001 | 172.57 | 207.79 | 116.49 | 225.06 | 271.94 | 1623.85 | 599.90 | 270.27 |
| Rel | 73.27 | 0.95 | 0.29 | 11.46 | 0.0007097 | 0.0224001 | 54.21 | 45.01 | 61.88 | 39.55 | 100.91 | 103.07 | 71.78 | 109.77 |
| Vgll3 | 977.55 | 0.55 | 0.16 | 11.46 | 0.000713 | 0.0224609 | 807.55 | 720.09 | 729.25 | 890.82 | 1146.17 | 1277.15 | 1361.96 | 887.39 |
| Mdn1 | 1380.39 | 0.37 | 0.11 | 11.42 | 0.000728 | 0.0228447 | 1244.51 | 1331.98 | 1087.20 | 1181.79 | 1668.80 | 1577.00 | 1484.24 | 1467.60 |
| Tst | 792.79 | -0.40 | 0.12 | 11.42 | 0.0007271 | 0.0228447 | 868.39 | 767.97 | 1043.52 | 879.52 | 795.63 | 642.79 | 661.93 | 682.60 |
| F13a1 | 2510.87 | -0.74 | 0.22 | 11.41 | 0.0007298 | 0.0228541 | 3593.03 | 3515.24 | 2561.47 | 3244.05 | 1033.57 | 1471.12 | 1956.54 | 2711.97 |
| Cox5b | 1271.21 | -0.43 | 0.13 | 11.40 | 0.0007332 | 0.022915 | 1412.66 | 1449.76 | 1667.20 | 1207.22 | 1103.68 | 916.40 | 1246.76 | 1165.96 |
| Nipbl | 2592.40 | 0.31 | 0.09 | 11.39 | 0.0007376 | 0.0230018 | 2306.49 | 2587.36 | 2087.04 | 2437.98 | 2851.08 | 2885.08 | 2593.65 | 2990.55 |
| Car4 | 30.60 | -1.96 | 0.56 | 11.39 | 0.0007388 | 0.0230018 | 32.08 | 36.39 | 47.32 | 77.22 | 7.44 | 4.69 | 22.15 | 17.53 |
| Zmat3 | 2156.75 | 0.42 | 0.13 | 11.38 | 0.0007408 | 0.0230196 | 1827.49 | 1594.36 | 1900.17 | 2109.34 | 2105.38 | 2755.77 | 2437.70 | 2523.79 |
| Dkk3 | 6190.05 | 0.39 | 0.12 | 11.37 | 0.0007467 | 0.023155 | 5007.90 | 5304.95 | 5554.91 | 5722.52 | 7123.46 | 8106.14 | 5886.45 | 6814.06 |
| Plch1 | 32.84 | 1.63 | 0.46 | 11.36 | 0.0007485 | 0.0231671 | 18.81 | 15.32 | 8.49 | 19.78 | 52.05 | 49.66 | 70.00 | 28.60 |
| Timm13 | 1170.99 | -0.40 | 0.12 | 11.35 | 0.0007529 | 0.023257 | 1258.89 | 1185.47 | 1658.71 | 1035.84 | 1169.54 | 985.74 | 1257.39 | 816.36 |
| Hook3 | 2852.91 | 0.33 | 0.10 | 11.32 | 0.0007676 | 0.0236652 | 2699.20 | 2957.94 | 2175.61 | 2537.80 | 3050.79 | 3421.05 | 2771.76 | 3209.17 |
| Pole3 | 1220.16 | -0.35 | 0.10 | 11.31 | 0.0007708 | 0.023718 | 1408.23 | 1405.71 | 1425.74 | 1217.58 | 1066.50 | 1011.04 | 1086.37 | 1140.13 |
| 4930402H24R | 7103.54 | 0.32 | 0.09 | 11.30 | 0.0007763 | 0.0238017 | 7040.05 | 6522.02 | 5826.71 | 5954.17 | 8024.25 | 7700.41 | 8058.31 | 7702.37 |
| Bmp1 | 3771.84 | 0.56 | 0.17 | 11.29 | 0.000778 | 0.0238017 | 3099.66 | 3533.44 | 2646.41 | 3300.55 | 3328.03 | 6002.53 | 4567.91 | 3696.21 |
| Nfasc | 622.07 | -0.79 | 0.24 | 11.30 | 0.000777 | 0.0238017 | 582.98 | 1067.69 | 694.06 | 877.64 | 356.92 | 373.87 | 326.98 | 696.44 |
| Fut8 | 1145.38 | 0.49 | 0.15 | 11.28 | 0.0007841 | 0.0238977 | 1009.99 | 1040.88 | 853.02 | 1007.59 | 1046.32 | 1702.56 | 1325.63 | 1177.03 |
| Clca1 | 1023.03 | 0.77 | 0.22 | 11.28 | 0.0007833 | 0.0238977 | 961.31 | 588.91 | 578.79 | 933.19 | 1494.59 | 1467.37 | 1244.10 | 915.98 |
| Mau2 | 3242.18 | 0.28 | 0.08 | 11.27 | 0.0007875 | 0.0239557 | 2980.18 | 2891.87 | 2822.35 | 3038.76 | 3537.30 | 3611.27 | 3690.66 | 3365.06 |
| Banf1 | 1641.02 | -0.41 | 0.12 | 11.25 | 0.0007949 | 0.0241333 | 1785.45 | 1971.64 | 2133.15 | 1494.43 | 1493.53 | 1252.79 | 1525.00 | 1472.21 |
| Cntfr | 1343.14 | -0.87 | 0.26 | 11.24 | 0.0007986 | 0.024201 | 1390.53 | 2361.37 | 1790.97 | 1455.82 | 950.72 | 645.61 | 677.88 | 1472.21 |
| Eda2r | 311.65 | 0.67 | 0.20 | 11.22 | 0.0008076 | 0.0244257 | 234.52 | 232.69 | 263.31 | 251.43 | 291.06 | 437.59 | 261.40 | 521.18 |
| Txn2 | 3907.82 | -0.31 | 0.09 | 11.21 | 0.0008141 | 0.0245775 | 4452.57 | 4168.31 | 4740.73 | 3788.33 | 3482.06 | 3322.66 | 3795.22 | 3512.65 |
| Mir22hg | 1581.33 | 0.53 | 0.16 | 11.20 | 0.0008169 | 0.024616 | 1797.62 | 1264.95 | 957.37 | 1392.73 | 1519.02 | 1998.66 | 1864.38 | 1855.95 |
| Sfi1 | 995.03 | 0.34 | 0.10 | 11.19 | 0.0008233 | 0.0247614 | 846.27 | 933.63 | 872.43 | 828.67 | 1191.85 | 1061.64 | 1172.33 | 1053.43 |
| Sbno1 | 4696.29 | 0.31 | 0.09 | 11.18 | 0.0008265 | 0.0248095 | 4469.17 | 4539.85 | 3807.63 | 4135.81 | 5435.54 | 5281.03 | 4684.88 | 5216.39 |
| Bscl2 | 2124.61 | -0.37 | 0.11 | 11.17 | 0.0008309 | 0.0248966 | 2121.75 | 2328.81 | 2903.65 | 2059.43 | 1904.62 | 1939.63 | 2084.14 | 1654.86 |
| Kat6a | 2767.74 | 0.31 | 0.09 | 11.15 | 0.0008401 | 0.025077 | 2424.86 | 2775.04 | 2350.34 | 2448.34 | 2988.11 | 3310.48 | 2831.13 | 3013.61 |
| Igf2 | 686.51 | -0.76 | 0.23 | 11.15 | 0.0008393 | 0.025077 | 663.74 | 939.38 | 905.19 | 948.26 | 485.45 | 330.77 | 499.77 | 719.50 |
| Myo1d | 3199.74 | 0.36 | 0.11 | 11.13 | 0.0008507 | 0.0252982 | 2873.98 | 2651.52 | 3004.36 | 2699.76 | 3125.14 | 3565.35 | 3794.34 | 3883.47 |
| 1500009L16R | 141.31 | 1.19 | 0.36 | 11.13 | 0.0008494 | 0.0252982 | 119.47 | 93.84 | 80.08 | 62.15 | 78.61 | 199.58 | 258.75 | 237.99 |
| Pcdhb7 | 78.63 | 0.94 | 0.29 | 11.11 | 0.0008587 | 0.0254891 | 49.78 | 59.37 | 49.75 | 64.03 | 76.48 | 149.92 | 93.93 | 85.79 |
| Tro | 160.61 | 1.02 | 0.31 | 11.10 | 0.0008637 | 0.0255888 | 92.92 | 120.65 | 114.06 | 103.58 | 142.34 | 350.44 | 219.76 | 141.13 |
| Sync | 246.14 | 1.16 | 0.35 | 11.09 | 0.0008661 | 0.025613 | 215.71 | 153.21 | 151.67 | 107.35 | 118.97 | 433.84 | 413.81 | 374.51 |
| mt-Rnr2 | 21500.23 | -0.69 | 0.21 | 11.08 | 0.0008719 | 0.0257362 | 24477.55 | 27866.29 | 27175.16 | 27808.41 | 15437.71 | 12502.62 | 11345.79 | 25388.29 |
| Atp2a3 | 1836.71 | -0.70 | 0.21 | 11.06 | 0.0008801 | 0.02593 | 1913.78 | 2439.89 | 2426.79 | 2314.62 | 1369.24 | 926.71 | 1296.38 | 2006.30 |
| Hif1a | 6281.32 | 0.37 | 0.11 | 11.06 | 0.0008835 | 0.0259661 | 6117.45 | 5760.75 | 5456.63 | 4962.59 | 5847.70 | 7761.32 | 6341.03 | 8003.08 |
| Lgi1 | 29.81 | -1.45 | 0.43 | 11.05 | 0.0008845 | 0.0259661 | 30.97 | 44.05 | 47.32 | 47.08 | 13.81 | 13.12 | 29.24 | 12.91 |
| Fam19a5 | 372.13 | 0.55 | 0.17 | 11.04 | 0.0008929 | 0.0261633 | 266.60 | 332.28 | 322.76 | 282.50 | 372.85 | 399.17 | 493.57 | 507.34 |
| Upk3a | 29594.34 | -0.41 | 0.12 | 11.03 | 0.0008965 | 0.0262048 | 34476.76 | 31072.24 | 36236.78 | 28674.75 | 34961.89 | 21779.10 | 27141.65 | 22411.58 |
| Anxa8 | 2576.62 | 0.51 | 0.15 | 11.02 | 0.0008992 | 0.0262048 | 2532.16 | 2047.29 | 1976.62 | 1858.85 | 3601.03 | 3151.19 | 2999.49 | 2446.31 |
| Chchd2 | 2373.60 | -0.39 | 0.12 | 11.03 | 0.0008979 | 0.0262048 | 2687.03 | 2627.58 | 3086.87 | 2292.02 | 2193.55 | 2013.65 | 1893.62 | 2194.48 |
| Krt80 | 180.21 | -0.63 | 0.19 | 11.01 | 0.0009038 | 0.0262901 | 206.86 | 168.53 | 241.47 | 240.13 | 166.77 | 119.94 | 156.84 | 141.13 |
| Srrm2 | 33335.73 | 0.42 | 0.12 | 11.01 | 0.0009056 | 0.0262943 | 30634.82 | 33159.74 | 21586.27 | 31938.57 | 38746.70 | 38017.61 | 36165.82 | 36436.34 |
| Clasp1 | 2652.18 | 0.35 | 0.10 | 11.00 | 0.0009099 | 0.0263708 | 2453.62 | 2678.33 | 2106.45 | 2253.41 | 2852.15 | 2799.81 | 2803.66 | 3270.05 |
| Bmyc | 565.14 | -0.41 | 0.12 | 11.00 | 0.0009133 | 0.0263901 | 543.16 | 637.74 | 770.50 | 534.87 | 588.49 | 391.67 | 599.90 | 454.76 |
| Ada | 190.42 | -0.59 | 0.18 | 10.99 | 0.0009139 | 0.0263901 | 222.35 | 221.20 | 273.01 | 194.93 | 144.47 | 143.36 | 151.53 | 172.50 |
| Tmeff1 | 571.34 | 0.49 | 0.15 | 10.99 | 0.000917 | 0.0264308 | 502.23 | 423.25 | 455.02 | 531.10 | 658.60 | 773.04 | 660.15 | 567.30 |
| Enho | 3121.32 | -0.43 | 0.13 | 10.98 | 0.0009192 | 0.0264488 | 3661.62 | 3607.17 | 4006.62 | 2884.33 | 2633.32 | 2245.09 | 3033.17 | 2899.23 |
| Adcy7 | 1818.85 | 0.78 | 0.24 | 10.98 | 0.0009231 | 0.0265135 | 1605.14 | 1252.50 | 1185.49 | 1531.15 | 1240.71 | 3574.72 | 2341.11 | 1819.97 |

| | | | | | | | | | | | | | |
|---|---|---|---|---|---|---|---|---|---|---|---|---|---|
| Rpl39 | 3063.77 | -0.30 | 0.09 | 10.96 | 0.0009325 | 0.0267341 | 3669.36 | 3352.46 | 3311.35 | 3149.88 | 2850.02 | 2652.70 | 2883.41 | 2640.94 |
| Ormdl2 | 775.44 | -0.44 | 0.13 | 10.95 | 0.0009358 | 0.0267598 | 918.17 | 762.23 | 1087.20 | 742.03 | 725.52 | 697.14 | 695.60 | 575.60 |
| Loxl4 | 452.21 | 0.75 | 0.23 | 10.95 | 0.0009384 | 0.0267598 | 471.25 | 336.11 | 306.99 | 273.08 | 344.17 | 644.67 | 634.46 | 606.96 |
| Lmtk3 | 120.82 | 0.71 | 0.21 | 10.95 | 0.000937 | 0.0267598 | 86.29 | 113.95 | 77.66 | 92.28 | 158.28 | 148.05 | 147.98 | 142.06 |
| Pdzrn4 | 30.02 | -1.53 | 0.46 | 10.94 | 0.0009412 | 0.0267917 | 37.61 | 69.90 | 29.12 | 47.08 | 14.87 | 22.49 | 6.20 | 12.91 |
| Kcnh1 | 275.48 | -1.08 | 0.32 | 10.93 | 0.0009454 | 0.0268619 | 262.18 | 583.16 | 304.56 | 377.61 | 200.77 | 118.06 | 100.13 | 257.36 |
| Ndufa12 | 711.12 | -0.44 | 0.13 | 10.92 | 0.0009504 | 0.0269575 | 819.72 | 790.95 | 899.12 | 744.86 | 584.24 | 509.74 | 651.29 | 689.06 |
| Slc7a11 | 144.14 | 0.96 | 0.29 | 10.91 | 0.000954 | 0.0270106 | 105.09 | 152.25 | 71.59 | 87.58 | 198.64 | 261.43 | 116.97 | 159.58 |
| Runx1 | 1867.16 | 0.51 | 0.15 | 10.89 | 0.0009653 | 0.0272339 | 1724.61 | 1438.27 | 1254.65 | 1903.11 | 2061.83 | 2572.11 | 2147.94 | 1834.73 |
| Fam171b | 420.53 | 0.50 | 0.15 | 10.90 | 0.0009642 | 0.0272339 | 344.04 | 406.01 | 320.34 | 351.24 | 464.20 | 601.57 | 437.74 | 439.08 |
| Med13l | 2678.12 | 0.34 | 0.10 | 10.87 | 0.000977 | 0.0275143 | 2244.54 | 2638.11 | 2127.08 | 2687.52 | 2758.67 | 3035.00 | 2542.26 | 3391.81 |
| Olfr78 | 73.56 | 1.83 | 0.56 | 10.86 | 0.0009824 | 0.0276186 | 25.44 | 27.77 | 42.47 | 33.90 | 64.80 | 281.11 | 78.86 | 34.13 |
| Astn2 | 264.33 | -0.82 | 0.25 | 10.85 | 0.0009894 | 0.0277664 | 234.52 | 384.94 | 334.90 | 411.51 | 149.78 | 157.42 | 163.93 | 277.65 |
| Hunk | 323.59 | -0.52 | 0.16 | 10.83 | 0.0009967 | 0.0278737 | 327.44 | 340.90 | 412.55 | 429.40 | 261.31 | 238.94 | 296.85 | 281.34 |
| 4930578C19R | 116.62 | 0.88 | 0.27 | 10.84 | 0.0009959 | 0.0278737 | 74.12 | 105.33 | 78.87 | 76.28 | 124.28 | 210.83 | 148.87 | 114.38 |
| Ndufa7 | 1997.20 | -0.34 | 0.10 | 10.83 | 0.0010002 | 0.0279204 | 2182.59 | 2146.88 | 2613.65 | 1844.73 | 1866.38 | 1666.02 | 1891.85 | 1765.55 |
| Wdr44 | 791.16 | 0.37 | 0.11 | 10.81 | 0.0010094 | 0.0281296 | 660.42 | 748.82 | 652.81 | 753.33 | 815.81 | 1019.48 | 792.19 | 886.46 |
| Clec3b | 5842.58 | -0.82 | 0.25 | 10.80 | 0.0010133 | 0.02814 | 6660.61 | 7998.59 | 7367.72 | 7628.46 | 4980.90 | 2080.18 | 4116.88 | 5907.30 |
| Syt7 | 128.04 | -0.80 | 0.24 | 10.80 | 0.0010132 | 0.02814 | 198.02 | 149.38 | 157.74 | 145.96 | 79.67 | 71.21 | 111.65 | 110.69 |
| D16Ertd472e | 262.24 | 0.64 | 0.19 | 10.80 | 0.0010177 | 0.0281635 | 233.41 | 221.20 | 172.30 | 213.76 | 377.10 | 323.27 | 232.16 | 324.70 |
| Atad2b | 193.21 | 0.71 | 0.21 | 10.80 | 0.0010167 | 0.0281635 | 147.13 | 188.64 | 107.99 | 156.32 | 268.75 | 256.74 | 214.44 | 205.70 |
| Tmem200c | 178.44 | -1.18 | 0.36 | 10.77 | 0.0010304 | 0.0284658 | 129.43 | 342.81 | 274.23 | 246.72 | 132.78 | 86.21 | 57.60 | 157.74 |
| Plekhg2 | 1921.40 | 0.59 | 0.18 | 10.74 | 0.0010467 | 0.0288651 | 1711.34 | 1421.04 | 1564.06 | 1528.33 | 1516.90 | 2774.51 | 2650.36 | 2204.63 |
| Rpl32 | 6501.97 | -0.26 | 0.08 | 10.74 | 0.0010498 | 0.0288653 | 7469.26 | 7249.77 | 6898.14 | 6753.65 | 5796.71 | 5959.43 | 6109.75 | 5779.08 |
| Ndufc1 | 908.95 | -0.43 | 0.13 | 10.73 | 0.0010521 | 0.0288653 | 1105.12 | 958.53 | 1202.47 | 808.89 | 910.35 | 654.04 | 867.50 | 764.70 |
| Kif26a | 691.65 | 0.68 | 0.21 | 10.73 | 0.0010515 | 0.0288653 | 564.18 | 335.15 | 695.27 | 519.80 | 708.52 | 858.31 | 884.34 | 967.64 |
| ATF7 | 2725.18 | 0.27 | 0.08 | 10.73 | 0.0010569 | 0.0289477 | 2492.34 | 2783.66 | 2233.86 | 2571.70 | 2771.42 | 3146.50 | 2614.92 | 3187.03 |
| 1700020I14Ri | 9254.27 | 0.39 | 0.12 | 10.72 | 0.0010613 | 0.0289669 | 8863.11 | 8484.08 | 6608.14 | 8969.39 | 10808.41 | 9670.02 | 8782.27 | 11848.73 |
| Smurf2 | 2976.84 | 0.31 | 0.09 | 10.72 | 0.0010598 | 0.0289669 | 2778.85 | 2656.30 | 2734.99 | 2523.67 | 3205.88 | 3191.48 | 3085.45 | 3638.10 |
| Ahnak2 | 3863.26 | 0.45 | 0.14 | 10.70 | 0.0010689 | 0.0290483 | 3377.32 | 3317.99 | 3226.41 | 3388.12 | 3817.73 | 3962.65 | 3911.30 | 5904.53 |
| Kcnk1 | 2844.72 | -0.35 | 0.11 | 10.70 | 0.0010689 | 0.0290483 | 2992.35 | 3026.88 | 3736.04 | 2653.62 | 3131.52 | 2184.19 | 2602.51 | 2430.63 |
| Acaca | 2027.79 | -0.69 | 0.21 | 10.70 | 0.0010697 | 0.0290483 | 1966.88 | 1636.49 | 4369.43 | 1422.86 | 1493.53 | 1032.59 | 3034.05 | 1266.51 |
| Abat | 1203.00 | 0.43 | 0.13 | 10.69 | 0.0010766 | 0.0291624 | 1071.94 | 1133.76 | 927.03 | 1065.97 | 1210.97 | 1230.30 | 1216.63 | 1767.39 |
| Slc10a5 | 132.56 | 1.03 | 0.30 | 10.69 | 0.0010776 | 0.0291624 | 103.99 | 96.71 | 75.23 | 54.62 | 234.76 | 182.72 | 209.12 | 103.31 |
| BC027231 | 527.23 | 0.37 | 0.11 | 10.68 | 0.0010801 | 0.0291805 | 477.89 | 445.27 | 458.66 | 452.00 | 641.60 | 580.95 | 560.02 | 601.43 |
| Crebbp | 1552.37 | 0.35 | 0.11 | 10.66 | 0.0010943 | 0.0294958 | 1383.89 | 1324.32 | 1246.15 | 1582.00 | 1602.94 | 1701.62 | 1757.16 | 1820.89 |
| Fam195a | 159.43 | -0.59 | 0.18 | 10.66 | 0.0010955 | 0.0294958 | 181.42 | 161.83 | 242.68 | 152.55 | 157.21 | 97.45 | 157.73 | 124.53 |
| Hibadh | 4080.36 | -0.34 | 0.11 | 10.64 | 0.0011057 | 0.0297209 | 4243.50 | 3997.86 | 5614.37 | 4012.45 | 4237.32 | 3204.60 | 3660.53 | 3672.23 |
| Nsd1 | 5313.69 | 0.29 | 0.09 | 10.61 | 0.0011252 | 0.0299309 | 4818.74 | 5085.66 | 4823.24 | 4509.65 | 5748.91 | 6315.50 | 5553.27 | 5654.55 |
| Fblim1 | 5217.77 | 0.40 | 0.12 | 10.62 | 0.0011196 | 0.0299309 | 4701.48 | 4530.27 | 4631.52 | 4347.68 | 4587.87 | 6065.31 | 6011.39 | 6866.64 |
| Itpripl2 | 3707.56 | 0.32 | 0.10 | 10.62 | 0.0011185 | 0.0299309 | 3471.35 | 3051.78 | 3291.94 | 3470.99 | 3787.99 | 4635.43 | 3970.67 | 3980.32 |
| Itga6 | 3660.28 | 0.39 | 0.12 | 10.60 | 0.0011296 | 0.0299309 | 3187.05 | 3171.48 | 3329.55 | 3036.88 | 3957.95 | 5057.08 | 3793.45 | 3748.79 |
| Nbea | 1097.88 | 0.42 | 0.13 | 10.60 | 0.0011304 | 0.0299309 | 1029.90 | 1147.17 | 803.27 | 910.59 | 968.77 | 1263.10 | 1171.44 | 1488.82 |
| Abcd2 | 664.40 | -0.82 | 0.25 | 10.63 | 0.0011155 | 0.0299309 | 528.78 | 660.72 | 1190.34 | 945.44 | 468.45 | 300.78 | 599.90 | 620.80 |
| Wisp1 | 366.46 | 1.83 | 0.56 | 10.61 | 0.0011273 | 0.0299309 | 223.46 | 98.63 | 189.29 | 140.31 | 231.57 | 1025.10 | 643.32 | 380.04 |
| Pikfyve | 350.57 | 0.44 | 0.13 | 10.61 | 0.0011238 | 0.0299309 | 351.78 | 296.85 | 275.44 | 294.74 | 372.85 | 431.03 | 345.58 | 436.31 |
| Inpp4b | 82.97 | 0.90 | 0.28 | 10.61 | 0.0011221 | 0.0299309 | 73.01 | 59.37 | 43.68 | 73.45 | 53.11 | 171.47 | 76.21 | 113.46 |
| Sod1 | 7616.39 | -0.35 | 0.11 | 10.58 | 0.0011402 | 0.0301399 | 8813.33 | 7820.49 | 9513.00 | 7264.97 | 7751.25 | 5650.22 | 7269.67 | 6848.19 |
| Gm15564 | 1549.97 | -1.32 | 0.41 | 10.58 | 0.0011454 | 0.0302277 | 2366.23 | 835.00 | 3226.41 | 2321.21 | 1026.14 | 1017.60 | 871.05 | 736.11 |
| Itpr1 | 9370.93 | 0.23 | 0.07 | 10.57 | 0.0011507 | 0.0302855 | 8992.54 | 8761.78 | 8374.84 | 8480.67 | 10193.37 | 10026.09 | 10004.22 | 10133.92 |
| Ptgfr | 186.80 | -0.79 | 0.25 | 10.57 | 0.0011514 | 0.0302855 | 255.54 | 275.78 | 246.32 | 178.92 | 135.97 | 144.30 | 84.18 | 173.42 |
| Grpel1 | 2081.86 | -0.32 | 0.10 | 10.56 | 0.0011567 | 0.0303753 | 2209.14 | 2221.57 | 2692.52 | 1950.20 | 2165.93 | 1702.56 | 1909.57 | 1803.37 |
| Gng12 | 6846.08 | 0.28 | 0.09 | 10.55 | 0.0011637 | 0.0305101 | 6225.86 | 6864.83 | 5847.34 | 6237.61 | 6466.99 | 8420.04 | 6942.70 | 7763.25 |
| 3010026O09R | 189.97 | -0.57 | 0.17 | 10.54 | 0.0011662 | 0.0305247 | 227.88 | 210.67 | 245.11 | 194.93 | 200.77 | 141.49 | 178.11 | 120.84 |
| Tet3 | 3107.23 | 0.30 | 0.09 | 10.52 | 0.001179 | 0.0307073 | 2976.86 | 2781.75 | 2469.26 | 3075.49 | 3435.32 | 3495.08 | 3239.63 | 3384.43 |
| S100pbp | 1379.92 | 0.31 | 0.10 | 10.53 | 0.0011771 | 0.0307073 | 1233.45 | 1405.71 | 1062.93 | 1336.23 | 1430.85 | 1578.87 | 1398.29 | 1593.05 |
| Npm3 | 372.38 | -0.40 | 0.12 | 10.53 | 0.0011757 | 0.0307073 | 422.58 | 407.93 | 459.88 | 393.62 | 315.49 | 308.28 | 344.70 | 326.54 |
| mt-Ti | 158.95 | -1.20 | 0.36 | 10.52 | 0.0011841 | 0.0307897 | 172.57 | 288.23 | 192.93 | 239.18 | 108.35 | 38.42 | 80.64 | 151.28 |
| Sptan1 | 11863.63 | 0.33 | 0.10 | 10.50 | 0.0011945 | 0.031009 | 11054.55 | 10623.30 | 10794.35 | 9737.79 | 11505.25 | 13617.68 | 14158.32 | 13417.80 |
| Ccnd3 | 2748.89 | -0.41 | 0.13 | 10.49 | 0.0012009 | 0.0310747 | 3282.18 | 2982.84 | 3552.82 | 2676.22 | 2059.71 | 2548.69 | 2594.54 | 2294.11 |
| 2410066E13R | 440.90 | 0.48 | 0.15 | 10.49 | 0.0012001 | 0.0310747 | 366.16 | 383.99 | 408.91 | 329.58 | 455.71 | 567.83 | 442.17 | 572.83 |
| Hist2h2be | 441.36 | -0.45 | 0.14 | 10.48 | 0.0012073 | 0.0311906 | 469.04 | 533.37 | 558.16 | 482.13 | 345.23 | 419.78 | 338.50 | 384.66 |
| Lamc1 | 12855.69 | 0.35 | 0.11 | 10.47 | 0.0012115 | 0.0312484 | 11339.96 | 12484.82 | 10882.92 | 11191.73 | 11492.50 | 14883.59 | 14819.36 | 15750.65 |
| Nedd9 | 1216.60 | 0.47 | 0.15 | 10.46 | 0.0012181 | 0.0313161 | 959.10 | 908.74 | 1115.11 | 1095.16 | 1276.83 | 1740.98 | 1403.60 | 1233.30 |
| Lrig2 | 1120.51 | 0.31 | 0.10 | 10.47 | 0.0012165 | 0.0313161 | 984.54 | 1050.46 | 969.50 | 1034.89 | 1154.67 | 1368.05 | 1194.48 | 1207.47 |
| Jak2 | 3481.87 | 0.31 | 0.10 | 10.45 | 0.0012297 | 0.0315318 | 3383.96 | 3348.63 | 2662.18 | 3338.21 | 3450.19 | 3654.37 | 3633.95 | 4383.43 |
| Smyd1 | 44.13 | -1.20 | 0.37 | 10.44 | 0.0012304 | 0.0315318 | 48.67 | 59.37 | 60.67 | 76.28 | 28.68 | 15.93 | 27.47 | 35.98 |
| Slain1 | 42.36 | -1.07 | 0.33 | 10.44 | 0.0012325 | 0.0315341 | 56.42 | 55.54 | 57.03 | 57.44 | 33.99 | 18.74 | 29.24 | 30.44 |
| Wdfy3 | 4068.13 | 0.28 | 0.09 | 10.42 | 0.0012493 | 0.0316654 | 3666.05 | 3859.01 | 3530.97 | 3632.96 | 4805.63 | 4516.42 | 4262.20 | 4271.82 |
| Mkl2 | 2559.92 | 0.33 | 0.10 | 10.42 | 0.0012487 | 0.0316654 | 2371.76 | 2492.56 | 2105.24 | 2243.05 | 2696.00 | 3228.03 | 2701.76 | 2640.94 |
| Rad54l2 | 1850.79 | 0.28 | 0.09 | 10.42 | 0.0012433 | 0.0316654 | 1648.28 | 1738.95 | 1565.28 | 1798.59 | 1981.10 | 1985.54 | 1927.30 | 2161.27 |
| Rassf8 | 1532.61 | 0.38 | 0.12 | 10.41 | 0.001251 | 0.0316654 | 1337.43 | 1518.71 | 1329.88 | 1214.75 | 1474.41 | 1975.23 | 1636.65 | 1773.85 |
| Mrps14 | 1199.92 | -0.36 | 0.11 | 10.42 | 0.0012499 | 0.0316654 | 1438.10 | 1247.72 | 1494.90 | 1182.74 | 1038.88 | 1011.04 | 1085.49 | 1100.47 |
| Rasa2 | 672.54 | 0.44 | 0.13 | 10.42 | 0.0012492 | 0.0316654 | 641.61 | 635.83 | 436.82 | 661.05 | 779.69 | 784.28 | 603.44 | 837.57 |
| Tmem120b | 432.76 | -0.68 | 0.22 | 10.41 | 0.0012515 | 0.0316654 | 388.29 | 360.05 | 982.85 | 292.86 | 340.98 | 262.36 | 587.49 | 247.21 |
| Alpl | 5354.39 | -0.47 | 0.15 | 10.37 | 0.0012807 | 0.0319095 | 5730.27 | 5154.61 | 7940.45 | 5214.96 | 4581.49 | 3314.23 | 6153.17 | 4745.95 |
| Itch | 3522.23 | 0.24 | 0.07 | 10.37 | 0.0012778 | 0.0319095 | 3236.83 | 3395.55 | 3089.30 | 3344.81 | 3592.54 | 3855.83 | 3682.69 | 3980.32 |
| Suco | 3379.76 | 0.36 | 0.11 | 10.38 | 0.001275 | 0.0319095 | 3390.59 | 2931.13 | 2669.47 | 2991.68 | 3662.64 | 3408.87 | 3803.20 | 4180.49 |
| Pros1 | 2525.95 | 0.46 | 0.14 | 10.38 | 0.001275 | 0.0319095 | 2085.24 | 2409.25 | 1858.92 | 2474.70 | 2131.94 | 3897.99 | 2602.51 | 2747.02 |
| Bach1 | 1117.54 | 0.32 | 0.10 | 10.37 | 0.0012792 | 0.0319095 | 952.46 | 999.70 | 984.06 | 1084.80 | 1161.04 | 1346.49 | 1112.96 | 1298.79 |
| Zscan29 | 1033.53 | 0.35 | 0.11 | 10.37 | 0.0012812 | 0.0319095 | 863.97 | 1005.45 | 809.33 | 984.04 | 1149.36 | 1153.47 | 1164.35 | 1138.29 |
| Zfp809 | 774.80 | 0.44 | 0.13 | 10.39 | 0.0012682 | 0.0319095 | 707.99 | 699.98 | 559.37 | 660.11 | 941.16 | 854.56 | 979.16 | 796.06 |
| Klf2 | 641.05 | -0.45 | 0.14 | 10.38 | 0.0012731 | 0.0319095 | 790.95 | 611.89 | 791.13 | 774.05 | 478.01 | 595.94 | 544.96 | 541.47 |
| Dio2 | 432.65 | 0.94 | 0.28 | 10.38 | 0.0012742 | 0.0319095 | 413.73 | 295.89 | 224.48 | 298.51 | 552.37 | 838.63 | 538.76 | 298.87 |
| Hs6st2 | 53.97 | 1.75 | 0.55 | 10.39 | 0.0012687 | 0.0319095 | 21.02 | 22.02 | 19.41 | 44.26 | 14.87 | 165.85 | 78.86 | 65.49 |
| Ndufs7 | 1706.84 | -0.34 | 0.11 | 10.36 | 0.0012881 | 0.0320105 | 1812.00 | 1936.21 | 2153.77 | 1597.07 | 1619.93 | 1333.38 | 1673.87 | 1528.48 |
| Orm1 | 156.16 | -1.06 | 0.33 | 10.36 | 0.0012893 | 0.0320105 | 172.57 | 163.74 | 317.91 | 155.38 | 123.22 | 36.54 | 147.09 | 132.83 |
| Wdr26 | 7334.21 | 0.31 | 0.10 | 10.34 | 0.0012993 | 0.0321607 | 7057.75 | 7105.18 | 5370.48 | 7461.78 | 7935.02 | 8227.95 | 6882.44 | 8633.11 |
| Gm4956 | 28.17 | -1.48 | 0.46 | 10.35 | 0.0012981 | 0.0321607 | 46.46 | 46.92 | 30.33 | 47.08 | 4.25 | 12.18 | 15.06 | 23.06 |
| Slc29a1 | 4260.33 | -0.60 | 0.19 | 10.34 | 0.0013053 | 0.0321937 | 4567.62 | 6094.94 | 5065.92 | 4854.30 | 3835.79 | 2445.62 | 2871.01 | 4347.46 |
| Osmr | 2999.52 | 0.28 | 0.09 | 10.33 | 0.0013059 | 0.0321937 | 2570.88 | 2735.78 | 2659.76 | 2904.11 | 3204.81 | 3528.81 | 3229.00 | 3163.04 |
| Chd9 | 2026.98 | 0.31 | 0.10 | 10.33 | 0.0013088 | 0.0321937 | 1967.98 | 2003.24 | 1657.50 | 1743.97 | 2197.80 | 2342.54 | 1944.13 | 2358.68 |
| Pkp1 | 834.41 | 0.77 | 0.24 | 10.33 | 0.0013084 | 0.0321937 | 783.21 | 608.06 | 581.22 | 542.40 | 1123.86 | 1525.46 | 830.29 | 680.76 |

| | | | | | | | | | | | | | | |
|---|---|---|---|---|---|---|---|---|---|---|---|---|---|---|
| Amy1 | 351.02 | -0.55 | 0.17 | 10.31 | 0.0013242 | 0.0325243 | 383.86 | 411.76 | 395.57 | 466.13 | 316.55 | 219.26 | 298.62 | 316.40 |
| Hsp90ab1 | 55282.28 | -0.31 | 0.10 | 10.29 | 0.0013375 | 0.0327028 | 58321.54 | 54624.66 | 71143.68 | 58035.97 | 49788.81 | 45434.10 | 52079.53 | 52829.93 |
| Esyt2 | 5010.04 | 0.43 | 0.14 | 10.29 | 0.0013377 | 0.0327028 | 4328.68 | 4690.18 | 4184.99 | 4224.33 | 4766.32 | 7625.45 | 4903.75 | 5356.60 |
| Nudc | 1353.44 | -0.29 | 0.09 | 10.29 | 0.0013344 | 0.0327028 | 1383.89 | 1396.14 | 1732.73 | 1290.09 | 1414.92 | 1056.96 | 1384.11 | 1168.73 |
| Ccdc124 | 1203.00 | -0.36 | 0.11 | 10.28 | 0.0013461 | 0.0328595 | 1306.46 | 1155.79 | 1644.15 | 1136.59 | 1157.85 | 877.05 | 1318.54 | 1027.60 |
| 0610010F05Rik | 671.78 | 0.37 | 0.11 | 10.27 | 0.001351 | 0.0329292 | 692.50 | 640.62 | 474.44 | 618.68 | 645.85 | 767.42 | 701.80 | 832.96 |
| Btbd3 | 2188.40 | 0.42 | 0.13 | 10.26 | 0.0013577 | 0.03304 | 2004.49 | 2001.32 | 1809.17 | 1743.97 | 2341.20 | 3034.06 | 2442.13 | 2130.83 |
| Lrrc14b | 95.76 | -1.64 | 0.51 | 10.26 | 0.0013626 | 0.033109 | 89.60 | 147.47 | 197.78 | 140.31 | 70.11 | 10.31 | 26.58 | 83.94 |
| Mgst3 | 4369.73 | -0.35 | 0.11 | 10.24 | 0.001376 | 0.0333839 | 4953.70 | 4417.28 | 5741.78 | 3938.06 | 4783.32 | 3686.23 | 4143.47 | 3294.03 |
| Trem2 | 195.74 | 0.87 | 0.27 | 10.23 | 0.0013802 | 0.0334342 | 203.55 | 135.02 | 106.78 | 138.43 | 171.02 | 357.00 | 263.18 | 190.94 |
| Ston2 | 755.12 | 0.50 | 0.16 | 10.22 | 0.0013858 | 0.0335193 | 768.83 | 635.83 | 558.16 | 565.94 | 811.56 | 995.11 | 946.37 | 759.17 |
| Tnks | 778.40 | 0.48 | 0.15 | 10.21 | 0.0013976 | 0.0337537 | 707.99 | 728.71 | 566.65 | 613.03 | 1059.07 | 925.77 | 816.11 | 809.90 |
| Exoc4 | 3476.77 | 0.26 | 0.08 | 10.19 | 0.0014095 | 0.0339727 | 3225.77 | 3482.69 | 2965.53 | 3167.77 | 3563.85 | 4138.81 | 3420.40 | 3849.34 |
| Dennd4a | 950.73 | 0.43 | 0.14 | 10.19 | 0.0014109 | 0.0339727 | 839.63 | 767.01 | 911.26 | 737.33 | 956.03 | 1279.97 | 1002.19 | 1112.46 |
| Kif5b | 6022.83 | 0.40 | 0.12 | 10.18 | 0.0014159 | 0.0339908 | 6044.44 | 5534.76 | 4642.44 | 5015.33 | 6707.06 | 6133.72 | 5876.70 | 8228.16 |
| Klrb1c | 32.87 | -1.54 | 0.48 | 10.19 | 0.0014156 | 0.0339908 | 34.29 | 33.52 | 50.96 | 75.33 | 26.56 | 15.93 | 9.75 | 16.60 |
| Egfr | 1827.58 | 0.41 | 0.13 | 10.17 | 0.0014241 | 0.0340849 | 1639.43 | 1617.34 | 1527.66 | 1594.24 | 1837.70 | 2603.03 | 1961.85 | 1839.34 |
| Htr2a | 37.82 | 1.39 | 0.43 | 10.18 | 0.0014235 | 0.0340849 | 18.81 | 28.73 | 19.41 | 16.95 | 36.12 | 71.21 | 74.43 | 36.90 |
| Uggt2 | 453.47 | 0.42 | 0.13 | 10.17 | 0.0014273 | 0.0341107 | 465.72 | 433.78 | 311.84 | 394.56 | 495.01 | 554.71 | 430.65 | 541.47 |
| Cfd | 14779.62 | -1.43 | 0.45 | 10.13 | 0.0014576 | 0.0344961 | 13426.31 | 9876.39 | 37243.89 | 20828.77 | 8768.89 | 3166.18 | 16710.32 | 8216.16 |
| Atp5e | 4000.53 | -0.37 | 0.12 | 10.15 | 0.0014458 | 0.0344961 | 4735.77 | 4685.40 | 4695.83 | 3654.62 | 3679.64 | 3168.06 | 4008.78 | 3376.13 |
| Cox6a1 | 3611.88 | -0.44 | 0.14 | 10.14 | 0.0014492 | 0.0344961 | 4164.96 | 4101.28 | 4848.72 | 3217.68 | 2958.37 | 2618.03 | 3726.99 | 3258.98 |
| Tnrc6b | 2545.42 | 0.28 | 0.09 | 10.14 | 0.0014522 | 0.0344961 | 2273.30 | 2408.29 | 2117.37 | 2508.60 | 2814.97 | 2855.09 | 2507.70 | 2878.01 |
| Adamdec1 | 1113.10 | 0.60 | 0.19 | 10.13 | 0.0014586 | 0.0344961 | 898.26 | 936.50 | 709.84 | 1136.59 | 1260.89 | 1838.43 | 1104.10 | 1020.22 |
| Sema6d | 839.85 | 0.37 | 0.12 | 10.13 | 0.001458 | 0.0344961 | 705.77 | 802.44 | 724.40 | 759.93 | 785.00 | 1081.32 | 847.12 | 1012.84 |
| Cck | 85.52 | -1.29 | 0.39 | 10.14 | 0.0014548 | 0.0344961 | 126.11 | 130.23 | 120.13 | 100.76 | 65.86 | 15.93 | 53.17 | 71.95 |
| Zfp397 | 1930.78 | 0.37 | 0.11 | 10.12 | 0.0014658 | 0.0345636 | 1845.19 | 1923.76 | 1443.94 | 1697.83 | 2258.35 | 2241.35 | 1773.11 | 2262.74 |
| Bcas1 | 459.55 | -0.62 | 0.19 | 10.12 | 0.001464 | 0.0345636 | 690.29 | 376.33 | 563.01 | 585.72 | 401.53 | 332.64 | 359.76 | 367.13 |
| Bace1 | 2972.63 | 0.37 | 0.12 | 10.11 | 0.0014744 | 0.0347151 | 2632.83 | 3057.53 | 2453.48 | 2418.20 | 2729.99 | 3504.45 | 3333.56 | 3651.01 |
| Ncor1 | 6113.88 | 0.24 | 0.08 | 10.10 | 0.0014832 | 0.0348699 | 5887.35 | 5950.35 | 5262.49 | 5529.48 | 6459.55 | 6546.00 | 6321.53 | 6954.27 |
| Epha3 | 732.84 | 1.32 | 0.42 | 10.09 | 0.0014941 | 0.0350751 | 459.09 | 547.73 | 339.75 | 435.05 | 622.48 | 2126.09 | 706.23 | 626.34 |
| Pcf11 | 2010.60 | 0.31 | 0.10 | 10.05 | 0.0015205 | 0.0356434 | 1918.20 | 1947.70 | 1509.46 | 1968.09 | 2229.67 | 2317.24 | 1932.61 | 2261.82 |
| Ldhb | 8276.85 | -0.33 | 0.10 | 10.04 | 0.0015358 | 0.0359089 | 9176.18 | 8779.02 | 9656.18 | 8204.76 | 9326.57 | 6308.00 | 8090.21 | 6673.85 |
| Wdr18 | 1115.06 | -0.35 | 0.11 | 10.03 | 0.0015364 | 0.0359089 | 1235.66 | 1050.46 | 1511.89 | 1117.76 | 1034.63 | 941.70 | 1059.79 | 968.56 |
| Mrps12 | 604.23 | -0.35 | 0.11 | 10.03 | 0.0015408 | 0.0359592 | 651.57 | 630.08 | 780.21 | 577.24 | 593.80 | 476.00 | 618.51 | 506.42 |
| Nes | 1049.22 | 0.51 | 0.16 | 10.02 | 0.0015468 | 0.0360464 | 816.40 | 972.89 | 872.43 | 796.65 | 961.34 | 1143.16 | 1509.05 | 1321.86 |
| Tet2 | 1871.22 | 0.39 | 0.12 | 10.02 | 0.0015512 | 0.0360668 | 1844.09 | 1727.46 | 1383.27 | 1652.63 | 2267.91 | 2267.58 | 1853.75 | 1973.10 |
| Rpl7a | 1437.94 | -0.30 | 0.09 | 10.02 | 0.0015522 | 0.0360668 | 1419.29 | 1406.67 | 1860.13 | 1386.14 | 1541.33 | 1057.89 | 1704.88 | 1127.22 |
| Rnf167 | 2863.32 | -0.35 | 0.11 | 10.01 | 0.0015604 | 0.0361382 | 2857.39 | 3357.25 | 3372.02 | 3129.16 | 2916.94 | 2312.56 | 2375.67 | 2585.60 |
| Mmp17 | 1777.38 | 0.75 | 0.24 | 10.00 | 0.0015621 | 0.0361382 | 1358.45 | 879.05 | 1801.89 | 1129.06 | 1469.09 | 2507.46 | 3180.26 | 1893.77 |
| Atp5k | 1774.80 | -0.45 | 0.14 | 10.01 | 0.0015603 | 0.0361382 | 2081.92 | 2250.29 | 2215.66 | 1634.74 | 1429.79 | 1368.98 | 1472.72 | 1744.33 |
| Mapk8ip2 | 30.88 | -1.57 | 0.49 | 10.00 | 0.0015664 | 0.0361856 | 37.61 | 33.52 | 66.74 | 37.67 | 13.81 | 4.69 | 34.56 | 18.45 |
| Ear2 | 69.12 | -0.90 | 0.28 | 9.97 | 0.0015897 | 0.0366703 | 68.59 | 97.67 | 78.87 | 118.65 | 44.61 | 51.54 | 47.85 | 45.20 |
| Pcp4 | 1298.12 | -1.92 | 0.59 | 9.97 | 0.001593 | 0.0366942 | 1684.79 | 2912.93 | 1760.63 | 1945.49 | 611.86 | 182.72 | 298.62 | 987.93 |
| Fbln1 | 32331.40 | -0.37 | 0.12 | 9.95 | 0.0016093 | 0.0370163 | 32754.36 | 35214.69 | 38502.18 | 39043.46 | 27227.64 | 24752.25 | 28490.31 | 32666.33 |
| 3425401B19Rik | 46.46 | -1.52 | 0.48 | 9.95 | 0.0016126 | 0.0370394 | 51.99 | 77.56 | 74.02 | 73.45 | 18.06 | 7.50 | 23.93 | 45.20 |
| Gm340 | 94.70 | 0.70 | 0.22 | 9.94 | 0.001619 | 0.0371314 | 79.65 | 67.99 | 74.02 | 64.03 | 120.03 | 122.75 | 126.71 | 102.39 |
| Gpr123 | 400.36 | 0.63 | 0.20 | 9.93 | 0.0016282 | 0.0372371 | 294.26 | 372.50 | 208.70 | 472.72 | 371.79 | 584.70 | 432.42 | 465.83 |
| Slc41a2 | 391.80 | 0.59 | 0.19 | 9.93 | 0.0016278 | 0.0372371 | 370.59 | 292.06 | 321.55 | 298.51 | 301.68 | 560.34 | 429.76 | 559.92 |
| D10Jhu81e | 1960.52 | -0.26 | 0.08 | 9.91 | 0.0016404 | 0.0373763 | 1981.26 | 2065.48 | 2436.49 | 1898.40 | 1968.35 | 1713.81 | 1962.74 | 1657.62 |
| Cystm1 | 1412.52 | -0.42 | 0.13 | 9.91 | 0.0016437 | 0.0373763 | 1589.65 | 1495.73 | 1886.83 | 1296.68 | 1663.49 | 1116.92 | 1212.20 | 1038.67 |
| Rapgef3 | 575.80 | 0.48 | 0.15 | 9.91 | 0.0016436 | 0.0373763 | 564.18 | 411.76 | 526.61 | 409.63 | 605.48 | 696.20 | 727.50 | 665.08 |
| Tmem243 | 354.13 | -0.50 | 0.16 | 9.91 | 0.0016425 | 0.0373763 | 420.37 | 404.10 | 359.16 | 500.03 | 277.25 | 260.49 | 267.61 | 344.07 |
| Thbs1 | 5944.84 | 1.16 | 0.37 | 9.91 | 0.0016471 | 0.0373966 | 3931.54 | 2596.93 | 3271.31 | 5617.99 | 4619.73 | 12834.33 | 7656.02 | 7030.83 |
| Ndufa8 | 1666.99 | -0.30 | 0.09 | 9.90 | 0.0016497 | 0.0373966 | 1782.14 | 1640.32 | 2157.41 | 1630.03 | 1651.80 | 1369.92 | 1612.73 | 1491.58 |
| Dnm3os | 581.18 | 0.83 | 0.26 | 9.90 | 0.0016516 | 0.0373966 | 419.26 | 508.47 | 321.55 | 530.16 | 521.57 | 1253.73 | 635.34 | 459.37 |
| Gsn | 99007.69 | -0.61 | 0.19 | 9.89 | 0.0016581 | 0.0374885 | 103184.69 | 103875.44 | 146410.46 | 117721.75 | 93610.94 | 51598.74 | 73008.63 | 102650.86 |
| Pfdn1 | 1346.12 | -0.30 | 0.10 | 9.87 | 0.0016814 | 0.0379621 | 1538.77 | 1431.57 | 1601.68 | 1322.10 | 1286.39 | 1163.78 | 1225.49 | 1199.17 |
| Gnas | 25608.84 | -0.30 | 0.10 | 9.85 | 0.0016961 | 0.0382398 | 26057.24 | 27079.16 | 32870.82 | 25901.53 | 22207.44 | 23565.05 | 24294.56 | 22894.94 |
| Slc16a10 | 969.91 | -0.78 | 0.25 | 9.85 | 0.0016991 | 0.0382541 | 1039.86 | 1184.52 | 1166.07 | 1589.54 | 678.78 | 466.63 | 512.17 | 1121.69 |
| Ep300 | 3157.55 | 0.36 | 0.11 | 9.83 | 0.0017139 | 0.0385322 | 2812.04 | 3401.29 | 2307.87 | 2796.76 | 3340.78 | 3441.67 | 3420.40 | 3739.57 |
| Gng8 | 173.38 | 0.62 | 0.20 | 9.83 | 0.00172 | 0.0386145 | 140.49 | 165.66 | 111.63 | 144.08 | 163.59 | 198.65 | 227.73 | 235.22 |
| Il13ra1 | 1175.15 | 0.44 | 0.14 | 9.82 | 0.0017301 | 0.0387878 | 946.93 | 1017.90 | 1005.90 | 1061.26 | 1257.71 | 1688.51 | 1265.37 | 1157.66 |
| Ndufb7 | 1499.57 | -0.39 | 0.13 | 9.81 | 0.0017328 | 0.0387944 | 1730.14 | 1679.58 | 1941.43 | 1381.43 | 1281.08 | 1168.46 | 1415.12 | 1399.34 |
| Ankhd1 | 3113.89 | 0.26 | 0.08 | 9.79 | 0.0017528 | 0.039131 | 2719.11 | 2791.32 | 2932.77 | 2779.81 | 3782.68 | 3293.62 | 3336.22 | 3275.58 |
| Reg3g | 96.97 | 1.51 | 0.48 | 9.79 | 0.0017523 | 0.039131 | 34.29 | 31.60 | 55.82 | 75.33 | 128.53 | 230.51 | 177.22 | 42.43 |
| Lmcd1 | 4151.26 | -1.14 | 0.36 | 9.78 | 0.0017638 | 0.0392866 | 4235.75 | 7367.55 | 5267.34 | 6287.52 | 2785.22 | 1920.89 | 1411.58 | 3934.20 |
| Ptx3 | 32.60 | 1.76 | 0.57 | 9.78 | 0.0017647 | 0.0392866 | 14.38 | 13.41 | 10.92 | 26.37 | 19.12 | 117.13 | 33.67 | 25.83 |
| Dusp11 | 5834.93 | 0.27 | 0.09 | 9.77 | 0.0017728 | 0.0394125 | 5449.29 | 5434.22 | 5125.37 | 5305.36 | 6431.93 | 7007.95 | 5827.08 | 6098.24 |
| Ppil1 | 674.71 | -0.40 | 0.13 | 9.75 | 0.0017902 | 0.0397152 | 757.77 | 798.61 | 842.09 | 645.04 | 571.49 | 526.60 | 638.00 | 618.03 |
| Epyc | 69.94 | 1.65 | 0.53 | 9.75 | 0.0017914 | 0.0397152 | 21.02 | 51.71 | 37.62 | 27.31 | 64.80 | 219.26 | 103.68 | 34.13 |
| Dock9 | 4279.53 | 0.32 | 0.10 | 9.74 | 0.0018007 | 0.0397853 | 3943.71 | 4072.55 | 3449.68 | 3696.99 | 5192.29 | 4522.05 | 4907.29 | 4451.69 |
| Rarres2 | 2948.45 | -0.43 | 0.14 | 9.74 | 0.0018016 | 0.0397853 | 3837.51 | 3041.25 | 3640.18 | 2906.93 | 2345.45 | 2172.94 | 2906.45 | 2736.88 |
| Eno3 | 1808.03 | -1.18 | 0.38 | 9.74 | 0.0018046 | 0.0397853 | 2128.39 | 2847.82 | 3065.03 | 1922.89 | 1419.17 | 629.68 | 846.24 | 1605.04 |
| Tuba1b | 865.02 | -0.41 | 0.13 | 9.74 | 0.0018029 | 0.0397853 | 934.76 | 885.75 | 1190.34 | 814.54 | 753.14 | 625.93 | 1012.83 | 702.90 |
| Cox7b | 2579.06 | -0.32 | 0.10 | 9.74 | 0.0018078 | 0.0398008 | 2850.75 | 2974.22 | 3126.91 | 2447.40 | 2444.24 | 2251.65 | 2209.08 | 2328.24 |
| Glis3 | 440.89 | 0.59 | 0.19 | 9.73 | 0.0018104 | 0.0398045 | 310.85 | 391.65 | 294.85 | 461.42 | 372.85 | 654.98 | 539.64 | 500.88 |
| Zfyve16 | 958.59 | 0.36 | 0.12 | 9.73 | 0.0018147 | 0.0398424 | 858.43 | 930.76 | 765.65 | 881.40 | 1061.19 | 1212.50 | 844.47 | 1114.31 |
| Dync1h1 | 14328.89 | 0.24 | 0.08 | 9.72 | 0.0018235 | 0.0399187 | 13243.78 | 13255.66 | 13332.77 | 12591.05 | 14917.20 | 15125.34 | 16563.23 | 15602.13 |
| Plin2 | 5105.97 | -0.35 | 0.11 | 9.72 | 0.001824 | 0.0399187 | 4877.37 | 4851.06 | 7837.31 | 4652.78 | 4684.53 | 4100.39 | 5668.47 | 4175.88 |
| Rplp2 | 4611.88 | -0.32 | 0.10 | 9.72 | 0.0018256 | 0.0399187 | 4869.62 | 4352.16 | 6269.60 | 4276.12 | 4405.16 | 3669.36 | 5439.85 | 3613.19 |
| Lancl3 | 251.24 | 0.57 | 0.18 | 9.71 | 0.0018366 | 0.0401032 | 245.58 | 245.14 | 154.10 | 190.22 | 309.12 | 312.96 | 263.18 | 289.65 |
| Klhl30 | 193.63 | -1.14 | 0.36 | 9.70 | 0.0018443 | 0.040216 | 153.77 | 341.85 | 237.83 | 348.42 | 138.09 | 80.58 | 59.37 | 189.10 |
| Gzma | 80.16 | -1.73 | 0.52 | 9.68 | 0.0018596 | 0.040494 | 57.52 | 69.90 | 66.74 | 314.52 | 36.12 | 41.23 | 23.93 | 31.36 |
| Wwc2 | 1552.02 | 0.38 | 0.12 | 9.68 | 0.0018633 | 0.0405206 | 1623.95 | 1295.59 | 1161.22 | 1410.62 | 1530.70 | 1834.68 | 1829.82 | 1729.57 |
| Sqrdl | 3402.57 | -0.33 | 0.11 | 9.67 | 0.001871 | 0.0405346 | 4027.78 | 3556.42 | 4018.76 | 3512.42 | 2884.01 | 2775.45 | 3112.03 | 3333.69 |
| Gm20721 | 455.66 | -0.89 | 0.28 | 9.67 | 0.0018712 | 0.0405346 | 350.67 | 417.50 | 959.79 | 389.85 | 852.99 | 208.02 | 277.35 | 189.10 |
| Adamts12 | 210.46 | 1.10 | 0.35 | 9.67 | 0.0018716 | 0.0405346 | 134.96 | 192.47 | 100.71 | 145.96 | 132.78 | 520.04 | 289.76 | 166.96 |
| Flnb | 13243.78 | 0.29 | 0.09 | 9.66 | 0.0018831 | 0.0406004 | 11945.07 | 11869.10 | 12592.60 | 10927.12 | 13605.32 | 15321.17 | 15925.23 | 13764.64 |
| Lama2 | 2919.88 | 0.57 | 0.18 | 9.66 | 0.0018815 | 0.0406004 | 2282.15 | 2675.45 | 2053.06 | 2712.95 | 2416.62 | 4958.70 | 3431.92 | 2828.20 |
| Aldh1l1 | 817.75 | -0.45 | 0.15 | 9.66 | 0.0018828 | 0.0406004 | 859.54 | 732.54 | 1261.93 | 815.48 | 681.97 | 596.88 | 907.38 | 686.29 |
| Lnx1 | 409.92 | -0.46 | 0.15 | 9.66 | 0.0018848 | 0.0406004 | 444.70 | 398.35 | 521.76 | 446.35 | 471.64 | 260.49 | 428.88 | 307.17 |
| Gm13476 | 130.83 | -0.74 | 0.24 | 9.64 | 0.0019013 | 0.0409002 | 122.79 | 165.66 | 152.89 | 221.29 | 86.04 | 111.51 | 87.73 | 98.70 |

| | | | | | | | | | | | | | | |
|---|---|---|---|---|---|---|---|---|---|---|---|---|---|---|
| Mtmr11 | 1193.51 | 0.40 | 0.13 | 9.63 | 0.0019167 | 0.0411748 | 1175.92 | 1100.25 | 936.74 | 973.69 | 1070.75 | 1416.77 | 1449.68 | 1424.25 |
| Opn1sw | 105.60 | 0.78 | 0.25 | 9.62 | 0.001921 | 0.0412114 | 103.99 | 89.05 | 64.31 | 69.68 | 81.79 | 149.92 | 132.03 | 154.05 |
| Pappa2 | 239.76 | -1.11 | 0.35 | 9.62 | 0.0019244 | 0.0412295 | 597.36 | 192.47 | 296.07 | 230.71 | 175.27 | 105.88 | 129.37 | 190.94 |
| Synj2 | 2937.46 | 0.27 | 0.09 | 9.61 | 0.001938 | 0.0414089 | 2632.83 | 2632.36 | 2773.82 | 2570.76 | 3372.65 | 2975.03 | 3165.20 | 3377.05 |
| Cdk12 | 1406.73 | 0.31 | 0.10 | 9.61 | 0.0019372 | 0.0414089 | 1295.40 | 1304.21 | 1113.90 | 1406.85 | 1547.70 | 1531.09 | 1406.26 | 1648.40 |
| Bckdha | 2488.33 | -0.26 | 0.08 | 9.60 | 0.0019428 | 0.0414574 | 2570.88 | 2342.22 | 3159.68 | 2496.36 | 2696.00 | 2075.49 | 2476.69 | 2089.32 |
| Faah | 1076.28 | -0.41 | 0.13 | 9.60 | 0.0019455 | 0.0414597 | 1247.83 | 1126.10 | 1335.95 | 1096.10 | 1096.24 | 709.32 | 1018.14 | 980.55 |
| Taf1 | 2241.60 | 0.35 | 0.11 | 9.59 | 0.0019595 | 0.0417014 | 1953.60 | 2259.87 | 1828.58 | 1943.60 | 2658.82 | 2648.95 | 2105.40 | 2533.94 |
| Zdhhc20 | 3599.52 | 0.40 | 0.13 | 9.58 | 0.001966 | 0.041784 | 3529.98 | 3184.88 | 2453.48 | 3535.02 | 4431.71 | 4051.66 | 3595.85 | 4013.53 |
| Gm20412 | 52.43 | -1.16 | 0.38 | 9.57 | 0.0019743 | 0.0419047 | 32.08 | 54.58 | 135.90 | 34.84 | 84.98 | 20.61 | 41.65 | 14.76 |
| Mcts2 | 197.68 | -0.53 | 0.17 | 9.57 | 0.0019817 | 0.0420056 | 213.50 | 223.11 | 274.23 | 207.17 | 150.84 | 155.54 | 201.15 | 155.89 |
| Arhgap21 | 2155.87 | 0.32 | 0.10 | 9.56 | 0.0019902 | 0.0420747 | 2101.84 | 2074.10 | 1832.22 | 1740.20 | 2160.62 | 2471.85 | 2423.52 | 2442.62 |
| Mrpl30 | 1322.80 | -0.33 | 0.11 | 9.56 | 0.0019888 | 0.0420747 | 1479.03 | 1543.61 | 1503.39 | 1324.93 | 1285.32 | 1063.51 | 1170.56 | 1212.08 |
| Sept6 | 1021.49 | 0.41 | 0.13 | 9.55 | 0.0020049 | 0.0423288 | 852.90 | 874.26 | 889.42 | 949.20 | 1039.94 | 1093.50 | 1021.69 | 1451.00 |
| Epg5 | 1264.04 | 0.36 | 0.12 | 9.53 | 0.0020184 | 0.0425583 | 1232.34 | 1146.21 | 1008.33 | 1056.55 | 1403.23 | 1510.47 | 1487.78 | 1267.43 |
| Agps | 1565.28 | 0.33 | 0.11 | 9.53 | 0.0020217 | 0.0425709 | 1430.36 | 1527.33 | 1298.33 | 1413.44 | 1662.42 | 2018.34 | 1447.02 | 1724.96 |
| Cep350 | 1896.52 | 0.31 | 0.10 | 9.52 | 0.0020338 | 0.0427678 | 1682.58 | 1918.02 | 1612.60 | 1632.85 | 2128.75 | 2190.75 | 1901.60 | 2105.01 |
| Fosl2 | 9831.93 | 0.33 | 0.11 | 9.51 | 0.0020399 | 0.0428416 | 8635.23 | 8554.94 | 8271.70 | 9291.44 | 10866.84 | 11715.53 | 11894.30 | 9425.48 |
| Gnao1 | 3341.98 | -0.70 | 0.23 | 9.51 | 0.0020436 | 0.0428618 | 3458.07 | 4948.73 | 4047.88 | 4280.83 | 2497.35 | 1674.45 | 1966.29 | 3862.25 |
| Zfp318 | 1170.51 | 0.31 | 0.10 | 9.51 | 0.0020479 | 0.0428963 | 1104.02 | 1124.19 | 959.79 | 1023.59 | 1349.06 | 1339.94 | 1201.57 | 1261.90 |
| Clec10a | 765.90 | -0.48 | 0.16 | 9.50 | 0.0020547 | 0.0429723 | 818.61 | 907.78 | 748.66 | 1176.14 | 491.82 | 681.21 | 639.77 | 663.23 |
| A4gnt | 108.76 | -0.68 | 0.22 | 9.50 | 0.002057 | 0.0429723 | 136.07 | 113.95 | 173.52 | 105.47 | 91.35 | 82.46 | 81.52 | 85.79 |
| Dok2 | 229.11 | -0.84 | 0.27 | 9.48 | 0.0020752 | 0.0432963 | 261.07 | 303.55 | 254.81 | 383.26 | 117.91 | 106.82 | 144.44 | 261.05 |
| Skp1a | 7401.63 | -0.29 | 0.10 | 9.47 | 0.0020849 | 0.0434115 | 8232.56 | 8405.56 | 8294.76 | 7697.20 | 7138.33 | 6641.58 | 5905.94 | 6897.08 |
| Slit2 | 808.48 | 0.70 | 0.23 | 9.47 | 0.0020861 | 0.0434115 | 608.43 | 789.04 | 490.21 | 706.25 | 848.74 | 1559.20 | 781.55 | 684.45 |
| Las1l | 2928.45 | 0.32 | 0.10 | 9.46 | 0.0020988 | 0.0435034 | 2694.78 | 2705.14 | 2293.31 | 2779.81 | 3440.63 | 3247.70 | 3260.01 | 3006.23 |
| Manf | 1655.85 | -0.50 | 0.17 | 9.46 | 0.0020973 | 0.0435034 | 1839.66 | 1461.25 | 2817.50 | 1508.55 | 1326.75 | 1459.87 | 1587.91 | 1245.29 |
| Figf | 595.32 | 0.76 | 0.25 | 9.46 | 0.0020962 | 0.0435034 | 425.90 | 469.21 | 451.38 | 465.18 | 434.46 | 1220.93 | 711.55 | 583.90 |
| Usp9x | 2986.20 | 0.31 | 0.10 | 9.45 | 0.0021143 | 0.0437185 | 2792.12 | 2757.81 | 2710.72 | 2438.92 | 3496.93 | 3423.86 | 2961.39 | 3307.87 |
| Pf4 | 591.91 | -0.75 | 0.25 | 9.45 | 0.0021147 | 0.0437185 | 840.73 | 841.71 | 748.66 | 565.94 | 280.43 | 338.26 | 503.31 | 616.19 |
| Pln | 292.64 | -0.53 | 0.17 | 9.44 | 0.0021174 | 0.0437185 | 304.21 | 403.14 | 303.35 | 403.98 | 206.08 | 300.78 | 182.54 | 237.07 |
| Igfbp3 | 2582.85 | -0.60 | 0.20 | 9.43 | 0.0021291 | 0.0439031 | 1956.92 | 2990.50 | 3648.67 | 3732.77 | 1978.98 | 2394.08 | 2207.31 | 1753.56 |
| Ncald | 3298.76 | -0.62 | 0.21 | 9.43 | 0.0021391 | 0.0440028 | 3773.35 | 3879.12 | 4390.06 | 4010.57 | 2617.39 | 1797.20 | 2129.33 | 3793.07 |
| Unc13d | 459.42 | -0.38 | 0.12 | 9.43 | 0.0021394 | 0.0440028 | 506.65 | 479.74 | 586.07 | 468.01 | 408.97 | 401.98 | 467.87 | 356.06 |
| Rps9 | 10975.71 | -0.24 | 0.08 | 9.42 | 0.0021489 | 0.044028 | 12021.40 | 12612.17 | 11744.43 | 11323.56 | 9570.89 | 10293.14 | 10353.35 | 9886.70 |
| Ndn | 1092.70 | -0.53 | 0.18 | 9.42 | 0.0021442 | 0.044028 | 1066.41 | 1480.41 | 1350.51 | 1316.45 | 688.34 | 797.40 | 882.57 | 1159.51 |
| Clca2 | 892.86 | 0.86 | 0.28 | 9.42 | 0.0021468 | 0.044028 | 806.44 | 335.15 | 782.64 | 551.82 | 1389.43 | 1531.09 | 1031.44 | 714.89 |
| Tmem109 | 4715.21 | -0.26 | 0.08 | 9.41 | 0.0021522 | 0.0440391 | 5020.07 | 4940.11 | 5647.13 | 4426.78 | 4973.46 | 3894.24 | 4894.89 | 3924.98 |
| Sbf2 | 2903.40 | 0.28 | 0.09 | 9.40 | 0.0021664 | 0.0442715 | 2840.80 | 2854.52 | 2381.89 | 2529.32 | 3066.72 | 3086.54 | 3099.62 | 3367.82 |
| Ndufa5 | 1345.49 | -0.37 | 0.12 | 9.38 | 0.0021925 | 0.0446127 | 1581.91 | 1602.97 | 1561.64 | 1338.11 | 1166.35 | 1057.89 | 1113.84 | 1341.23 |
| Uhrf1bp1 | 994.70 | 0.30 | 0.10 | 9.38 | 0.0021943 | 0.0446127 | 855.12 | 861.81 | 959.79 | 847.50 | 1078.19 | 1035.40 | 1219.29 | 1100.47 |
| Cbl | 178.82 | 0.86 | 0.28 | 9.38 | 0.0021901 | 0.0446127 | 136.07 | 130.23 | 123.77 | 96.05 | 244.32 | 271.74 | 304.82 | 123.61 |
| B3galt5 | 54.95 | 1.65 | 0.54 | 9.38 | 0.0021934 | 0.0446127 | 36.51 | 18.19 | 33.98 | 18.83 | 38.24 | 164.92 | 94.81 | 34.13 |
| Asna1 | 1606.99 | -0.33 | 0.11 | 9.37 | 0.0022057 | 0.0446178 | 1761.12 | 1740.86 | 2073.69 | 1505.73 | 1444.66 | 1375.54 | 1490.44 | 1463.91 |
| Mysm1 | 1337.42 | 0.37 | 0.12 | 9.37 | 0.0022009 | 0.0446178 | 1229.02 | 1324.32 | 935.53 | 1323.99 | 1543.45 | 1553.58 | 1237.90 | 1551.54 |
| 9430008C03R | 1282.10 | 0.34 | 0.11 | 9.37 | 0.0022037 | 0.0446178 | 1137.20 | 1267.82 | 1005.90 | 1146.01 | 1513.71 | 1407.40 | 1408.92 | 1369.82 |
| Barx2 | 99.19 | 1.07 | 0.35 | 9.37 | 0.0022014 | 0.0446178 | 66.37 | 69.90 | 70.38 | 45.20 | 135.97 | 194.90 | 145.32 | 65.49 |
| Hspa8 | 5523.18 | -0.37 | 0.12 | 9.34 | 0.0022393 | 0.04524 | 5262.34 | 5509.87 | 8645.43 | 4912.68 | 5102.00 | 4493.94 | 5539.98 | 4719.20 |
| Heatr5b | 1445.23 | 0.32 | 0.11 | 9.34 | 0.0022449 | 0.0452756 | 1250.04 | 1291.76 | 1431.80 | 1145.07 | 1522.21 | 1678.20 | 1610.95 | 1631.79 |
| B3gnt5 | 209.25 | 0.57 | 0.19 | 9.34 | 0.0022467 | 0.0452756 | 153.77 | 208.75 | 161.38 | 164.79 | 186.96 | 275.48 | 251.66 | 271.20 |
| Padi4 | 1698.93 | 0.53 | 0.17 | 9.33 | 0.0022513 | 0.0453099 | 1371.72 | 1510.09 | 1475.49 | 1236.41 | 1792.02 | 2692.99 | 1956.54 | 1556.15 |
| Alpk1 | 1326.22 | 0.44 | 0.14 | 9.33 | 0.0022579 | 0.0453858 | 1255.57 | 1306.13 | 963.43 | 995.34 | 1660.30 | 1554.51 | 1525.89 | 1348.61 |
| Tmbim6 | 19529.60 | -0.23 | 0.08 | 9.32 | 0.0022666 | 0.0455017 | 19810.36 | 18688.92 | 23840.75 | 20530.26 | 19746.20 | 17273.92 | 19182.58 | 17163.82 |
| Bcap31 | 4955.51 | -0.28 | 0.09 | 9.32 | 0.0022694 | 0.0455017 | 5167.20 | 5262.81 | 6325.42 | 4635.83 | 4651.60 | 4261.56 | 4905.52 | 4434.16 |
| Exoc5 | 1798.49 | 0.27 | 0.09 | 9.29 | 0.0023048 | 0.0460958 | 1816.43 | 1727.46 | 1431.80 | 1724.20 | 1801.58 | 1997.72 | 1694.25 | 2194.48 |
| 2700097O09R | 542.28 | 0.34 | 0.11 | 9.29 | 0.0023023 | 0.0460958 | 474.57 | 542.94 | 440.46 | 496.26 | 554.50 | 645.61 | 529.01 | 654.93 |
| Slc52a3 | 2094.32 | -0.31 | 0.10 | 9.28 | 0.0023167 | 0.0461605 | 2107.37 | 2248.38 | 2778.67 | 1921.95 | 2181.87 | 1724.11 | 1978.69 | 1813.52 |
| Trf | 1238.53 | -0.53 | 0.18 | 9.28 | 0.0023124 | 0.0461605 | 1265.53 | 1358.79 | 1759.42 | 1407.79 | 888.04 | 779.60 | 1192.71 | 1256.36 |
| Otud3 | 289.79 | 0.45 | 0.15 | 9.28 | 0.0023142 | 0.0461605 | 258.86 | 235.56 | 249.96 | 239.18 | 287.87 | 354.19 | 361.53 | 331.16 |
| Grb10 | 2712.56 | 0.45 | 0.15 | 9.27 | 0.0023277 | 0.0463218 | 2310.91 | 2725.25 | 2161.05 | 2191.26 | 2478.23 | 4163.17 | 2847.97 | 2822.66 |
| Htt | 2628.41 | 0.26 | 0.08 | 9.26 | 0.0023452 | 0.0465021 | 2438.13 | 2306.79 | 2370.97 | 2366.41 | 2921.19 | 2736.09 | 3155.45 | 2732.26 |
| Arhgef26 | 695.96 | -0.80 | 0.25 | 9.26 | 0.0023455 | 0.0465021 | 825.25 | 1358.79 | 592.14 | 861.63 | 542.81 | 341.07 | 406.73 | 639.25 |
| Oas3 | 99.49 | -1.68 | 0.52 | 9.26 | 0.0023437 | 0.0465021 | 68.59 | 68.95 | 88.58 | 399.27 | 40.37 | 43.10 | 36.33 | 50.73 |
| Acap2 | 4875.23 | 0.30 | 0.10 | 9.24 | 0.0023647 | 0.0467078 | 4781.13 | 4435.47 | 4333.03 | 3988.91 | 5649.06 | 5532.15 | 4845.27 | 5436.86 |
| Gpa33 | 3514.37 | -0.25 | 0.08 | 9.24 | 0.0023618 | 0.0467078 | 3723.57 | 3507.58 | 4201.98 | 3446.51 | 3842.16 | 2956.29 | 3598.51 | 2838.34 |
| Chka | 1524.24 | 0.30 | 0.10 | 9.24 | 0.0023647 | 0.0467078 | 1496.73 | 1337.73 | 1292.26 | 1323.99 | 1802.64 | 1662.27 | 1666.78 | 1611.50 |
| Pdhb | 2194.47 | -0.42 | 0.14 | 9.24 | 0.0023697 | 0.0467492 | 2474.64 | 2007.07 | 3441.18 | 1846.61 | 2006.59 | 1658.52 | 2279.97 | 1841.19 |
| Mstn | 14.05 | -2.89 | 0.94 | 9.22 | 0.0023917 | 0.0471253 | 5.53 | 38.30 | 33.98 | 19.78 | 0.00 | 5.62 | 0.89 | 8.30 |
| Akap9 | 2831.89 | 0.29 | 0.09 | 9.22 | 0.0023979 | 0.0471878 | 2720.22 | 2674.50 | 2328.50 | 2570.76 | 3282.36 | 3181.17 | 2892.27 | 3005.31 |
| Ndufb9 | 3481.54 | -0.35 | 0.12 | 9.21 | 0.0024027 | 0.0472243 | 3864.06 | 3902.10 | 4408.26 | 3228.04 | 3166.57 | 2682.68 | 3294.57 | 3306.02 |
| Dsel | 906.11 | 0.37 | 0.12 | 9.21 | 0.0024057 | 0.0472243 | 805.34 | 902.03 | 712.26 | 839.97 | 834.93 | 1294.96 | 839.15 | 1020.22 |
| Pla2g4f | 108.39 | -0.77 | 0.26 | 9.21 | 0.00241 | 0.0472522 | 161.51 | 135.02 | 135.90 | 122.42 | 76.48 | 97.45 | 46.08 | 92.24 |
| Ccdc6 | 5328.46 | 0.25 | 0.08 | 9.19 | 0.0024331 | 0.0476461 | 4824.27 | 4902.77 | 5005.25 | 4787.44 | 5742.53 | 5613.67 | 5493.02 | 6258.75 |
| Mcm5 | 319.72 | -0.41 | 0.13 | 9.18 | 0.0024402 | 0.0477257 | 377.22 | 394.52 | 315.48 | 408.68 | 208.20 | 308.28 | 241.91 | 303.48 |
| Fam160b1 | 1951.36 | 0.28 | 0.09 | 9.18 | 0.0024486 | 0.0478325 | 1635.01 | 1832.79 | 1840.72 | 1751.50 | 2006.59 | 2186.06 | 2167.43 | 2190.79 |
| Tomm7 | 2482.27 | -0.35 | 0.12 | 9.17 | 0.0024555 | 0.0479074 | 2966.91 | 2772.17 | 2948.55 | 2297.67 | 2461.24 | 1909.64 | 2282.63 | 2219.39 |
| Kirrel | 2623.26 | 0.48 | 0.16 | 9.17 | 0.0024598 | 0.0479328 | 2227.95 | 2083.68 | 1994.82 | 2694.11 | 2106.45 | 3583.15 | 3256.47 | 3039.44 |
| Ephx2 | 978.74 | -0.38 | 0.13 | 9.16 | 0.002472 | 0.0481121 | 1005.56 | 911.61 | 1260.72 | 1205.34 | 902.91 | 791.78 | 870.16 | 881.85 |
| Gpt | 2009.09 | -0.38 | 0.13 | 9.15 | 0.0024894 | 0.048332 | 2251.18 | 1895.99 | 2889.09 | 1707.25 | 2022.53 | 1400.84 | 2221.49 | 1684.37 |
| Pdzd8 | 1989.43 | 0.31 | 0.10 | 9.15 | 0.0024875 | 0.048332 | 1812.00 | 1921.85 | 1657.50 | 1837.20 | 2249.85 | 2300.38 | 1812.10 | 2324.55 |
| Dclk1 | 661.31 | 0.64 | 0.21 | 9.14 | 0.0024971 | 0.0483779 | 655.99 | 624.34 | 427.11 | 465.18 | 395.16 | 1259.35 | 599.90 | 863.40 |
| Lct | 14.92 | 1.81 | 0.61 | 9.14 | 0.0024978 | 0.0483779 | 9.96 | 7.66 | 7.28 | 2.83 | 12.75 | 31.86 | 22.15 | 24.91 |
| Atp8b5 | 41.08 | 1.30 | 0.41 | 9.12 | 0.0025312 | 0.0489651 | 28.76 | 21.07 | 12.13 | 35.78 | 70.11 | 58.10 | 59.37 | 43.35 |
| Pdgfrl | 324.07 | 0.93 | 0.31 | 9.11 | 0.0025372 | 0.0490218 | 177.00 | 228.86 | 243.89 | 255.19 | 300.62 | 727.13 | 420.02 | 239.83 |
| Pex11a | 398.20 | -0.56 | 0.19 | 9.11 | 0.0025487 | 0.0491847 | 436.96 | 355.26 | 578.79 | 483.08 | 291.06 | 266.11 | 439.51 | 334.85 |
| Rab23 | 1023.89 | 0.34 | 0.11 | 9.10 | 0.002555 | 0.0492465 | 920.38 | 903.95 | 839.67 | 1014.18 | 946.47 | 1203.13 | 1168.78 | 1194.56 |
| Nfatc3 | 2358.41 | 0.25 | 0.08 | 9.10 | 0.002561 | 0.0493024 | 2196.97 | 2242.63 | 2107.66 | 2198.80 | 2367.76 | 2688.30 | 2293.26 | 2771.93 |
| Ugt2b34 | 5758.05 | 0.70 | 0.23 | 9.09 | 0.0025732 | 0.0494782 | 6031.16 | 2620.87 | 4359.72 | 4352.39 | 7660.96 | 8461.27 | 7433.60 | 5144.44 |
| Gdf11 | 262.69 | 0.71 | 0.24 | 9.08 | 0.0025822 | 0.0495916 | 224.56 | 159.91 | 194.14 | 240.13 | 257.06 | 488.19 | 275.58 | 261.97 |
| Gm15816 | 81.30 | -1.09 | 0.36 | 9.08 | 0.0025903 | 0.049686 | 81.86 | 116.82 | 158.95 | 61.21 | 103.04 | 27.17 | 47.85 | 53.50 |

**Table S2:** Comparison of differentially expressed genes from human obstruction (UA, (47)) and mouse NeMO revealed 160 genes in common from 796 human orthologs. Human PBO statistics are shown here.

**Upregulated:**

| Gene | baseMean | log2FoldChange | lfcSE | stat | pvalue | padj | C1 | C2 | C3 | C4 | C5 | C6 | UA1 | UA2 | UA3 | UA4 | UA5 | UA6 |
|---|---|---|---|---|---|---|---|---|---|---|---|---|---|---|---|---|---|---|
| CXCL6* | 106.09 | -6.22 | 1.10 | -5.66 | 1.51E-08 | 3.43E-06 | 5.92 | 0.76 | 3.77 | 3.28 | - | 3.10 | 399.05 | 89.74 | 489.81 | 79.51 | - | 198.13 |
| IGHG3 | 1641.32 | -4.21 | 1.08 | -3.91 | 9.29E-05 | 1.56E-03 | 549.38 | 15.96 | 16.04 | 22.15 | 27.49 | 376.71 | 4,631.61 | 282.86 | 3,834.02 | 3,638.95 | 122.88 | 6,177.77 |
| THBS1 | 7771.33 | -4.04 | 0.65 | -6.17 | 7.01E-10 | 3.57E-07 | 578.22 | 514.40 | 815.06 | 2,309.92 | 677.34 | 462.38 | 18,681.27 | 6,830.63 | 3,768.07 | 47,609.13 | 1,275.60 | 9,733.90 |
| CD80 | 17.85 | -3.74 | 0.73 | -5.15 | 2.65E-07 | 2.41E-05 | 5.92 | 1.52 | 2.83 | 2.46 | 1.96 | - | 17.44 | 26.13 | 57.10 | 25.99 | 1.34 | 71.49 |
| PTX3 | 63.52 | -3.43 | 0.72 | -4.77 | 1.87E-06 | 9.70E-05 | 13.31 | 9.88 | 11.32 | 9.02 | 10.80 | 10.32 | 66.51 | 19.31 | 19.30 | 515.26 | 24.04 | 53.11 |
| ADAMDEC1 | 304.23 | -3.10 | 0.83 | -3.74 | 1.84E-04 | 2.59E-03 | 235.87 | 23.55 | 36.79 | 45.94 | 23.56 | 14.45 | 336.90 | 531.64 | 1,242.62 | 272.16 | 29.39 | 857.88 |
| HTRA4 | 8.14 | -2.85 | 0.61 | -4.65 | 3.38E-06 | 1.49E-04 | 2.96 | 1.52 | 1.89 | 1.64 | 0.98 | 3.10 | 10.90 | 28.40 | 19.30 | 4.59 | 4.01 | 18.38 |
| INHBA | 54.04 | -2.80 | 0.65 | -4.30 | 1.68E-05 | 4.72E-04 | 14.05 | 16.72 | 10.38 | 6.56 | 10.80 | 22.71 | 160.27 | 73.84 | 12.06 | 273.69 | 32.06 | 15.32 |
| BARX2 | 4.54 | -2.67 | 0.82 | -3.27 | 1.09E-03 | 9.54E-03 | 2.22 | 1.52 | - | 2.46 | 0.98 | - | 21.81 | 9.09 | 4.02 | 1.53 | 2.67 | 8.17 |
| GREM1 | 12.86 | -2.49 | 0.78 | -3.19 | 1.43E-03 | 1.16E-02 | 4.44 | 4.56 | 1.89 | 3.28 | - | 9.29 | - | 49.98 | 33.78 | 18.35 | 9.35 | 19.40 |
| SLC7A11 | 100.75 | -2.45 | 0.44 | -5.55 | 2.81E-08 | 5.19E-06 | 24.40 | 27.35 | 28.30 | 57.42 | 33.38 | 15.48 | 164.64 | 94.29 | 228.42 | 331.79 | 36.06 | 167.49 |
| CRLF1 | 47.45 | -2.32 | 0.74 | -3.15 | 1.62E-03 | 1.27E-02 | 17.01 | 29.63 | 11.32 | 6.56 | 13.74 | 16.51 | 299.83 | 7.95 | 6.43 | 45.87 | 26.71 | 87.83 |
| STK17B | 881.05 | -2.31 | 0.42 | -5.52 | 3.36E-08 | 5.75E-06 | 518.33 | 249.22 | 205.65 | 389.63 | 230.69 | 181.65 | 1,103.39 | 2,174.26 | 2,173.98 | 1,227.76 | 268.48 | 1,849.55 |
| B3GALT5 | 185.71 | -1.93 | 0.30 | -6.40 | 1.53E-10 | 1.32E-07 | 93.91 | 71.42 | 53.77 | 77.93 | 55.95 | 111.47 | 452.48 | 354.42 | 319.30 | 201.82 | 121.55 | 314.56 |
| MYBL1 | 65.76 | -1.91 | 0.38 | -4.96 | 6.96E-07 | 4.77E-05 | 39.19 | 28.11 | 27.36 | 17.23 | 25.52 | 28.90 | 130.84 | 84.06 | 189.81 | 93.27 | 22.71 | 102.13 |
| B3GNT5 | 237.17 | -1.83 | 0.53 | -3.47 | 5.28E-04 | 5.61E-03 | 95.38 | 98.02 | 68.86 | 193.59 | 71.66 | 97.02 | 224.60 | 330.57 | 161.66 | 1,230.82 | 100.18 | 173.62 |
| PLA1A | 41.38 | -1.81 | 0.45 | -4.00 | 6.25E-05 | 1.19E-03 | 17.75 | 9.12 | 16.04 | 25.43 | 26.50 | 15.48 | 63.24 | 57.93 | 57.10 | 82.56 | 8.01 | 117.45 |
| UGCG | 1325.40 | -1.73 | 0.39 | -4.44 | 9.08E-06 | 3.07E-04 | 630.72 | 564.54 | 478.28 | 977.78 | 576.23 | 448.96 | 2,303.81 | 1,655.12 | 1,745.30 | 3,892.76 | 396.70 | 2,234.58 |
| BCL2L11 | 420.89 | -1.67 | 0.30 | -5.56 | 2.70E-08 | 5.06E-06 | 301.68 | 217.31 | 178.29 | 178.82 | 126.63 | 207.45 | 412.13 | 665.68 | 852.54 | 750.73 | 265.81 | 893.63 |
| ARID5B | 971.86 | -1.65 | 0.34 | -4.84 | 1.27E-06 | 7.25E-05 | 555.30 | 449.05 | 335.83 | 573.38 | 402.48 | 501.60 | 2,389.95 | 1,158.70 | 867.82 | 2,723.10 | 597.06 | 1,108.10 |
| ADCY7 | 197.63 | -1.63 | 0.36 | -4.54 | 5.56E-06 | 2.15E-04 | 139.01 | 90.42 | 97.17 | 98.43 | 58.90 | 94.95 | 128.66 | 319.21 | 468.90 | 324.14 | 105.52 | 446.30 |
| SLC41A2 | 61.26 | -1.62 | 0.31 | -5.20 | 2.03E-07 | 1.96E-05 | 42.15 | 25.07 | 22.64 | 28.71 | 40.25 | 21.67 | 134.11 | 93.15 | 113.40 | 76.45 | 33.39 | 104.17 |
| CLCA2 | 187.75 | -1.60 | 0.54 | -2.97 | 2.96E-03 | 1.99E-02 | 194.47 | 34.95 | 49.05 | 105.00 | 29.45 | 145.52 | 138.47 | 439.62 | 176.94 | 299.68 | 570.35 | 69.45 |
| TNFRSF10C | 39.53 | -1.54 | 0.55 | -2.82 | 4.75E-03 | 2.79E-02 | 18.49 | 28.87 | 16.04 | 13.94 | 35.34 | 8.26 | 40.34 | 21.58 | 67.56 | 119.26 | 6.68 | 98.04 |
| CHST11 | 322.20 | -1.52 | 0.30 | -5.07 | 4.06E-07 | 3.30E-05 | 218.13 | 164.88 | 157.54 | 165.70 | 159.03 | 135.20 | 358.71 | 387.37 | 646.64 | 512.21 | 173.64 | 787.41 |
| FILIP1L | 1098.77 | -1.50 | 0.40 | -3.78 | 1.54E-04 | 2.27E-03 | 400.76 | 452.09 | 492.43 | 1,040.12 | 617.46 | 442.77 | 3,157.52 | 991.71 | 883.91 | 2,901.99 | 984.41 | 820.09 |
| ZNF365 | 15.11 | -1.47 | 0.52 | -2.84 | 4.54E-03 | 2.69E-02 | 10.35 | 7.60 | 3.77 | 11.48 | 10.80 | 4.13 | 53.42 | 26.13 | 26.54 | 9.17 | 6.68 | 11.23 |
| ABL2 | 344.48 | -1.33 | 0.28 | -4.72 | 2.30E-06 | 1.12E-04 | 214.43 | 210.47 | 175.46 | 199.33 | 146.27 | 230.16 | 721.78 | 336.25 | 365.15 | 840.93 | 260.46 | 433.03 |
| OSMR | 812.83 | -1.28 | 0.21 | -6.19 | 5.92E-10 | 3.15E-07 | 433.30 | 419.42 | 407.53 | 488.89 | 659.67 | 437.61 | 1,492.63 | 1,012.16 | 1,233.77 | 1,414.30 | 655.83 | 1,098.91 |
| AKAP5 | 35.38 | -1.28 | 0.34 | -3.80 | 1.43E-04 | 2.15E-03 | 31.79 | 15.20 | 16.98 | 21.33 | 18.65 | 20.64 | 67.60 | 44.30 | 80.43 | 29.05 | 21.37 | 57.19 |
| CIT | 52.92 | -1.21 | 0.37 | -3.30 | 9.55E-04 | 8.61E-03 | 32.53 | 38.75 | 16.04 | 29.53 | 26.50 | 49.54 | 56.70 | 93.15 | 144.77 | 53.51 | 30.72 | 63.32 |
| C12orf75 | 91.60 | -1.18 | 0.43 | -2.76 | 5.69E-03 | 3.18E-02 | 43.63 | 60.03 | 38.68 | 59.06 | 65.77 | 69.15 | 188.62 | 159.04 | 65.15 | 59.63 | 32.06 | 258.39 |
| BACH1 | 465.95 | -1.14 | 0.20 | -5.63 | 1.85E-08 | 3.93E-06 | 305.38 | 302.41 | 238.67 | 373.23 | 207.13 | 320.98 | 718.51 | 580.48 | 642.62 | 894.45 | 410.06 | 597.45 |
| BDNF | 48.43 | -1.13 | 0.43 | -2.60 | 9.23E-03 | 4.54E-02 | 33.27 | 15.96 | 32.07 | 30.35 | 49.08 | 21.67 | 112.30 | 23.86 | 57.91 | 116.20 | 20.04 | 68.43 |
| PLEKHG2 | 463.95 | -1.11 | 0.22 | -4.99 | 6.14E-07 | 4.30E-05 | 305.38 | 287.97 | 266.97 | 228.04 | 319.04 | 359.17 | 642.19 | 566.85 | 768.90 | 934.20 | 339.27 | 549.45 |
| NEDD9 | 707.60 | -1.10 | 0.31 | -3.56 | 3.75E-04 | 4.33E-03 | 412.59 | 426.26 | 423.56 | 754.66 | 310.20 | 377.74 | 1,092.48 | 815.63 | 653.08 | 1,845.47 | 519.59 | 859.92 |
| HIF1A | 1860.71 | -1.08 | 0.30 | -3.63 | 2.85E-04 | 3.55E-03 | ####### | 1,268.89 | 1,067.87 | 1,567.56 | 1,101.42 | 1,069.24 | 2,175.16 | 1,621.04 | 3,731.88 | 3,773.50 | 952.36 | 2,891.27 |
| CHD1 | 432.03 | -1.03 | 0.25 | -4.16 | 3.17E-05 | 7.42E-04 | 320.91 | 317.60 | 216.03 | 324.83 | 230.69 | 297.24 | 491.73 | 441.90 | 799.46 | 880.69 | 313.89 | 549.45 |
| FOSL2 | 4137.12 | -1.00 | 0.36 | -2.77 | 5.64E-03 | 3.16E-02 | ####### | 3,144.88 | 2,530.07 | 3,767.56 | 2,121.36 | 2,334.59 | 5,275.98 | 3,812.34 | 2,842.34 | 13,887.65 | 3,386.01 | 3,856.38 |
| DCLK1 | 163.11 | -0.97 | 0.26 | -3.68 | 2.34E-04 | 3.08E-03 | 119.78 | 111.69 | 69.81 | 84.49 | 95.22 | 179.58 | 309.65 | 209.02 | 155.23 | 279.80 | 142.92 | 200.17 |
| BDKRB2 | 735.98 | -0.96 | 0.36 | -2.66 | 7.78E-03 | 4.00E-02 | 429.60 | 479.44 | 532.05 | 626.70 | 485.92 | 440.70 | 1,404.31 | 418.04 | 863.80 | 1,582.49 | 281.83 | 1,286.82 |
| COL4A1 | 1724.74 | -0.93 | 0.28 | -3.38 | 7.22E-04 | 7.03E-03 | 882.86 | 705.87 | 1,149.94 | 1,077.03 | 1,881.84 | 1,417.06 | 3,406.11 | 1,901.63 | 2,016.34 | 2,495.28 | 2,544.52 | 1,218.40 |
| SYNJ2 | 389.16 | -0.91 | 0.22 | -4.11 | 3.94E-05 | 8.55E-04 | 304.64 | 276.57 | 272.63 | 273.97 | 262.10 | 227.06 | 577.86 | 423.72 | 513.13 | 850.11 | 324.58 | 363.58 |
| PIKFYVE | 544.55 | -0.91 | 0.27 | -3.37 | 7.64E-04 | 7.29E-03 | 477.66 | 454.37 | 282.06 | 397.84 | 254.25 | 403.55 | 406.68 | 652.05 | 1,228.95 | 658.99 | 488.87 | 829.29 |
| NIN | 600.02 | -0.87 | 0.24 | -3.54 | 4.02E-04 | 4.57E-03 | 598.92 | 472.61 | 327.34 | 451.16 | 324.93 | 376.71 | 552.78 | 756.56 | 1,138.06 | 631.47 | 507.57 | 1,062.14 |
| F2R | 334.11 | -0.86 | 0.31 | -2.74 | 6.23E-03 | 3.39E-02 | 303.16 | 282.65 | 120.75 | 237.88 | 208.11 | 271.44 | 726.14 | 324.89 | 413.40 | 483.16 | 166.96 | 470.81 |
| SULF1 | 413.36 | -0.86 | 0.31 | -2.81 | 4.94E-03 | 2.87E-02 | 233.65 | 368.51 | 205.65 | 221.48 | 161.97 | 571.78 | 773.03 | 650.92 | 457.64 | 444.93 | 475.51 | 395.24 |
| TNFRSF10B | 1049.33 | -0.86 | 0.25 | -3.40 | 6.78E-04 | 6.70E-03 | 738.67 | 695.99 | 765.06 | 1,024.53 | 641.02 | 609.96 | 1,419.58 | 1,137.11 | 1,429.21 | 2,287.34 | 677.20 | 1,166.31 |
| MSN | 3925.15 | -0.83 | 0.18 | -4.66 | 3.15E-06 | 1.41E-04 | ####### | 2,305.29 | 2,708.36 | 2,640.49 | 3,619.38 | 2,896.04 | 4,799.52 | 4,997.16 | 6,312.03 | 5,583.81 | 3,022.70 | 5,422.02 |
| AKAP13 | 1754.47 | -0.83 | 0.17 | -4.96 | 7.05E-07 | 4.81E-05 | ####### | 1,392.75 | 1,047.12 | 1,173.83 | 1,004.24 | 1,356.16 | 2,025.78 | 1,928.89 | 2,849.58 | 2,389.78 | 1,732.41 | 2,533.82 |
| CBL | 561.36 | -0.81 | 0.17 | -4.72 | 2.32E-06 | 1.13E-04 | 441.43 | 483.24 | 365.08 | 392.10 | 326.89 | 434.51 | 485.19 | 767.92 | 952.27 | 764.49 | 597.06 | 726.14 |
| RDH10 | 650.35 | -0.79 | 0.27 | -2.91 | 3.59E-03 | 2.27E-02 | 400.02 | 474.13 | 519.79 | 550.41 | 514.39 | 397.35 | 665.09 | 465.75 | 875.87 | 1,343.97 | 447.46 | 1,149.97 |
| RASA2 | 363.87 | -0.78 | 0.20 | -3.94 | 8.01E-05 | 1.41E-03 | 340.87 | 278.09 | 188.67 | 331.39 | 226.76 | 244.61 | 402.32 | 435.08 | 635.38 | 461.75 | 327.25 | 494.30 |
| ROBO1 | 737.85 | -0.77 | 0.20 | -3.92 | 8.99E-05 | 1.52E-03 | 617.41 | 598.74 | 340.55 | 604.55 | 463.34 | 646.09 | 814.46 | 1,314.33 | 1,036.72 | 824.12 | 731.97 | 861.97 |
| C1GALT1 | 246.98 | -0.76 | 0.22 | -3.54 | 4.01E-04 | 4.57E-03 | 187.07 | 183.12 | 154.71 | 232.96 | 197.31 | 145.52 | 393.60 | 337.39 | 387.67 | 258.40 | 164.29 | 321.71 |
| SLC39A14 | 447.19 | -0.76 | 0.23 | -3.23 | 1.23E-03 | 1.04E-02 | 270.62 | 265.18 | 385.83 | 361.75 | 460.40 | 249.77 | 617.11 | 423.72 | 481.77 | 758.37 | 351.29 | 740.43 |
| CTHRC1 | 173.58 | -0.75 | 0.26 | -2.89 | 3.89E-03 | 2.41E-02 | 176.72 | 123.85 | 117.92 | 72.18 | 154.12 | 130.04 | 237.69 | 152.22 | 256.57 | 330.26 | 132.23 | 199.15 |
| LDLRAD4 | 331.51 | -0.75 | 0.25 | -2.95 | 3.19E-03 | 2.09E-02 | 389.67 | 277.33 | 192.44 | 270.69 | 129.58 | 223.96 | 380.52 | 401.00 | 511.52 | 457.16 | 259.13 | 485.11 |
| CTSK | 2133.69 | -0.73 | 0.25 | -2.98 | 2.85E-03 | 1.93E-02 | ####### | 1,676.92 | 1,246.17 | 1,090.98 | 1,445.98 | 2,083.79 | 2,898.03 | 2,309.44 | 2,854.41 | 3,427.96 | 1,307.66 | 3,187.44 |
| TGFBR1 | 720.18 | -0.72 | 0.20 | -3.54 | 3.95E-04 | 4.51E-03 | 527.20 | 522.75 | 407.53 | 646.38 | 674.40 | 480.95 | 1,116.47 | 719.07 | 863.00 | 1,304.21 | 661.17 | 718.99 |
| MICAL2 | 798.63 | -0.72 | 0.22 | -3.22 | 1.30E-03 | 1.08E-02 | 447.34 | 489.32 | 616.01 | 604.55 | 938.47 | 530.49 | 1,341.07 | 721.35 | 1,152.54 | 1,038.17 | 1,009.79 | 694.48 |
| JAK2 | 313.88 | -0.69 | 0.21 | -3.27 | 1.08E-03 | 9.47E-03 | 285.41 | 256.06 | 181.12 | 285.46 | 149.21 | 284.86 | 342.36 | 346.47 | 474.53 | 441.87 | 265.81 | 453.45 |
| GATAD2B | 44.87 | -0.69 | 0.24 | -2.80 | 5.04E-03 | 2.91E-02 | 31.79 | 47.11 | 32.07 | 43.48 | 22.58 | 28.90 | 34.89 | 77.25 | 61.13 | 47.40 | 58.77 | 53.11 |
| DIP2B | 466.03 | -0.67 | 0.24 | -2.82 | 4.87E-03 | 2.84E-02 | 391.15 | 401.94 | 288.67 | 414.24 | 240.51 | 425.22 | 360.89 | 561.17 | 780.16 | 733.91 | 363.31 | 631.16 |
| C21orf91 | 205.67 | -0.66 | 0.23 | -2.91 | 3.62E-03 | 2.29E-02 | 198.90 | 161.84 | 116.98 | 182.10 | 147.25 | 152.75 | 285.66 | 205.61 | 373.19 | 171.24 | 172.31 | 300.26 |
| PCNX1 | 808.92 | -0.66 | 0.17 | -3.85 | 1.19E-04 | 1.87E-03 | 764.55 | 684.60 | 441.49 | 615.21 | 527.15 | 736.91 | 807.92 | 906.51 | 1,162.19 | 1,016.77 | 833.48 | 1,210.23 |
| PXDN | 806.20 | -0.65 | 0.13 | -4.81 | 1.48E-06 | 8.15E-05 | 550.12 | 538.71 | 655.63 | 556.97 | 727.41 | 745.17 | 1,091.39 | 808.82 | 1,036.72 | 1,149.79 | 918.97 | 894.65 |
| KRAS | 428.90 | -0.64 | 0.17 | -3.84 | 1.24E-04 | 1.92E-03 | 340.87 | 336.60 | 335.83 | 375.69 | 352.42 | 268.34 | 611.66 | 583.89 | 654.69 | 452.58 | 337.93 | 496.35 |
| RAD54L2 | 416.60 | -0.61 | 0.10 | -5.91 | 3.45E-09 | 1.15E-06 | 306.86 | 360.91 | 287.72 | 348.62 | 305.30 | 373.62 | 444.84 | 479.38 | 515.55 | 498.44 | 518.25 | 559.67 |
| HIPK2 | 635.72 | -0.60 | 0.22 | -2.69 | 7.15E-03 | 3.75E-02 | 640.33 | 684.60 | 349.04 | 481.51 | 305.30 | 575.91 | 577.86 | 677.04 | 943.43 | 782.83 | 707.92 | 902.82 |
| APAF1 | 273.50 | -0.60 | 0.22 | -2.76 | 5.83E-03 | 3.23E-02 | 231.44 | 255.30 | 151.88 | 260.85 | 153.14 | 252.86 | 227.87 | 336.25 | 385.25 | 461.75 | 297.86 | 267.58 |
| RUNX1 | 787.55 | -0.59 | 0.16 | -3.81 | 1.38E-04 | 2.09E-03 | 735.72 | 697.51 | 553.75 | 733.33 | 537.95 | 511.92 | 1,108.84 | 857.66 | 974.79 | 931.14 | 699.91 | 1,108.10 |
| CDK12 | 596.00 | -0.58 | 0.13 | -4.39 | 1.16E-05 | 3.61E-04 | 520.55 | 514.40 | 385.83 | 502.83 | 399.54 | 541.85 | 613.84 | 687.27 | 875.06 | 736.96 | 634.46 | 739.41 |
| MED13L | 845.59 | -0.58 | 0.21 | -2.78 | 5.46E-03 | 3.09E-02 | 764.55 | 805.41 | 487.71 | 776.81 | 453.53 | 785.42 | 609.48 | 1,048.51 | 1,193.56 | 1,085.57 | 1,067.23 | 1,069.29 |
| FAM160B1 | 342.11 | -0.57 | 0.21 | -2.70 | 6.88E-03 | 3.65E-02 | 349.00 | 278.09 | 225.46 | 319.91 | 195.35 | 282.79 | 347.81 | 408.95 | 509.11 | 236.99 | 434.10 | 517.79 |
| ITSN2 | 676.37 | -0.57 | 0.19 | -2.95 | 3.14E-03 | 2.07E-02 | 673.60 | 665.60 | 462.24 | 576.66 | 378.92 | 509.85 | 576.77 | 839.49 | 1,105.89 | 746.14 | 699.91 | 881.37 |
| VASH1 | 331.60 | -0.54 | 0.18 | -2.97 | 2.99E-03 | 2.00E-02 | 334.21 | 230.98 | 289.61 | 236.24 | 231.67 | 303.43 | 375.06 | 424.86 | 534.04 | 380.71 | 256.46 | 381.96 |
| RNF19A | 962.22 | -0.53 | 0.14 | -3.67 | 2.46E-04 | 3.20E-03 | 916.87 | 820.60 | 663.18 | 963.83 | 575.25 | 784.39 | 1,097.94 | 1,108.71 | 1,216.08 | 1,334.79 | 997.77 | 1,067.25 |
| TET3 | 378.22 | -0.53 | 0.20 | -2.65 | 8.11E-03 | 4.12E-02 | 326.82 | 368.51 | 220.74 | 379.79 | 222.84 | 338.53 | 299.83 | 480.52 | 460.86 | 593.24 | 422.08 | 424.86 |
| B4GALT5 | 1130.94 | -0.52 | 0.19 | -2.70 | 6.93E-03 | 3.66E-02 | 808.92 | 1,031.83 | 970.71 | 1,063.09 | 867.79 | 834.96 | 1,393.41 | 1,259.80 | 1,213.67 | 2,097.75 | 885.57 | 1,143.84 |
| LPIN2 | 500.34 | -0.51 | 0.12 | -4.38 | 1.18E-05 | 3.68E-04 | 428.12 | 414.10 | 388.66 | 444.59 | 384.81 | 421.09 | 586.58 | 556.63 | 706.97 | 617.70 | 446.13 | 608.69 |
| SEMA3C | 732.27 | -0.50 | 0.19 | -2.62 | 8.77E-03 | 4.37E-02 | 521.29 | 536.43 | 569.78 | 768.61 | 469.23 | 770.97 | 1,053.23 | 886.06 | 857.37 | 795.07 | 1,013.80 | 545.37 |
| IL13RA1 | 925.61 | -0.50 | 0.17 | -2.90 | 3.72E-03 | 2.33E-02 | 920.57 | 754.50 | 752.79 | 794.03 | 686.18 | 697.69 | 1,059.78 | 1,222.31 | 1,217.69 | 986.19 | 653.16 | 1,362.40 |
| EDEM3 | 508.34 | -0.49 | 0.17 | -2.91 | 3.60E-03 | 2.28E-02 | 510.93 | 442.97 | 288.67 | 490.53 | 342.60 | 456.18 | 511.35 | 622.52 | 718.23 | 651.34 | 475.51 | 589.28 |
| ATAD2B | 226.94 | -0.49 | 0.18 | -2.68 | 7.29E-03 | 3.80E-02 | 226.26 | 221.11 | 150.94 | 177.18 | 158.05 | 200.23 | 216.97 | 218.11 | 385.25 | 255.34 | 235.08 | 278.81 |
| SENP5 | 468.15 | -0.48 | 0.12 | -3.94 | 8.18E-05 | 1.44E-03 | 378.58 | 429.30 | 346.21 | 397.84 | 376.96 | 414.90 | 536.43 | 497.56 | 534.04 | 712.50 | 450.13 | 543.33 |
| PDZD8 | 317.23 | -0.46 | 0.14 | -3.30 | 9.59E-04 | 8.64E-03 | 268.41 | 301.65 | 199.99 | 295.30 | 255.23 | 278.66 | 423.04 | 316.94 | 402.95 | 376.13 | 395.37 | 293.11 |
| KAT6A | 720.49 | -0.44 | 0.17 | -2.56 | 1.04E-02 | 4.95E-02 | 717.23 | 740.06 | 457.53 | 592.24 | 459.42 | 699.76 | 612.75 | 792.91 | 925.73 | 981.60 | 876.22 | 790.48 |
| TULP4 | 310.31 | -0.44 | 0.16 | -2.74 | 6.14E-03 | 3.35E-02 | 357.14 | 310.77 | 218.86 | 236.24 | 222.84 | 233.25 | 318.37 | 408.95 | 377.21 | 288.98 | 379.34 | 371.75 |
| TMF1 | 366.75 | -0.44 | 0.14 | -3.15 | 1.64E-03 | 1.28E-02 | 353.44 | 351.04 | 233.95 | 328.11 | 282.72 | 319.95 | 450.30 | 372.60 | 501.87 | 431.17 | 345.95 | 429.96 |
| WDR26 | 1498.12 | -0.43 | 0.13 | -3.35 | 7.97E-04 | 7.53E-03 | ####### | 1,357.79 | 1,041.46 | 1,513.42 | 1,083.75 | 1,279.79 | 1,725.95 | 1,694.88 | 1,977.74 | 1,885.22 | 1,371.77 | 1,671.85 |
| AP4E1 | 185.65 | -0.39 | 0.15 | -2.58 | 9.79E-03 | 4.74E-02 | 201.12 | 177.80 | 141.50 | 165.70 | 146.27 | 134.17 | 196.25 | 232.88 | 256.57 | 177.36 | 181.66 | 216.51 |
| NUP98 | 1218.94 | -0.37 | 0.12 | -3.09 | 1.97E-03 | 1.47E-02 | ####### | 1,067.54 | 1,010.33 | 1,113.95 | 1,120.07 | 1,042.41 | 1,433.75 | 1,098.49 | 1,631.89 | 1,605.42 | 1,123.33 | 1,363.42 |
| KIF5B | 1412.96 | -0.37 | 0.14 | -2.62 | 8.82E-03 | 4.39E-02 | ####### | 1,253.70 | 1,070.70 | 1,554.44 | 1,272.23 | 1,116.72 | 1,694.33 | 1,444.96 | 1,648.78 | 1,877.58 | 1,720.39 | 1,175.50 |
| ZZEF1 | 908.67 | -0.37 | 0.13 | -2.95 | 3.17E-03 | 2.08E-02 | 834.06 | 847.96 | 706.57 | 835.05 | 620.41 | 914.43 | 827.54 | 1,015.56 | 1,172.65 | 1,007.59 | 1,032.50 | 1,089.71 |
| HTT | 1082.75 | -0.36 | 0.11 | -3.18 | 1.50E-03 | 1.20E-02 | ####### | 977.13 | 944.29 | 906.41 | 833.43 | 1,008.35 | 963.83 | 1,162.10 | 1,470.23 | 1,273.63 | 1,148.71 | 1,283.76 |
| EXOC5 | 481.43 | -0.36 | 0.13 | -2.80 | 5.11E-03 | 2.94E-02 | 459.91 | 419.42 | 348.10 | 487.25 | 426.04 | 394.26 | 585.49 | 605.48 | 629.75 | 490.80 | 460.82 | 469.79 |
| AGPS | 458.87 | -0.34 | 0.12 | -2.81 | 4.98E-03 | 2.89E-02 | 403.72 | 368.51 | 342.44 | 513.50 | 439.78 | 359.17 | 574.59 | 525.96 | 526.81 | 466.34 | 468.83 | 516.77 |
| LARP4B | 741.58 | -0.27 | 0.09 | -3.09 | 2.00E-03 | 1.48E-02 | 678.78 | 734.74 | 703.74 | 666.07 | 619.43 | 638.86 | 865.70 | 831.54 | 865.41 | 759.90 | 693.23 | 841.54 |
| INO80 | 467.93 | -0.22 | 0.08 | -2.65 | 8.04E-03 | 4.09E-02 | 411.85 | 450.57 | 389.60 | 451.98 | 455.49 | 440.70 | 477.55 | 521.41 | 541.28 | 455.63 | 498.22 | 520.86 |

* CXCL6 is the human homologue of mouse CXCL5, which is differentially expressed.

**Downreulated:**

| Gene | baseMea | log2FoldChan | lfcSE | stat | pvalue | padj | C1_star | C2_starAligr | C3_starAligr | C4_starAligr | C5_starAligr | C6_starAligr | UA1_starAli | UA2_starA | UA3_starA | UA4_starAli | UA5_starAli | UA6_starAligned.out.bam |
|---|---|---|---|---|---|---|---|---|---|---|---|---|---|---|---|---|---|---|
| KRT20 | 110.56 | 2.953017 | 1 | 3 | 0.0025 | 0.0172 | ##### | 107.89 | 479.22 | 321.55 | 88.35 | 45.41 | 8.72 | 4.54 | 0.80 | 65.75 | 70.79 | 2.04 |
| UPK3A | 1349.4 | 2.388926 | 0.6 | 4 | ####### | 0.0011 | ##### | 1,328.92 | 2,547.99 | 2,349.29 | 5,064.38 | 975.32 | 281.30 | 441.90 | 94.10 | 117.73 | 1,427.87 | 233.88 |
| RBP4 | 103.21 | 2.294362 | 0.6 | 3.8 | 0.0001 | 0.0022 | 48.80 | 261.38 | 199.99 | 150.11 | 219.89 | 147.59 | 7.63 | 22.72 | 22.52 | 27.52 | 124.22 | 6.13 |
| UPK2 | 1521.1 | 2.288727 | 0.8 | 2.9 | 0.0033 | 0.0214 | ##### | 1,314.48 | 2,810.24 | 2,711.86 | 4,570.60 | 1,468.66 | 164.64 | 316.94 | 38.61 | 134.55 | 2,134.46 | 312.51 |
| UPK1A | 4062 | 2.137364 | 0.5 | 4 | ####### | 0.0013 | ##### | 4,163.80 | 7,613.79 | 7,395.68 | 10,402.64 | 3,959.09 | 1,212.42 | 1,242.76 | 200.27 | 694.15 | 4,761.79 | 917.12 |
| CA4 | 34.32 | 2.078731 | 0.6 | 3.8 | 0.0002 | 0.0025 | 22.18 | 94.22 | 72.64 | 63.98 | 43.19 | 35.09 | 8.72 | 4.54 | 4.83 | 25.99 | 33.39 | 3.06 |
| SOSTDC1 | 92.334 | 1.843335 | 0.4 | 4.3 | ####### | 0.0004 | ##### | 291.77 | 99.05 | 69.72 | 70.68 | 118.69 | 27.26 | 35.22 | 15.28 | 47.40 | 70.79 | 46.98 |
| REEP6 | 293.93 | 1.821039 | 0.4 | 5.1 | ####### | ####### | ##### | 351.80 | 572.61 | 425.73 | 670.47 | 348.85 | 88.31 | 137.45 | 57.10 | 114.67 | 296.53 | 85.79 |
| SFRP5 | 9.1474 | 1.801142 | 0.6 | 3.1 | 0.0018 | 0.0136 | 8.13 | 17.48 | 17.92 | 9.84 | 9.82 | 20.64 | 3.27 | 3.41 | 1.61 | 15.29 | 1.34 | 1.02 |
| SPOCK3 | 109.36 | 1.73706 | 0.5 | 3.6 | 0.0003 | 0.0041 | ##### | 334.32 | 99.05 | 67.26 | 130.56 | 216.74 | 65.42 | 11.36 | 31.37 | 25.99 | 110.86 | 58.21 |
| LGI1 | 24.234 | 1.712562 | 0.4 | 4.3 | ####### | 0.0005 | 43.63 | 50.91 | 28.30 | 29.53 | 15.71 | 53.67 | 7.63 | 11.36 | 5.63 | 9.17 | 24.04 | 11.23 |
| PSCA | 5543.2 | 1.680823 | 0.6 | 2.9 | 0.0036 | 0.0227 | ##### | 6,568.62 | 11,671.14 | 7,383.37 | 15,078.28 | 4,274.91 | 7,239.62 | 727.03 | 2,566.47 | 4,036.49 | 804.09 | 441.20 |
| SLC9A2 | 86.407 | 1.675982 | 0.4 | 4.4 | ####### | 0.0004 | ##### | 95.74 | 138.67 | 146.83 | 151.18 | 100.11 | 34.89 | 29.54 | 16.09 | 38.22 | 106.86 | 23.49 |
| ENHO | 4.9703 | 1.557827 | 0.6 | 2.8 | 0.0047 | 0.0277 | 4.44 | 9.88 | 9.43 | 9.02 | 4.91 | 7.22 | 4.36 | 1.14 | 4.02 | 1.53 | 2.67 | 1.02 |
| PAPPA2 | 19.715 | 1.345093 | 0.5 | 2.9 | 0.0033 | 0.0214 | 28.10 | 43.31 | 14.15 | 47.58 | 16.69 | 18.58 | 9.81 | 3.41 | 4.83 | 16.82 | 20.04 | 13.28 |
| ST6GAL2 | 31.056 | 1.344303 | 0.5 | 2.7 | 0.0072 | 0.0379 | 28.84 | 55.47 | 27.36 | 45.94 | 57.92 | 50.57 | 22.90 | 6.82 | 1.61 | 22.93 | 40.07 | 12.26 |
| KAZALD1 | 84.692 | 1.330487 | 0.4 | 3.8 | 0.0002 | 0.0024 | ##### | 149.68 | 125.47 | 84.49 | 81.48 | 156.88 | 46.88 | 30.67 | 18.50 | 103.97 | 56.10 | 35.75 |
| UPK3B | 415.09 | 1.307964 | 0.4 | 3.2 | 0.0013 | 0.0108 | ##### | 335.08 | 867.88 | 702.98 | 938.47 | 298.27 | 111.21 | 389.64 | 108.58 | 388.36 | 152.27 | 283.92 |
| SP6 | 383.04 | 1.303503 | 0.3 | 4.2 | ####### | 0.0006 | ##### | 576.70 | 583.93 | 492.17 | 418.19 | 464.44 | 201.71 | 212.43 | 316.89 | 94.80 | 379.34 | 119.49 |
| FABP4 | 1248.7 | 1.280334 | 0.5 | 2.7 | 0.0076 | 0.0393 | ##### | 1,492.28 | 3,726.24 | 1,868.61 | 2,081.12 | 685.31 | 1,232.04 | 234.01 | 282.30 | 1,322.56 | 681.21 | 618.90 |
| MGST1 | 1960.2 | 1.264262 | 0.2 | 6.5 | ####### | ####### | ##### | 2,530.19 | 3,069.66 | 2,903.81 | 3,561.46 | 1,976.45 | 1,281.11 | 1,018.97 | 844.50 | 925.03 | 1,668.30 | 1,177.55 |
| CLEC3B | 135.68 | 1.232216 | 0.4 | 3.4 | 0.0007 | 0.0066 | ##### | 274.29 | 243.38 | 90.23 | 155.10 | 166.17 | 71.96 | 72.70 | 45.04 | 119.26 | 145.59 | 33.70 |
| DGAT2 | 312.35 | 1.115446 | 0.3 | 4 | ####### | 0.001 | ##### | 450.57 | 599.03 | 445.41 | 422.11 | 277.63 | 110.12 | 124.96 | 158.44 | 304.27 | 277.83 | 210.39 |
| ASTN2 | 113.6 | 1.033153 | 0.3 | 3.7 | 0.0002 | 0.0031 | ##### | 148.16 | 149.05 | 169.80 | 147.25 | 183.71 | 61.06 | 94.29 | 45.04 | 64.22 | 141.58 | 42.89 |
| CPE | 938.89 | 0.98176 | 0.3 | 3.2 | 0.0012 | 0.0101 | ##### | 1,211.15 | 1,164.10 | 1,066.37 | 1,309.53 | 1,878.40 | 437.21 | 600.93 | 468.09 | 773.66 | 1,187.44 | 321.71 |
| EPHX2 | 305.44 | 0.964109 | 0.3 | 3.8 | 0.0002 | 0.0022 | ##### | 408.78 | 454.70 | 375.69 | 332.78 | 524.30 | 194.07 | 249.92 | 121.45 | 177.36 | 357.97 | 142.98 |
| GSN | 13193 | 0.897375 | 0.3 | 2.9 | 0.0041 | 0.025 | ##### | 16,940.89 | 19,094.37 | 12,199.26 | 17,110.31 | 20,942.12 | 12,579.93 | 6,893.11 | 4,963.24 | 9,372.60 | 17,155.80 | 4,342.52 |
| PEX11A | 250.72 | 0.861792 | 0.2 | 3.5 | 0.0005 | 0.0057 | ##### | 307.73 | 371.68 | 315.81 | 271.92 | 332.33 | 147.19 | 165.85 | 135.12 | 114.67 | 349.95 | 156.26 |
| PLA2G4F | 214.83 | 0.797011 | 0.2 | 3.4 | 0.0008 | 0.0074 | ##### | 285.69 | 353.76 | 269.87 | 300.39 | 231.19 | 223.51 | 131.77 | 122.25 | 137.61 | 228.41 | 99.06 |
| LNX1 | 113.1 | 0.753489 | 0.3 | 2.9 | 0.0032 | 0.0209 | ##### | 152.72 | 145.28 | 167.34 | 118.78 | 142.43 | 53.42 | 92.01 | 70.78 | 82.56 | 156.28 | 52.09 |
| ATP5ME | 364.66 | 0.733723 | 0.2 | 3.1 | 0.0016 | 0.0128 | ##### | 348.76 | 583.93 | 454.44 | 703.85 | 294.15 | 321.64 | 202.20 | 255.76 | 243.11 | 285.84 | 333.96 |
| BEND5 | 55.935 | 0.718253 | 0.2 | 3.3 | 0.0009 | 0.0082 | 60.63 | 74.46 | 70.75 | 59.88 | 77.55 | 75.34 | 31.62 | 45.44 | 40.21 | 21.41 | 58.77 | 55.15 |
| TST | 364.64 | 0.712457 | 0.2 | 3.7 | 0.0002 | 0.003 | ##### | 325.20 | 613.18 | 434.75 | 598.81 | 348.85 | 315.10 | 290.81 | 242.09 | 227.82 | 256.46 | 324.77 |
| ALDH1L1 | 1333 | 0.709911 | 0.2 | 3.8 | 0.0002 | 0.0025 | ##### | 1,226.35 | 1,929.15 | 1,491.28 | 2,256.83 | 1,558.45 | 938.75 | 1,210.95 | 728.68 | 1,033.59 | 1,294.30 | 864.01 |
| PXMP4 | 257.84 | 0.709172 | 0.2 | 3.2 | 0.0014 | 0.0116 | ##### | 311.53 | 350.93 | 298.58 | 336.71 | 320.98 | 186.44 | 235.15 | 151.21 | 103.97 | 317.90 | 179.75 |
| HADH | 763.94 | 0.682215 | 0.2 | 4.3 | ####### | 0.0004 | ##### | 797.81 | 1,153.72 | 1,023.71 | 1,095.53 | 754.46 | 570.23 | 706.58 | 521.18 | 492.33 | 706.59 | 522.90 |
| FAAH | 352.2 | 0.668057 | 0.2 | 3.2 | 0.0012 | 0.0104 | ##### | 398.90 | 483.00 | 429.01 | 450.58 | 476.83 | 357.62 | 316.94 | 166.49 | 252.28 | 352.63 | 187.92 |
| QPCT | 331.36 | 0.667145 | 0.2 | 3.3 | 0.0012 | 0.0099 | ##### | 361.67 | 648.08 | 435.57 | 385.79 | 342.65 | 285.66 | 197.66 | 267.02 | 247.69 | 257.79 | 279.83 |
| SYT7 | 88.96 | 0.66142 | 0.2 | 2.8 | 0.0058 | 0.0321 | ##### | 121.57 | 100.94 | 111.56 | 69.70 | 122.82 | 79.59 | 45.44 | 65.15 | 77.98 | 104.19 | 42.89 |
| DBI | 772.38 | 0.620488 | 0.2 | 3.1 | 0.0022 | 0.0157 | ##### | 770.45 | 1,219.75 | 949.07 | 1,225.11 | 655.38 | 512.44 | 614.56 | 752.01 | 458.69 | 547.64 | 764.94 |
| RPL39 | 396.33 | 0.616504 | 0.2 | 3.8 | 0.0001 | 0.0019 | ##### | 392.07 | 551.86 | 470.02 | 596.85 | 420.06 | 282.39 | 273.77 | 337.00 | 259.93 | 297.86 | 423.83 |
| ACTR3B | 57.96 | 0.59685 | 0.2 | 3.2 | 0.0014 | 0.0113 | 62.85 | 64.58 | 83.96 | 62.34 | 84.42 | 59.86 | 47.97 | 35.22 | 40.21 | 44.34 | 62.78 | 46.98 |
| FKBP4 | 847.13 | 0.594996 | 0.2 | 3.5 | 0.0004 | 0.0047 | ##### | 843.40 | 1,235.79 | 963.01 | 1,215.29 | 841.15 | 698.88 | 637.28 | 517.16 | 839.41 | 821.46 | 538.22 |
| MGST3 | 755.92 | 0.579976 | 0.2 | 3 | 0.0031 | 0.0204 | ##### | 696.75 | 1,148.06 | 855.56 | 1,239.84 | 710.08 | 573.50 | 538.45 | 455.23 | 698.74 | 794.74 | 577.03 |
| SOD1 | 1173 | 0.57003 | 0.2 | 3.6 | 0.0004 | 0.0042 | ##### | 1,192.15 | 1,813.12 | 1,359.21 | 1,751.28 | 1,038.28 | 981.27 | 908.78 | 830.02 | 837.88 | 1,107.30 | 999.84 |
| NDN | 460.68 | 0.557219 | 0.2 | 3.2 | 0.0015 | 0.0121 | ##### | 520.47 | 584.88 | 455.26 | 686.18 | 616.16 | 486.28 | 337.39 | 263.00 | 400.59 | 407.39 | 343.15 |
| BCAP31 | 2337.8 | 0.518789 | 0.1 | 3.5 | 0.0004 | 0.0046 | ##### | 2,509.68 | 3,347.95 | 2,601.12 | 3,629.19 | 2,127.14 | 2,027.97 | 1,984.55 | 1,678.54 | 1,989.19 | 1,954.14 | 1,897.55 |
| TOMM7 | 1643.5 | 0.517108 | 0.2 | 3 | 0.0025 | 0.0174 | ##### | 1,753.66 | 2,285.74 | 1,849.74 | 2,474.76 | 1,523.36 | 1,378.14 | 1,028.06 | 1,097.85 | 1,383.72 | 1,457.26 | 1,767.85 |
| RPL32 | 8243.4 | 0.508951 | 0.1 | 3.5 | 0.0004 | 0.0045 | ##### | 8,523.63 | 11,764.53 | 9,366.00 | 12,325.71 | 8,272.19 | 6,175.48 | 6,261.51 | 6,645.00 | 7,099.02 | 6,492.86 | 8,150.90 |
| PLIN2 | 889.47 | 0.501545 | 0.2 | 3 | 0.003 | 0.0199 | ##### | 918.62 | 1,278.24 | 1,066.37 | 1,129.89 | 814.32 | 632.38 | 639.56 | 563.80 | 1,007.59 | 742.65 | 835.41 |
| RPL7A | 11523 | 0.489893 | 0.1 | 4 | ####### | 0.0014 | ##### | 11,982.32 | 15,676.61 | 12,949.82 | 16,892.38 | 12,203.42 | 9,002.64 | 9,217.32 | 9,049.81 | 10,742.56 | 9,315.21 | 10,185.31 |
| HSP90AB1 | 12796 | 0.48563 | 0.1 | 4.2 | ####### | 0.0006 | ##### | 14,738.18 | 16,551.09 | 15,350.79 | 15,349.22 | 14,463.69 | 10,027.52 | 9,458.15 | 9,628.09 | 13,156.80 | 12,275.13 | 9,435.68 |
| SKP1 | 1938 | 0.479314 | 0.1 | 3.6 | 0.0003 | 0.004 | ##### | 1,940.57 | 2,734.77 | 2,400.15 | 2,731.96 | 1,844.34 | 1,797.91 | 1,540.39 | 1,499.19 | 1,603.89 | 1,719.05 | 1,553.38 |
| BANF1 | 638.79 | 0.475715 | 0.2 | 2.6 | 0.0087 | 0.0433 | ##### | 544.79 | 905.62 | 717.75 | 1,084.73 | 639.90 | 542.97 | 485.06 | 555.76 | 487.74 | 540.96 | 592.35 |
| COX4I1 | 2928.7 | 0.473986 | 0.2 | 3 | 0.003 | 0.02 | ##### | 2,868.31 | 4,691.28 | 3,265.55 | 4,179.90 | 2,681.37 | 2,548.04 | 2,370.79 | 2,293.02 | 2,342.39 | 2,384.23 | 2,771.78 |
| HIBADH | 563.03 | 0.473414 | 0.1 | 3.7 | 0.0002 | 0.0026 | ##### | 650.40 | 737.70 | 701.34 | 645.93 | 632.67 | 471.01 | 477.11 | 409.38 | 522.91 | 577.02 | 374.81 |
| GSTZ1 | 195.43 | 0.409803 | 0.2 | 2.7 | 0.0076 | 0.0393 | ##### | 200.59 | 299.99 | 232.14 | 224.80 | 215.71 | 161.37 | 178.35 | 161.66 | 140.67 | 177.65 | 185.87 |
| DHRS7 | 550.91 | 0.404229 | 0.1 | 2.9 | 0.004 | 0.0244 | ##### | 640.53 | 729.21 | 609.47 | 720.54 | 468.57 | 457.93 | 420.31 | 447.18 | 490.80 | 427.43 | 601.54 |
| NUDC | 746.58 | 0.400649 | 0.2 | 2.6 | 0.0099 | 0.0477 | ##### | 715.75 | 1,109.38 | 786.65 | 1,088.66 | 695.63 | 665.09 | 552.09 | 636.99 | 683.45 | 681.21 | 642.39 |
| TMBIM6 | 10455 | 0.322145 | 0.1 | 3.5 | 0.0004 | 0.0048 | ##### | 11,330.40 | 13,118.24 | 12,330.51 | 12,688.92 | 10,007.13 | 8,913.23 | 9,644.45 | 8,749.81 | 9,979.60 | 9,185.65 | 9,286.57 |

### Supplementary Table S3: In vivo M1(=LPS+) and M2(=LPS-) associated genes as reported in Buscher et al Nat Comm 2017)

**Supplementary Table S3**
**Full *In vivo* M1(=LPS+) and M2(=LPS-) as reported in the original paper (Buscher et al Nat Comm 2017)**

| M1 (=LPS+) Signature | M2 (=LPS-) Signature |
|---|---|
| 2610001J05Rik | 0610009B22Rik |
| 2610002M06Rik | 1810058I24Rik |
| A630072M18Rik | 2010204K13Rik |
| AARS | 2310039H08Rik |
| ABCA1 | 2510002D24Rik |
| ABCB1 | 2610524H06Rik |
| ABCF1 | 2700097O09Rik |
| ABHD16A | 2900097C17Rik |
| ABHD2 | 9130221H12Rik |
| ABI1 | AAAS |
| ABRACL | AACS |
| ACBD3 | AAGAB |
| Acot5 | AAK1 |
| ACOT7 | AAMP |
| Acp5 | AARSD1 |
| ACSL1 | ABCA7 |
| ACSL5 | ABCB6 |
| ACTA1 | ABCB8 |
| ACTL7B | ABCD3 |
| ACTN1 | ABCD4 |
| ACTR3 | ABCF3 |
| ACVR1 | ABHD12 |
| ADAM17 | ABHD17A |
| Adam1b | ABHD4 |
| ADAMTS4 | ABHD5 |
| ADAR | ABHD8 |
| ADGRB1 | ABL1 |
| ADHFE1 | ABTB1 |
| ADM | ACAA2 |
| ADORA2A | ACAD10 |
| ADORA2B | ACAD11 |
| ADPRH | ACAD8 |
| AEBP2 | ACADM |
| AEN | ACADS |
| AFF1 | ACADVL |
| AFTPH | ACAT2 |
| AGFG1 | ACBD6 |
| AGRN | ACD |
| AGTRAP | ACLY |
| AIDA | ACO1 |
| AIM1 | ACO2 |
| AK4 | ACOT13 |
| AKAP4 | ACOX1 |
| AKNA | ACP6 |
| AKT3 | ACRBP |
| ALDH3A1 | ACSS1 |
| ALG9 | ACTR1A |
| ALPK2 | ACTR6 |
| ANKFY1 | ACTR8 |
| ANKIB1 | ADAM10 |
| ANKRD17 | ADAM15 |
| ANXA7 | ADAM8 |
| Ap2b1 | ADAMTS10 |
| AP3B1 | ADCK1 |
| AP3B2 | ADCK3 |
| APBB1IP | ADCK4 |
| API5 | ADCY7 |
| APOBEC3B | ADCY9 |
| APOBR | ADD1 |
| APPBP2 | ADD3 |
| APPL1 | ADGRE5 |
| APTX | ADH5 |
| AQP9 | ADIPOR1 |
| AQR | ADIPOR2 |
| ARCN1 | ADK |
| AREG | ADSSL1 |
| ARF4 | AES |
| ARFRP1 | Afg3l1 |
| ARG2 | AFG3L2 |
| ARHGAP35 | AGAP1 |
| ARHGEF2 | AGAP3 |
| ARHGEF3 | AGFG2 |
| ARHGEF40 | AGGF1 |
| ARID4B | AGL |
| ARID5A | AGO2 |
| ARIH1 | AHCYL1 |
| ARIH2 | AHNAK |
| ARL14EP | AHSA1 |
| ARMCX3 | Ahsa2 |
| ARPP21 | AI413582 |
| ARRDC4 | AI987944 |
| ART3 | AIFM1 |
| ASAP1 | AKAP10 |
| ASB13 | Akap17b |
| ASCC2 | AKAP8 |
| ASNS | AKAP8L |
| ATAD1 | AKIP1 |
| ATAD2B | AKR1A1 |
| ATF2 | AKR1B10 |
| ATF4 | AKR7A2 |
| ATL3 | AKT1 |
| ATP10A | AKTIP |
| ATP11A | ALAD |
| ATP11C | ALDH3B1 |
| ATP13A3 | ALDH6A1 |
| AW112010 | ALDH9A1 |
| AXIN1 | ALDOA |
| AXL | ALG1 |
| AZGP1 | ALG12 |
| AZI2 | ALG14 |
| B3GNT2 | ALG2 |
| BATF | AMACR |
| BATF2 | AMOT |
| BATF3 | AMZ2 |
| BC147527 | ANAPC1 |
| BCL10 | ANAPC13 |
| BCL2L11 | Anapc15 |
| BIRC2 | ANAPC16 |
| BIRC3 | ANAPC2 |
| BLOC1S4 | ANAPC4 |
| BLZF1 | ANAPC5 |
| BRAF | ANGEL1 |
| BRD2 | ANGEL2 |
| BRIX1 | ANGPT2 |
| BROX | ANGPTL6 |
| Bst2 | ANKMY2 |
| BTG2 | ANKRD10 |
| BZW1 | ANKRD24 |
| C11orf68 | ANKRD40 |
| C11orf96 | ANKRD46 |
| C14orf105 | ANKRD54 |
| C16orf87 | ANKS1A |
| C17orf96 | ANKS3 |
| C19orf66 | Anp32a |
| C1orf123 | Anp32b |
| C21orf91 | ANTXR2 |
| C2orf47 | ANXA9 |
| C2orf76 | AP1B1 |
| C3orf38 | AP1M1 |
| C5AR1 | AP2A2 |
| C5orf30 | AP2M1 |
| C8orf33 | AP2S1 |
| C8orf44-SGK3 | AP3D1 |
| SGK3 | AP3S1 |
| CA13 | AP5S1 |
| CAB39L | APBA3 |
| CACYBP | APBB3 |
| CALCRL | APC |
| CAMK2D | APEH |
| CAMLG | APH1A |
| CAPZA2 | APLF |
| CARHSP1 | APPL2 |
| CASK | APRT |
| CASP1 | ARAP1 |
| CASP4 | ARFGAP1 |
| CASP7 | ARFIP2 |
| CAV1 | ARG1 |
| CBWD1 | ARHGAP12 |
| Ccdc50 | ARHGAP18 |
| CCDC71L | ARHGAP24 |
| CCDC86 | ARHGAP27 |
| CCDC93 | ARHGAP9 |
| CCL17 | ARHGDIB |
| Ccl2 | ARHGEF1 |
| CCL22 | ARHGEF10L |
| CCL24 | ARHGEF18 |
| Ccl7 | ARID3A |
| Ccl9 | ARL2 |
| CCNC | ARL5A |
| CCND2 | ARL6 |
| CCNG2 | ARL6IP4 |
| CCNJ | ARL8B |
| CCNL1 | ARMC1 |
| CCR9 | ARMC10 |
| CCRL2 | ARMT1 |
| CD14 | ARPC1A |
| CD274 | ARPC5L |
| CD2AP | ARRB1 |
| CD2BP2 | ARRB2 |
| CD38 | ARRDC1 |
| CD40 | ARSA |
| CD44 | ARSG |
| CD47 | ASB3 |
| CD70 | GPR75-ASB3 |
| CD80 | ASB8 |
| CD83 | ASCC1 |
| CD86 | ASF1B |
| CDC37 | ASNSD1 |
| Cdc42 | ASPSCR1 |
| CDC73 | ASRGL1 |
| CDH16 | ASXL1 |
| CDK5R1 | ATF1 |
| CDK6 | ATF6 |
| CDKN1A | ATG101 |
| CDV3 | ATG13 |
| CDYL | ATG2A |
| CEBPD | ATG2B |
| Celf1 | ATG5 |
| CEP350 | ATG7 |
| CEP85 | ATMIN |
| CEPT1 | ATOX1 |
| CFLAR | ATP13A2 |
| CH25H | ATP1A3 |
| CHAC1 | ATP5A1 |
| CHD1 | ATP5C1 |
| CHIC2 | ATP5D |
| CHMP4B | Atp5e |
| Chtop | ATP5G1 |
| CHUK | ATP5G2 |
| CISH | ATP5G3 |
| CITED2 | ATP5H |
| CKLF | ATP5J |
| CLCF1 | ATP5J2 |
| CLCN5 | Atp5k |
| CLCN6 | ATP5L |
| CLDND1 | ATP5S |
| Clec2d | ATP5SL |
| CLEC3B | ATP6AP1 |
| CLEC5A | ATP6V0A1 |
| CLIC4 | ATP6V0A2 |
| CLINT1 | ATP6V0D1 |
| CLIP4 | ATP6V0E1 |
| CLMP | ATP6V1A |
| CLN5 | ATP6V1B2 |
| CLOCK | ATP6V1C1 |
| CLP1 | ATP6V1D |
| CMPK2 | ATPAF2 |
| CMTM6 | ATPIF1 |
| CMTR1 | ATRAID |
| CNEP1R1 | ATXN10 |
| CNIH4 | ATXN2 |
| CNOT2 | AU020206 |
| CNOT4 | AUH |
| CNOT7 | AUP1 |
| COG2 | B3GALNT2 |
| COX15 | B3GALT4 |
| CP | B3GAT3 |
| CPD | B3GNT8 |
| CPN2 | B3GNTL1 |
| CPNE2 | B4GALNT1 |
| CPSF2 | B4GALT1 |
| CR1L | B4GALT6 |
| CREB1 | B4GAT1 |
| CRK | BABAM1 |
| CRKL | BACE1 |
| CRLF3 | BACH1 |
| CS | BAD |
| CSDE1 | BAG1 |
| CSF1 | BAG3 |
| CSF2 | BAG4 |
| CSNK1A1 | BAIAP2 |
| CST7 | BANP |
| Ctsq | BAP1 |
| CUEDC1 | BASP1 |
| CUL2 | BAX |
| CXCL10 | BAZ1B |
| Cxcl11 | BAZ2A |
| Cxcl15 | BBS2 |
| CXCL16 | BBS9 |
| CXCL3 | BC029722 |
| Cxcl9 | BCAP31 |
| CYBB | BCAS2 |
| CYCS | BCAS3 |
| CYSLTR2 | BCDIN3D |
| DAB2 | BCKDHA |
| DAPK3 | BCKDK |
| DAXX | BCL6 |
| DAZ2 | BCS1L |
| DBNL | BET1 |
| DBR1 | BIN3 |
| DCAF13 | BLOC1S1 |
| DCBLD2 | BLOC1S5 |
| DCK | Bmyc |
| DCP1A | BNIP2 |
| DCTN4 | BNIP3 |
| DCUN1D1 | BNIP3L |
| DCUN1D5 | BOLA1 |
| DDA1 | BORCS5 |
| DDHD1 | BORCS7 |
| DDIT4 | BPGM |
| DDX10 | BPTF |
| DDX17 | BRAP |
| DDX21 | BRAT1 |
| DDX24 | BRD3 |
| DDX27 | BRK1 |
| DDX3X | BSCL2 |
| DDX60 | BSG |
| DENND5A | BST1 |
| DENR | BTBD1 |
| DGCR14 | BTBD2 |
| DHX15 | BTRC |
| DHX58 | C10orf54 |
| DIMT1 | C11orf1 |
| DIS3 | C11orf31 |
| DLD | C11orf54 |
| DLGAP4 | C11orf74 |
| DLL1 | C11orf98 |
| DLX3 | C12orf10 |
| DNAJA1 | C12orf43 |
| DNAJA2 | C12orf57 |
| DNAJB5 | C14orf1 |
| DNAJB6 | C14orf166 |
| DNAJC1 | C15orf39 |
| DNAJC13 | C15orf40 |
| DPF2 | C16orf70 |
| DR1 | C16orf91 |
| DTWD1 | C17orf62 |
| DUSP11 | C18orf8 |
| DUSP16 | C19orf44 |
| DUSP2 | C19orf47 |
| DUSP8 | C19orf52 |
| DYNC1LI1 | C19orf53 |
| E2F3 | C19orf60 |
| E2F5 | C1D |
| E2F8 | C1orf159 |
| EBI3 | C1orf27 |
| EBNA1BP2 | C1QBP |
| EDN1 | C20orf27 |
| EDNRB | C21orf2 |
| EEF1E1 | C2CD2L |
| EFR3A | C2CD3 |
| EGR1 | C2orf42 |
| EHD1 | C2orf49 |
| Eif1 | C3AR1 |
| EIF1AX | C3orf58 |
| EIF2AK2 | C3orf70 |
| EIF2S2 | C4orf27 |
| EIF2S3 | C5orf22 |
| EIF3C | C6orf136 |
| EIF4E | C6orf203 |
| EIF4E2 | C6orf89 |
| EIF4E3 | C7orf25 |
| EIF6 | C7orf55 |
| ELAVL1 | C8orf37 |
| ELL2 | C8orf82 |
| ELP5 | C9orf16 |
| ENDOD1 | C9orf3 |
| ENG | C9orf40 |
| EPS15L1 | C9orf69 |
| EPSTI1 | C9orf91 |
| ERAL1 | CAB39 |
| ERAP1 | CACFD1 |
| ERBB2IP | Calm1 |
| ERCC8 | CAMK1 |
| ERGIC2 | CAMK1D |
| ERO1A | CAMK2G |
| ERRFI1 | CANT1 |
| ESF1 | CAPN7 |
| ESRRB | CAPNS1 |
| ETF1 | CARKD |
| ETV6 | CASD1 |
| EVI2A | CASP2 |
| EXOSC10 | CASP6 |
| EXT1 | CASP9 |
| EZR | CBFA2T2 |
| F10 | CBR1 |
| F3 | CBR3 |
| FABP3 | CBR4 |
| FAM107B | CBX2 |
| FAM129A | CBX3 |
| FAM133B | CBX5 |
| FAM188A | CBX8 |
| FAM208B | CBY1 |
| FAM96B | CC2D1A |
| FAR1 | CCAR2 |
| FBXL3 | CCDC107 |
| FBXO11 | CCDC115 |
| FBXO42 | CCDC12 |
| FBXW11 | CCDC137 |
| FCER1G | CCDC22 |
| FCGR2B | CCDC28A |
| FEM1B | CCDC34 |
| FGL1 | CCDC84 |
| FITM1 | CCDC91 |
| FKBP5 | CCDC97 |
| FLNB | CCNF |
| FMR1 | CCNG1 |
| FNBP1 | CCNH |
| FNBP4 | CCPG1 |
| FNDC3A | CCR1 |
| FRMD6 | CCS |
| FUBP1 | CCZ1 |
| Fus | CCZ1B |
| FYN | CD151 |
| FZD1 | CD200R1 |
| FZD5 | CD28 |
| GABPA | CD300A |
| GABPB1 | CD300LF |
| GADD45B | CD320 |
| Gar1 | CD36 |
| GARS | CD84 |
| GAS7 | CD99L2 |
| GATA1 | CDADC1 |
| GATAD2A | CDC123 |
| GBP2 | CDC23 |
| GBP7 | CDC25A |
| GCA | CDC25B |
| GCC1 | CDC37L1 |
| GCH1 | CDC42EP3 |
| GDF15 | CDC42SE1 |
| GEM | CDC5L |
| GFI1 | CDCA5 |
| GFPT1 | CDCA7L |
| GFPT2 | CDIP1 |
| GFRA2 | CDIPT |
| GGCT | CDK11A |
| GHITM | CDK16 |
| GIT2 | CDK18 |
| Gk | CDK2 |
| GLCE | CDK20 |
| Gm15807 | CDK2AP1 |
| Hmgn5 | CDK2AP2 |
| Gm9706 | CDK5 |
| GMEB1 | CDK5RAP2 |
| GMFB | CDK5RAP3 |
| GMPPB | CDK9 |
| GNA13 | CDKN2AIPNL |
| GNAI3 | CDKN2C |
| GNAQ | CDPF1 |
| GNB4 | CDT1 |
| GNPNAT1 | CEBPA |
| GOLPH3 | Cebpzos |
| GOSR1 | CELA1 |
| GOSR2 | CELF2 |
| GPBP1 | CELSR3 |
| GPD2 | CENPA |
| GPR84 | CENPO |
| GPR85 | CENPV |
| GPR87 | CEP131 |
| GPRC5C | CEP152 |
| GRAMD1A | CEP19 |
| GRHL1 | CEP57L1 |
| GRIN1 | CEP89 |
| GSPT1 | CERK |
| GTF2B | CERS2 |
| GTF2F1 | CERS5 |
| GTF2F2 | CETN2 |
| GTPBP2 | CFAP20 |
| GTPBP4 | CFDP1 |
| GUCA2B | CFL1 |
| GYPC | CFL2 |
| H3F3A | CGNL1 |
| H3F3B | CHCHD10 |
| HACD2 | CHCHD5 |
| Hamp | CHCHD7 |
| Hamp2 | CHD1L |
| HAT1 | CHD6 |
| HBEGF | CHD8 |
| HCK | CHEK2 |
| HDC | CHERP |
| HEATR6 | CHFR |
| HELZ2 | CHID1 |
| HERPUD1 | CHKA |
| HIF1A | CHMP1A |
| HIPK1 | CHMP3 |
| HIPK2 | CHMP6 |
| HIVEP1 | CHMP7 |
| HIVEP2 | CHPF2 |
| HIVEP3 | CHST12 |
| HK2 | CHST14 |
| HLA-G | CIB1 |
| HMGCR | CIC |
| HNRNPAB | CIPC |
| HOMER1 | CIR1 |
| HSP90AA1 | CISD1 |
| HSPA5 | CIZ1 |
| HSPA9 | CKAP4 |
| HTR1B | CLCC1 |
| HUS1 | CLCN3 |
| IARS | CLCN4 |
| IBTK | CLIP1 |
| ICAM1 | CLK2 |
| ICAM4 | CLK4 |
| ICOSLG | CLN6 |
| LOC102723996 | CLN8 |
| IDNK | CLNS1A |
| IER2 | CLPB |
| IER3 | CLPP |
| IER5 | CLPTM1L |
| IFI16 | CLPX |
| IFI35 | CLTA |
| Ifi47 | CLTB |
| IFIH1 | CLTC |
| IFIT2 | CLUAP1 |
| Ifitm1 | CLUH |
| IFITM3 | CMAS |
| IFNAR1 | CMC2 |
| IFNAR2 | CMC4 |
| IFNB1 | CMIP |
| IGF1R | CNNM2 |
| IGF2BP1 | CNNM3 |
| IGSF8 | CNOT6L |
| IGSF9 | CNOT8 |
| Igtp | CNPY3 |
| Iigp1 | CNR2 |
| IKBKB | COA3 |
| IKZF1 | Coa6 |
| IL10RA | COASY |
| IL12A | COG4 |
| IL12B | COG6 |
| IL13RA1 | COMMD5 |
| IL15 | Commd6 |
| IL15RA | COMMD9 |
| IL17RA | COMT |
| IL17RD | COPB2 |
| IL18 | COPG1 |
| IL1RAP | COPG2 |
| IL1RN | COPRS |
| IL23A | COPS3 |
| IL2RG | COPS7A |
| IL4R | COPS7B |
| INHBA | COPZ1 |
| INO80 | COQ10A |
| INPP5B | COQ7 |
| INTS12 | COQ9 |
| IRF1 | CORO1B |
| IRF2 | CORO1C |
| IRF5 | COTL1 |
| IRF7 | COX14 |
| IRF9 | COX4I1 |
| IRG1 | COX6A1 |
| IRGM | COX6B1 |
| Irgm1 | Cox6c |
| ISG20 | COX7A2 |
| IST1 | CPEB2 |
| ITGA4 | CPOX |
| ITGA5 | CPPED1 |
| ITGAL | CPSF1 |
| ITGAV | CPSF3 |
| ITGB1 | CPT1A |
| JAG1 | CPT2 |
| JAK1 | CRACR2B |
| JAK2 | CRAT |

### Supplementary Table S3

| | |
|---|---|
| JARID2 | CREBRF |
| JDP2 | CREBZF |
| JUNB | CREG1 |
| KALRN | CRELD1 |
| KATNA1 | CRIP1 |
| KATNBL1 | CROT |
| KBTBD2 | CRTAP |
| KCND3 | CSAD |
| KCTD10 | CSGALNACT2 |
| KCTD5 | CSK |
| KDM7A | CSNK1D |
| KEAP1 | CSNK1E |
| KHDRBS1 | CSNK1G2 |
| KIAA1033 | CSTB |
| KIAA1551 | CSTF3 |
| KIF1B | CTAGE5 |
| KIF3B | CTBP1 |
| KIF5B | CTC1 |
| KLF3 | CTCF |
| KLF5 | CTDP1 |
| KLF6 | CTDSP1 |
| KLF7 | CTDSP2 |
| KLHL25 | CTDSPL |
| Klrk1 | CTNNB1 |
| KMO | CTNNBIP1 |
| KMT5B | CTNNBL1 |
| KPNA3 | CTNS |
| KPNA4 | CTPS2 |
| KPNA6 | CTSA |
| KREMEN1 | CTSB |
| KRI1 | CTSD |
| KRIT1 | CUEDC2 |
| Ktn1 | CUL4B |

| | |
|---|---|
| LAD1 | CUL7 |
| LANCL2 | CUTA |
| LARP1 | Cux1 |
| LBP | CWC27 |
| LCK | CXCR4 |
| LCN2 | CXorf38 |
| LCP1 | CXorf40A |
| LCP2 | CXorf40B |
| LENG1 | CXXC1 |
| LGALS3BP | CXXC5 |
| LGALS8 | CYB561D2 |
| LGALS9B | CYB5A |
| LIF | Cyb5r3 |
| LIG3 | CYHR1 |
| LIMS1 | Cyp4f16 |
| LIN7C | Cyp4f37 |
| LMNB2 | DAD1 |
| LMO4 | DAG1 |
| LPAR1 | DALRD3 |
| LRRC8C | DAP |
| LRRK2 | DAP3 |
| LSG1 | DAZAP2 |
| LSM8 | DBF4 |
| LTA | DBI |
| LTV1 | DBNDD2 |
| LYN | DBP |
| LYRM1 | DCAF11 |
| LZTFL1 | DCAF4 |
| MAD2L1 | DCAF8 |
| MAFF | DCLRE1A |
| MAFK | DCPS |
| MAGOHB | DCTN2 |
| MAK16 | DCTN6 |

| | |
|---|---|
| MAN2A1 | DCTPP1 |
| MAP2K1 | DDB1 |
| MAP2K4 | DDHD2 |
| MAP3K5 | DDRGK1 |
| Map3k7 | DDX50 |
| MAP3K8 | DEAF1 |
| MAP4 | DEDD2 |
| MAPK14 | DEF8 |
| MAPK6 | DENND4B |
| MAPK8 | DENND4C |
| MAPRE1 | DENND6A |
| Mapre2 | DEPDC7 |
| Marcks | DERA |
| MARCKSL1 | DERL1 |
| MARK1 | DESI1 |
| MARK2 | DESI2 |
| MAT2A | DEXI |
| MBOAT1 | DFNA5 |
| MCL1 | DGAT1 |
| MDFIC | Dgcr6 |
| MDM2 | DGKA |
| MDM4 | DGKZ |
| MECP2 | DGUOK |
| MED1 | DHCR7 |
| MED11 | DHFR |
| MED14 | DHODH |
| MED19 | DHRS1 |
| MED21 | DHRS3 |
| MED27 | DHRS4 |
| MED6 | DHRS7 |
| MED7 | DHX16 |
| MEFV | DHX30 |
| MET | DHX32 |

| | |
|---|---|
| METTL6 | DHX57 |
| MFSD1 | DICER1 |
| MFSD14B | DIO2 |
| MFSD4B | DLG3 |
| MITD1 | DNAJB1 |
| MITF | DNAJB9 |
| MKL2 | DNAJC11 |
| MLLT6 | DNAJC12 |
| MMADHC | DNAJC14 |
| MMP10 | DNAJC28 |
| MMP13 | DNAJC3 |
| MMP14 | DNAJC30 |
| MOB3B | DNAJC5 |
| MORC3 | DNAJC7 |
| MORF4L2 | DNAJC9 |
| MOV10 | DNAL4 |
| MPP5 | DNLZ |
| MRPL21 | DNMBP |
| MRPL32 | DNMT1 |
| MRPS22 | DNMT3A |
| Ms4a4b | DNPEP |
| MS4A6A | DNTTIP1 |
| Ms4a6b | DOK1 |
| Ms4a6c | DOK2 |
| MSANTD3 | DOK3 |
| MSN | DOLK |
| MTDH | DOLPP1 |
| MTF2 | DOPEY2 |
| MTHFD2 | DPAGT1 |
| MTHFR | DPCD |
| MTMR14 | DPH7 |
| MTMR7 | DPM1 |
| MTPN | DPP3 |

| | |
|---|---|
| Mucl1 | DPY30 |
| Mucl2 | DRAP1 |
| MUL1 | DSCR3 |
| MVP | DSTYK |
| Mx1/Mx2 | DTNBP1 |
| MYADM | DTX3 |
| MYBBP1A | DUS1L |
| MYC | DUSP13 |
| MYCL | DUSP19 |
| MYD88 | DUSP22 |
| MYL12B | DUSP3 |
| MYNN | DVL2 |
| MYO10 | DYM |
| MYO1C | DYNLL1 |
| MYO1G | DYNLL2 |
| N4BP1 | E2F1 |
| NAA15 | E2F6 |
| NACC1 | E4F1 |
| NAE1 | EBAG9 |
| NAMPT | EBPL |
| NARS | ECD |
| NAT1 | ECH1 |
| NCF1 | ECHDC1 |
| NCK1 | ECHS1 |
| NCOA3 | ECSIT |
| NCOA5 | EDEM1 |
| NDEL1 | EDEM2 |
| NEDD9 | EDEM3 |
| NEURL1 | EED |
| NFAT5 | EEF2 |
| NFATC1 | EEF2KMT |
| NFATC2 | EEFSEC |
| NFIL3 | EEPD1 |

| | |
|---|---|
| NFKB1 | EFCAB7 |
| NFKB2 | EFTUD2 |
| NFKBIA | EGLN1 |
| NFKBIB | EGLN2 |
| NFKBIE | EHBP1 |
| NFKBIZ | Ehbp1l1 |
| NFXL1 | EHMT2 |
| NFYB | EI24 |
| NIF3L1 | EID1 |
| NMD3 | EIF1B |
| NMI | EIF2AK1 |
| NOC3L | EIF2AK3 |
| NOCT | EIF2B4 |
| NOL11 | EIF2D |
| NOP58 | EIF3I |
| NOS2 | EIF3K |
| NOTCH1 | EIF4B |
| NOTCH2 | EIF4EBP2 |
| NR3C1 | ELAC2 |
| NRAS | ELF2 |
| NRF1 | ELK3 |
| NRIP3 | ELL |
| NT5C3A | ELMO2 |
| NUB1 | ELMOD3 |
| NUBP1 | ELOVL1 |
| NUDT9 | ELOVL5 |
| NUFIP1 | ELP2 |
| NUP54 | ELP3 |
| NUP62 | ELP4 |
| NUP98 | EMC10 |
| NUPR1 | EMC2 |
| OAF | EMC3 |
| Oas1b | EMC8 |

| | |
|---|---|
| OAS2 | EMC9 |
| OAS3 | EMILIN1 |
| OASL | EML3 |
| Oasl2 | EMSY |
| OCSTAMP | ENDOG |
| OGFR | ENO3 |
| OLAH | ENOPH1 |
| OPA3 | ENTHD2 |
| OSGIN2 | ENTPD5 |
| OTUD4 | ENTPD6 |
| Otud5 | ENTPD7 |
| P2RY14 | EPB41 |
| P4HA1 | EPB41L1 |
| PAFAH1B2 | EPN1 |
| PANK3 | EPS8 |
| PAPD4 | ERCC3 |
| PAPSS2 | ERCC4 |
| PARP12 | ERCC5 |
| PARP8 | ERCC6L2 |
| PARP9 | ERF |
| Pcmt1 | ERGIC1 |
| PCNX | ERGIC3 |
| PDCD10 | ERI3 |
| PDE12 | ERLEC1 |
| PDE4B | ERLIN1 |
| PDGFRB | ERMP1 |
| PDLIM5 | ERP29 |
| PDPN | ESCO1 |
| PDSS1 | ETAA1 |
| PDZD11 | ETFB |
| PELI1 | ETFBKMT |
| PENK | ETFDH |
| PES1 | ETHE1 |

| | |
|---|---|
| PFKP | EXD2 |
| Pgap2 | EXOC8 |
| PHLDB1 | EXOSC4 |
| PHLPP1 | EXOSC5 |
| PHTF2 | EXOSC7 |
| PHYKPL | EXT2 |
| PICALM | EXTL3 |
| PIK3AP1 | F8A1 |
| PIK3R1 | FAAP20 |
| PILRA | FADS1 |
| PIM3 | FAF1 |
| PINX1 | FAHD1 |
| PIP5K1A | FAIM |
| PITPNB | FAM102A |
| PKMYT1 | FAM104A |
| PKP4 | FAM105A |
| PLA1A | FAM111A |
| PLA2G16 | FAM117A |
| PLAGL1 | FAM117B |
| PLAGL2 | FAM118B |
| PLAUR | FAM120A |
| PLCD1 | FAM120B |
| PLCL2 | FAM126B |
| PLCZ1 | FAM129B |
| PLEC | FAM134A |
| PLEK | FAM134B |
| PLEKHA3 | FAM160A2 |
| PLEKHF1 | FAM162A |
| PLET1 | FAM173A |
| PLK2 | FAM189B |
| PLSCR1 | Fam193b |
| PMEPA1 | FAM195A |
| PML | FAM198B |

| | |
|---|---|
| PMPCA | FAM210B |
| PMPCB | FAM214B |
| PNP | FAM216A |
| POFUT2 | FAM21A |
| POLA2 | FAM21C |
| POLDIP3 | FAM234B |
| POLR2C | FAM32A |
| POLR2G | FAM3C |
| POLR3C | FAM50A |
| POLR3D | FAM53A |
| POP4 | FAM58A |
| POP7 | FAM63A |
| Pou3f1 | FAM65A |
| PPA1 | FAM73B |
| PPM1A | FAM76A |
| PPM1B | FAM76B |
| PPP1CB | FAM89B |
| PPP1R11 | FAM96A |
| PPP1R15A | FAM98C |
| PPP1R15B | FANCE |
| PPP1R2 | FANCF |
| PPP2CB | FANCG |
| PPP2R2A | FASTK |
| PPP2R2D | FASTKD1 |
| PPP3CC | FBXL14 |
| PPP4C | FBXL20 |
| PPP4R1 | FBXL6 |
| PRDM1 | FBXL8 |
| PRDM4 | FBXO18 |
| PRELID3B | FBXO21 |
| PREP | FBXO22 |
| PRKCD | FBXO25 |
| PRKCQ | FBXO28 |

| | |
|---|---|
| PRKRIP1 | FBXO32 |
| PRKX | FBXO33 |
| PRMT6 | FBXO4 |
| PRPF38A | FBXO8 |
| PRPF40A | FBXO9 |
| PSMA2 | FBXW2 |
| PSMB10 | FBXW4 |
| PSMB6 | FBXW7 |
| PSMB8 | FBXW9 |
| PSMB9 | FCGR2A |
| PSMD7 | FCHO1 |
| PSME1 | FDXR |
| PSME2 | FECH |
| PSPC1 | FEN1 |
| PSTPIP2 | FERMT3 |
| PTBP1 | FES |
| PTBP3 | FGD4 |
| PTGES | FGFR1 |
| PTGES3 | FH |
| PTGS2 | FHDC1 |
| PTPN11 | FIGNL1 |
| PTPN12 | FIS1 |
| PTPN2 | FKBP1A |
| PTPRC | FKBP1B |
| PTPRJ | FKBP2 |
| PTX3 | FKBP3 |
| PUM1 | FKBP4 |
| PUM2 | FKBP8 |
| PUM3 | FKBPL |
| PURB | FLAD1 |
| Pvr | FLCN |
| PWP1 | FLI1 |
| PXN | FLOT2 |

| | |
|---|---|
| RAB11A | FLYWCH1 |
| RAB12 | FMNL3 |
| RAB22A | FNTA |
| RAB32 | FOPNL |
| RAB38 | FOS |
| RAB3IP | FOXJ2 |
| RAB5A | FOXK2 |
| RABGEF1 | FPGS |
| RALB | FRMD8 |
| RAN | FTO |
| RANBP1 | FTSJ1 |
| RAP1B | FTSJ2 |
| RAP2C | FUCA2 |
| RARS | FUNDC1 |
| RASA2 | FUNDC2 |
| RASGRP1 | FXN |
| RBBP8 | FXYD5 |
| RBM4 | FYCO1 |
| RBM43 | G6PC3 |
| RBM7 | GAA |
| RBMS1 | GAB1 |
| RBMX | GABARAP |
| RBPJ | GABARAPL1 |
| RBPMS | GADD45G |
| RCAN1 | GALK1 |
| RCL1 | GALK2 |
| RCN1 | GALNT1 |
| RCSD1 | GALNT2 |
| RDH11 | GALNT4 |
| RDH12 | GALT |
| RDX | GART |
| REL | GATAD1 |
| RELA | GATC |

| | |
|---|---|
| REPS1 | GBA |
| RFFL | GBA2 |
| RFWD2 | GBAS |
| Rfx5 | GBF1 |
| RGPD4 | GCLC |
| RGS1 | GDI2 |
| RGS3 | GDPD3 |
| RHBDD1 | GEMIN6 |
| RHBDF1 | GEMIN8 |
| RHOT1 | GFOD2 |
| RIOK2 | GGA2 |
| RIPK2 | Ggta1 |
| RLIM | GID4 |
| RMDN3 | GIGYF1 |
| RND3 | GINS1 |
| RNF11 | GINS4 |
| RNF114 | GIT1 |
| RNF135 | GKAP1 |
| RNF14 | GLA |
| RNF141 | GLG1 |
| RNF145 | GLMN |
| RNF19A | GLMP |
| RNF19B | GLOD4 |
| RNF2 | GLRX2 |
| RNF214 | GLRX3 |
| RNF225 | GLTP |
| RNF4 | GLUL |
| RNGTT | Gm10374 |
| RNPC3 | Josd1 |
| ROCK1 | Gm10651 |
| ROCK2 | Gm21596 |
| RPE | Hmgb1 |
| RPF2 | GM2A |

| | |
|---|---|
| RPL7L1 | Gm561 |
| RPS6KA2 | Gm8587 |
| RPS6KA4 | Snrpa |
| RPS6KC1 | GMCL1 |
| RRN3 | GMIP |
| RRP15 | GMNN |
| RRP7A | GMPPA |
| RRS1 | GMPR |
| RS1 | GNA11 |
| RSAD2 | GNAI2 |
| RSL1D1 | GNB1L |
| RSL24D1 | GNB5 |
| RSPRY1 | GNE |
| Rsrc2 | GNG2 |
| RTCB | GNPAT |
| RTP4 | GNPDA1 |
| RWDD4 | GNPDA2 |
| SAMD11 | GNS |
| SAMHD1 | GOLGA2 |
| SAMSN1 | GOLGA4 |
| SAP30 | GOLGA5 |
| SAR1A | GOLM1 |
| SASS6 | GOLPH3L |
| SAT1 | GON4L |
| SAV1 | GORASP1 |
| SBDS | GOT2 |
| SBNO1 | GPAA1 |
| SDAD1 | GPATCH8 |
| SDC4 | GPCPD1 |
| SDCCAG3 | GPI |
| SEC22B | GPN3 |
| SEC23B | GPNMB |
| SEC24B | GPR146 |

| | |
|---|---|
| SEH1L | GPR180 |
| SEL1L | GPR35 |
| SEMA4A | GPR65 |
| SENP1 | GPRC5B |
| | GPS1 |
| SERBP1 | GPSM3 |
| Serpina3g | GPT |
| SERPINB2 | GPX1 |
| SERPINE1 | GPX4 |
| SERTAD1 | GRAMD4 |
| SERTAD2 | GRB2 |
| SESN2 | GRHPR |
| Sf1 | GRK6 |
| SF3A3 | GRN |
| SFPQ | GRSF1 |
| SGMS1 | GSG2 |
| SH3BGRL | GSK3A |
| SH3BP5 | GSN |
| SHMT2 | GSTM4 |
| SIAH2 | GSTM5 |
| SIX1 | GSTO1 |
| SKIL | GTF2E2 |
| SKIV2L2 | GTF2H2 |
| SLAMF9 | GTF2H4 |
| SLC11A2 | GTF2I |
| SLC12A4 | GTF2IRD1 |
| SLC15A3 | Gtf2ird2 |
| SLC1A4 | GTF3A |
| SLC23A2 | GTF3C1 |
| SLC25A17 | GTF3C2 |
| SLC25A22 | GTF3C4 |
| SLC25A25 | GTPBP1 |
| SLC25A37 | GUK1 |

| | |
|---|---|
| SLC27A2 | GUSB |
| SLC30A6 | GYG1 |
| SLC30A7 | H1F0 |
| SLC31A1 | H2AFJ |
| SLC31A2 | H2AFV |
| SLC35A3 | H2AFZ |
| SLC38A1 | H6PD |
| SLC39A14 | HABP4 |
| SLC3A2 | HACD3 |
| SLC4A7 | HACD4 |
| SLC7A11 | HACE1 |
| SLC7A2 | HACL1 |
| SLC7A5 | HADH |
| SLC7A7 | HADHB |
| SLCO3A1 | HAGH |
| Slfn1 | HARS2 |
| SLFN12L | HAUS1 |
| SLFN13 | HAUS3 |
| Slfn2 | HAUS4 |
| SLU7 | HAVCR2 |
| SLURP1 | HBP1 |
| SMAD2 | HCFC1 |
| SMG8 | HCFC1R1 |
| SMIM7 | HDAC3 |
| SMNDC1 | HDAC5 |
| SMU1 | HDAC6 |
| SMYD5 | HDAC7 |
| SNAPC3 | HDDC2 |
| SNW1 | HDGF |
| SNX10 | HDLBP |
| SNX20 | HEBP1 |
| SNX9 | HECTD3 |
| SOCS1 | HELB |

| | |
|---|---|
| SOCS2 | HERC1 |
| SOCS3 | HERPUD2 |
| SOCS4 | HES6 |
| SOCS5 | HEXA |
| SOCS7 | HEXB |
| SOD2 | HEXDC |
| SOWAHC | HEXIM1 |
| SP1 | HGH1 |
| Sp100 | HHEX |
| SPATA5 | HIBADH |
| SPHK1 | HIBCH |
| SPI1 | HIGD2A |
| SPINT2 | HINT1 |
| SPPL2A | HINT2 |
| SPRED1 | HIP1 |
| SPRY1 | HIST1H3G |
| SPRYD7 | HLTF |
| SPTAN1 | HM13 |
| SPTBN2 | HMBS |
| SPTLC2 | HMG20A |
| SQSTM1 | HMG20B |
| SRC | HMGCL |
| SREBF2 | HMGN1 |
| SRFBP1 | HMHA1 |
| SRGAP2 | HMOX2 |
| Srgn | HNRNPL |
| SRPRA | HP1BP3 |
| SRSF2 | HPS4 |
| SRSF3 | HRAS |
| Srsf5 | HS2ST1 |
| SRSF6 | HS6ST1 |
| SRSF7 | HSCB |
| SSFA2 | HSD17B10 |

### Supplementary Table S3

| | |
|---|---|
| ST3GAL1 | HSD17B4 |
| ST3GAL3 | HSD3B7 |
| ST8SIA1 | HSP90B1 |
| ST8SIA4 | HSPH1 |
| STAM2 | HTATIP2 |
| STARD3 | HTATSF1 |
| STARD5 | HTRA2 |
| STAT1 | HYAL1 |
| STAT2 | HYAL2 |
| STAT5A | IARS2 |
| STAT5B | ICMT |
| STAU1 | ICT1 |
| STIP1 | IDE |
| STK40 | IDH1 |
| STX3 | IDH2 |
| STX6 | IDH3B |
| STXBP3 | IDH3G |
| STXBP6 | IER3IP1 |
| SUCO | IFFO1 |
| SUFU | IFI30 |
| SUGT1 | IFT140 |
| SUMO1 | IFT20 |
| SUPV3L1 | IFT27 |
| SURF4 | IFT46 |
| SURF6 | IFT74 |
| SUSD6 | IFT80 |
| SUZ12 | IFT88 |
| SWAP70 | IGBP1 |
| SYF2 | IK |
| SYK | IKBIP |
| SYNCRIP | IKBKG |
| SYNE2 | IL16 |
| TAB2 | IL6R |

| | |
|---|---|
| TAF7 | ILK |
| TAGLN2 | IMP3 |
| TANK | IMPA1 |
| TAOK1 | IMPACT |
| TAP1 | IMPDH1 |
| TAP2 | Inafm2 |
| TAPBP | INCENP |
| TAPBPL | ING2 |
| TARS | ING4 |
| TBATA | INO80B |
| TBK1 | INPP5A |
| TBL1X | INPP5D |
| TBRG1 | INPP5E |
| TBRG4 | INSIG2 |
| TCEA1 | INTS3 |
| TCERG1 | INTS4 |
| TCF15 | INTS5 |
| TCF4 | INTS6L |
| TCOF1 | INVS |
| Tcstv1 | IP6K1 |
| TDRD7 | IPO11 |
| TEX12 | IPO8 |
| TEX15 | IPP |
| TFAM | IQSEC1 |
| TFEC | IRAK1 |
| TGIF1 | IRAK1BP1 |
| TGM2 | IRAK4 |
| TGS1 | IRF2BP1 |
| THBS1 | ISCA1 |
| TIMELESS | ISCU |
| TIMM10 | ITGA6 |
| TIMM17A | ITGB2 |
| TIMM23 | ITGB5 |

| | |
|---|---|
| TIPARP | ITM2C |
| TIRAP | ITPR2 |
| TJAP1 | ITPR3 |
| TLK2 | IVD |
| TLR2 | IVNS1ABP |
| TLR3 | JADE1 |
| TLR7 | JKAMP |
| TLR8 | JMJD1C |
| TM2D1 | JMY |
| TMCO3 | JTB |
| TMEM11 | JUN |
| TMEM128 | KANK3 |
| TMEM135 | KAT2A |
| TMEM168 | KAT7 |
| TMEM183A | KATNB1 |
| TMEM2 | KCNAB2 |
| TMEM209 | KCNK13 |
| TMEM229B | KCTD17 |
| TMEM243 | KCTD2 |
| TMEM39A | KDELC1 |
| TMOD3 | KDELR1 |
| TMPRSS2 | KDM2B |
| Tmsb4x | KDM3A |
| TNFAIP2 | KDM4B |
| TNFRSF1B | KIAA0100 |
| TNFRSF8 | KIAA0141 |
| TNFSF10 | KIAA0319L |
| TNIP1 | KIAA0922 |
| TNMD | KIAA0930 |
| TNPO3 | KIAA1522 |
| TOE1 | KIF16B |
| TOMM70A | KIF1C |
| TOP1 | KIF3A |

| | |
|---|---|
| TOR1AIP1 | KIFAP3 |
| TOR1AIP2 | KIFC3 |
| TOR3A | KLC1 |
| TOX4 | KLC4 |
| TPBG | Klf16 |
| TPM3 | KLHDC2 |
| TPST1 | KLHDC3 |
| TRA2A | KLHL21 |
| TRA2B | KLHL22 |
| TRAF1 | KLHL26 |
| TRAF2 | KLHL42 |
| TRAF5 | KLHL5 |
| TRAF6 | KLHL9 |
| TRAFD1 | KMT5C |
| TREX1 | KNOP1 |
| TRIB1 | KPTN |
| TRIB3 | KRAS |
| TRIM13 | KRTCAP2 |
| TRIM21 | LACE1 |
| TRIM25 | LACTB |
| TRIM26 | Lage3 |
| Trim30a | LAMP1 |
| Trim30d | LAMP2 |
| TRIM35 | LAMTOR1 |
| TRIM6 | LAMTOR2 |
| TRIM34 | LAMTOR4 |
| TRIOBP | LANCL1 |
| TRIP10 | LAPTM4B |
| TRMT5 | LAPTM5 |
| TRNT1 | LARGE |
| TSLP | LAS1L |
| TSPAN33 | LASP1 |
| TSR1 | LAT2 |

| | |
|---|---|
| TSR2 | LCMT1 |
| TSSC1 | LDB1 |
| TTC1 | LDHB |
| TTC39B | LDLRAP1 |
| TTC39C | LENEP |
| TTL | LEPROT |
| Tug1 | LETMD1 |
| TUT1 | LFNG |
| TXNDC9 | LGALS1 |
| TXNL1 | LIAS |
| TXNRD1 | LIG1 |
| TYK2 | LIME1 |
| TYW5 | LIMK1 |
| U2SURP | LIMK2 |
| UBA3 | LIN37 |
| UBA7 | LIPA |
| UBE2D2 | LIPT1 |
| UBE2D3 | LIPT2 |
| UBE2E1 | LMAN2 |
| UBE2F | LMNA |
| UBE2H | LMO2 |
| UBE2N | LNX2 |
| UBE2S | LOC102637129 |
| UBE2V2 | S100a11 |
| UBE2W | LOC102724828 |
| UBE2Z | MRPL23 |
| UBE4A | LOC284023 |
| UBR2 | LPGAT1 |
| UBXN2A | LPIN1 |
| UGCG | LPXN |
| UPP1 | LRP1 |
| USB1 | LRP6 |
| USP12 | LRPAP1 |

| | |
|---|---|
| USP15 | LRRC20 |
| USP16 | LRRC40 |
| USP18 | LRRC61 |
| USP25 | LRRC8A |
| USP3 | LRRFIP1 |
| USP47 | LRWD1 |
| USP9X | LSM14B |
| UTP11L | LSM3 |
| UTP6 | LSS |
| VAMP2 | LTA4H |
| VASP | LTBR |
| VAV3 | LTC4S |
| VBP1 | LUC7L2 |
| VCAM1 | LXN |
| VCAN | LYL1 |
| VCL | LYRM5 |
| Vnn3 | LYST |
| VOPP1 | LZTR1 |
| VPRBP | MAD2L1BP |
| VPS37C | MAF1 |
| VPS54 | MAN1A2 |
| VTA1 | MAN2C1 |
| WAC | MANBA |
| WARS | MAP1LC3A |
| WDR33 | MAP1LC3B |
| WDR36 | MAP2K2 |
| WDR43 | MAP2K5 |
| WDR48 | MAP2K7 |
| WDR5 | MAP3K12 |
| WDR74 | MAP3K3 |
| WNT8B | MAP3K4 |
| WRN | MAP4K1 |
| XIAP | MAP7D1 |

| | |
|---|---|
| XPO4 | MAPK3 |
| XRN1 | MAPK7 |
| XRN2 | MAPKAP1 |
| YARS | MAPRE3 |
| YME1L1 | MARVELD1 |
| YRDC | MAST3 |
| YTHDC1 | MAT2B |
| YTHDF1 | MAVS |
| YWHAE | MAZ |
| ZBP1 | MBD3 |
| ZBTB3 | MBD4 |
| ZBTB37 | MBP |
| ZBTB46 | MBTPS1 |
| ZC3H11A | MCAT |
| ZC3H12A | MCM2 |
| Zc3h15 | MCM3 |
| ZC3H7A | MCM3AP |
| ZC3HC1 | MCOLN1 |
| ZDHHC5 | MCTS1 |
| ZFAND3 | MDH1 |
| ZFP36 | MDH2 |
| ZFX | MDM1 |
| ZNF131 | MDP1 |
| ZNF143 | MED17 |
| ZNF319 | MED20 |
| ZNF598 | MED22 |
| ZNFX1 | MED24 |
| ZRSR2 | MED25 |
| ZUFSP | MED30 |
| ZYX | MEF2D |
| | MEPCE |
| | METAP2 |
| | METRN |

| | |
|---|---|
| | METRNL |
| | METTL13 |
| | METTL17 |
| | METTL21A |
| | METTL7A |
| | Mettl7a2 |
| | Mettl7a3 |
| | METTL8 |
| | METTL9 |
| | Mff |
| | MFN2 |
| | MFNG |
| | MFSD3 |
| | MFSD5 |
| | MFSD6 |
| | MIA3 |
| | MIB2 |
| | MICAL1 |
| | MICAL2 |
| | MICU1 |
| | MID1IP1 |
| | MIDN |
| | MIF |
| | MIIP |
| | MINOS1 |
| | MIOS |
| | MIS18A |
| | MIS18BP1 |
| | MKL1 |
| | MKNK1 |
| | MKNK2 |
| | MKRN1 |
| | MKRN2 |

| | |
|---|---|
| | MLEC |
| | MLF2 |
| | MLH1 |
| | MLH3 |
| | MLLT10 |
| | MLLT3 |
| | MLST8 |
| | MLXIP |
| | MLYCD |
| | MMAB |
| | Mmgt2 |
| | MMP19 |
| | MMS19 |
| | MMS22L |
| | MNT |
| | MOB2 |
| | MOCOS |
| | MOSPD1 |
| | MPC2 |
| | MPDU1 |
| | MPHOSPH8 |
| | MPHOSPH9 |
| | MPND |
| | MPP6 |
| | MPV17 |
| | MRFAP1 |
| | MRI1 |
| | MRM1 |
| | MROH1 |
| | MRPL1 |
| | MRPL18 |
| | MRPL2 |
| | MRPL20 |

| | |
|---|---|
| | MRPL28 |
| | MRPL34 |
| | MRPL36 |
| | MRPL40 |
| | MRPL41 |
| | MRPL42 |
| | MRPL43 |
| | MRPL44 |
| | MRPL49 |
| | MRPL51 |
| | MRPL57 |
| | MRPS12 |
| | MRPS15 |
| | MRPS16 |
| | MRPS21 |
| | MRPS24 |
| | MRPS25 |
| | MRPS26 |
| | MRPS33 |
| | MRPS34 |
| | MRPS35 |
| | MRPS36 |
| | MRRF |
| | MSH2 |
| | MSH6 |
| | MSL1 |
| | MSL3 |
| | MSRB1 |
| | MSRB2 |
| | MSTO1 |
| | MTCH1 |
| | MTCH2 |
| | MTERF4 |

| | |
|---|---|
| | MTFMT |
| | MTFP1 |
| | MTFR1L |
| | MTG1 |
| | MTHFD1 |
| | MTSS1 |
| | MUM1 |
| | MUS81 |
| | MUT |
| | MVB12B |
| | MVD |
| | MVK |
| | MXI1 |
| | MYEOV2 |
| | MYL12A |
| | MYO18A |
| | MYO1E |
| | MYO7A |
| | MYO9A |
| | MYO9B |
| | MZT2A |
| | N4BP2L1 |
| | N4BP3 |
| | NAA35 |
| | NAA38 |
| | NAA60 |
| | NABP2 |
| | NAGA |
| | NAGLU |
| | Naip1 |
| | NANP |
| | NANS |
| | NAP1L1 |

| | |
|---|---|
| | NAPA |
| | NAPG |
| | NARF |
| | NARFL |
| | NASP |
| | NAT6 |
| | NAT8B |
| | NATD1 |
| | NBEAL2 |
| | NBR1 |
| | NCAPD2 |
| | NCAPH |
| | NCAPH2 |
| | NCBP1 |
| | NCBP2 |
| | NCBP2-AS2 |
| | NCF2 |
| | NCKAP5L |
| | NCKIPSD |
| | NCLN |
| | NCOA1 |
| | NCOR2 |
| | NDC1 |
| | NDRG3 |
| | NDST1 |
| | NDUFA1 |
| | NDUFA11 |
| | NDUFA13 |
| | NDUFA2 |
| | NDUFA3 |
| | NDUFA5 |
| | NDUFA6 |
| | NDUFA8 |

| | |
|---|---|
| | NDUFAF4 |
| | NDUFAF7 |
| | NDUFB10 |
| | NDUFB11 |
| | NDUFB3 |
| | NDUFB5 |
| | NDUFB7 |
| | NDUFB8 |
| | NDUFB9 |
| | NDUFC1 |
| | NDUFS1 |
| | NDUFS2 |
| | NDUFS3 |
| | NDUFS7 |
| | NDUFS8 |
| | NDUFV1 |
| | NDUFV2 |
| | NDUFV3 |
| | Neat1 |
| | NECAP1 |
| | NEK3 |
| | NEK6 |
| | NEK9 |
| | NELFA |
| | NELFB |
| | NELFCD |
| | NENF |
| | NEU1 |
| | NEURL4 |
| | NF2 |
| | NFAM1 |
| | NFATC3 |
| | NFIC |

| | |
|---|---|
| | NFS1 |
| | NFX1 |
| | NFYC |
| | NGFRAP1 |
| | NGRN |
| | NICN1 |
| | NINJ1 |
| | Nisch |
| | NIT1 |
| | NKAIN1 |
| | NLK |
| | NLN |
| | NME1 |
| | NME2 |
| | NME7 |
| | NOP10 |
| | NOSIP |
| | NPC1 |
| | NPEPL1 |
| | NPEPPS |
| | NPHP1 |
| | NPRL2 |
| | NR1D2 |
| | NR2C2AP |
| | NR2F6 |
| | NRDC |
| | NRDE2 |
| | NSA2 |
| | NSF |
| | NSMCE4A |
| | NSUN4 |
| | NT5C |
| | NT5C2 |

| | |
|---|---|
| | NT5DC3 |
| | NTHL1 |
| | NTPCR |
| | NUCB2 |
| | NUCKS1 |
| | NUDC |
| | NUDCD2 |
| | NUDT1 |
| | NUDT16 |
| | NUDT16L1 |
| | NUDT19 |
| | NUDT2 |
| | NUDT22 |
| | NUDT6 |
| | NUDT7 |
| | NUMA1 |
| | NUP107 |
| | NUP133 |
| | NUP155 |
| | NUP93 |
| | NUTF2 |
| | OARD1 |
| | OAZ1 |
| | ODF2 |
| | OGDH |
| | OGFOD2 |
| | OGFOD3 |
| | OGG1 |
| | OGT |
| | OIT3 |
| | OLA1 |
| | OMA1 |
| | OPHN1 |

| | |
|---|---|
| | ORAI1 |
| | ORAOV1 |
| | ORC5 |
| | ORMDL1 |
| | ORMDL3 |
| | OSBP |
| | OSBPL11 |
| | OSBPL2 |
| | OSBPL9 |
| | OSGEP |
| | OTULIN |
| | OXA1L |
| | OXCT1 |
| | OXNAD1 |
| | OXR1 |
| | P2RX4 |
| | P2RY1 |
| | P3H1 |
| | P4HB |
| | PA2G4 |
| | PACS2 |
| | PAFAH1B3 |
| | PAFAH2 |
| | PAG1 |
| | PAGR1 |
| | PAICS |
| | PAIP2 |
| | PAIP2B |
| | PALM |
| | PAN2 |
| | PANK1 |
| | PANX1 |
| | PAPSS1 |

### Supplementary Table S3

PARD6A
PARK7
PARN
PARP1
PARP16
PARP6
PARS2
PATZ1
Paxip1
PBX2
PBX3
PCBP4
PCCA
PCGF1
PCGF6
PCMTD1
PCMTD2
PCNXL3
PCNXL4
PCTP
PCYOX1
PCYT1A
PCYT2
PDCD4
PDCD6
PDCL
PDE6D
PDE7B
PDGFA
PDGFC
PDHA1
PDHB
PDIK1L
PDK1
PDK2
PDK3
PDXK
PEBP1
PELI2
PELO
PER1
PER2
PER3
PEX11A
PEX12
PEX26
PEX6
PFDN1
PFDN5
PFKFB3
PFKL
PFKM
PFN2
PGAM1
PGD
PGLS
PGM1
PGP
PGRMC2
PHAX
PHB2
PHC1
PHF1
PHF12
PHF2
PHF20
PHF21A
PHF23
PHF7
PHKA2
PHKG2
PHLDA3
PHOSPHO2
PHPT1
PIAS3
PIGA
PIGF
PIGH
PIGK
PIGQ
PIGS
PIGY
PIH1D1
PIK3R2
PIK3R4
PINK1
PIP4K2A
PIP4K2C
PIP5K1C
PITHD1
PITPNA
Pitpnc1
PITPNM1
PJA1
PJA2
PKD1
PKDCC
PKIB
PKIG
PKM
PKN1
PLA2G12A
PLA2G15
PLA2G6
PLCG2
PLD3
PLEKHA1
PLEKHG2
PLEKHH3
PLEKHJ1
PLEKHO1
PLGRKT
PLIN2
PLIN3
PLPP2
PLXDC1
PLXNB2
PLXNC1
PMM1
PMS2
PMVK
PNKD
PNN
PNPLA2
PNPLA6
PNPLA7
PNPLA8
PNPO
POGLUT1
POLD1
POLD4
POLDIP2

POLE2
POLG2
POLI
POLK
POLR1D
POLR2A
POLR2B
POLR2E
POLR2F
POLR2I
POLR2J
POLR3G
POLR3GL
POLR3H
POLR3K
POLRMT
POMK
POMT1
PON2
POP5
POR
POT1
POU6F1
PPA2
PPARG
PPCDC
PPCS
PPIB
PPIH
PPM1G
PPM1M
PPOX
PPP1CA
PPP1R21
PPP1R3F
PPP2R1A
PPP2R4
Ppp2r5c
PPP3CA
PPP3R1
PPP4R3B
PPP5C
PQBP1
PQLC1
PQLC2
PQLC3
PRADC1
PRCP
PRDX2
PRDX4
PREB
PRELID1
PREPL
PRICKLE4
PRIM1
PRKACA
PRKACB
PRKAG1
PRKAG2
PRKAR1B
PRKCB
PRKCH
PRKD2
PRKRA
PRKRIR
PRMT3
PRNP
PROKR1
PRPF39
PRPF6
PRPSAP1
PRR12
Prr13
PRUNE
PSAP
PSEN2
PSENEN
PSMB1
PSMC3
PSMD13
PSMD2
PSME4
PSMG1
PSMG2
PSRC1
PSTK
PSTPIP1
PTCD2
PTDSS1
PTDSS2
PTEN
PTER
PTGER4
PTGR1
PTGR2
PTOV1
PTP4A2
PTPN21
PTPN22
PTPRA
PTPRS
PTRH2
PTTG1IP
PURA
PURG
PXMP2
PXMP4
PYCR2
PYCRL
PYGB
PYGL
Qars
QDPR
QRSL1
QSER1
R3hcc1
R3HDM1
RAB11B
RAB11FIP5
RAB14
RAB24
RAB28
RAB29
RAB31
RAB3D
RAB40C
RAB4A
RAB7B
RABAC1
RABGAP1
RABGGTA
RABL6

RAD1
RAD50
RAF1
RALBP1
RANBP10
RANBP9
RANGRF
RAPGEF6
RARB
RARS2
RASA3
RASAL3
RASSF3
RASSF5
RASSF7
RB1
RB1CC1
RBBP4
RBBP7
RBBP9
RBL2
RBM10
RBM14
Rbx1
RCAN3
RCBTB1
RCC2
RDH14
RECQL
REEP3
REEP5
RERE
REV3L
REXO2
RFC1
RFC2
RFC4
RFNG
RFWD3
RFXAP
RGL2
RGS10
RGS18
RGS19
RGS2
RHBDD3
RHEBL1
Rhno1
RHOT2
RHOV
RIC8A
RILPL1
RILPL2
RIT1
RMI1
RMND1
RMND5A
RMND5B
RNASEL
RNF10
RNF103
Rnf113a1
RNF123
RNF126
RNF128
RNF130
RNF166
RNF181
RNF187
RNF220
RNF26
RNF44
RNF5
RNH1
RNPEPL1
ROGDI
ROMO1
ROPN1L
RP9
RPA1
RPA2
RPAP1
RPAP2
RPL13A
RPL14
RPL18
RPL27
RPL3
RPL35
RPL36AL
RPL38
RPL5
RPLP2
RPN1
RPP21
RPP25L
RPRD1A
RPRD1B
RPS10
RPS21
RPS25
RPS28
RPS29
RPS6KA1
RQCD1
RRAGC
RRAS
RRM1
RRP1B
RSRC1
RSRP1
RTN3
RTTN
RUFY1
RUSC1
RXRA
RXRB
S100A13
S100A4
S100A6
SAC3D1
SAE1
SAFB
SAFB2
SAMD8
SAMM50
SAP18
SAP30L
SASH3
SBK1
SCAF11
SCAMP1

SCAMP3
SCAMP4
SCAMP5
SCAND1
SCARB1
SCARB2
SCCPDH
SCLY
SCPEP1
SCRIB
SDC1
SDF2
SDF4
SDHA
SDHAF1
SDHAF4
SDHB
SDHC
SDR42E1
SEC11C
SEC14L1
SEC16A
SEC24D
SEC61A1
SEC61B
SEL1L3
SELENBP1
SELM
SELPLG
SENP3
SEPHS2
09-Sep
SERAC1
SERF1A
SERF1B
SERF2
SERGEF
SERHL2
SESN1
SET
SETD1A
SETDB1
SF3B2
SF3B4
SF3B5
SFT2D1
SFXN1
SFXN3
SGF29
SGK1
SGPP1
SGSH
SGSM3
SGTA
SH2B1
SH2D3C
SH3BGRL3
SH3GLB2
SHARPIN
SHFM1
SHKBP1
SHPRH
SIDT2
SIGMAR1
SIN3A
SIPA1
SIRPA
SIRT3
SIRT4
SIRT7
SIVA1
SKI
SKP2
SLAIN2
SLC12A9
SLC15A4
SLC16A3
SLC16A6
SLC19A2
SLC25A13
SLC25A14
SLC25A15
SLC25A19
SLC25A20
SLC25A26
SLC25A28
SLC25A35
SLC25A38
SLC25A39
SLC25A44
SLC25A45
SLC26A6
SLC27A1
SLC29A3
SLC2A8
SLC30A5
SLC30A9
SLC33A1
SLC35A1
Slc35a2
SLC35A5
SLC35B4
SLC35C1
SLC35C2
SLC35F6
SLC37A4
SLC38A10
SLC39A11
SLC39A6
SLC39A7
SLC41A1
SLC44A2
SLC46A1
SLC48A1
SLC50A1
SLC52A2
SLC6A8
SLC8B1
SLC9A3R2
SLF1
SLF2
SLMAP
SLX1A
SLX1B
SMARCA2
SMARCAL1
SMARCB1
SMARCC1
SMARCD1
SMARCD2
SMC1A
SMC2

### Supplementary Table S3

SMC3
SMDT1
Smim11
SMIM12
SMIM14
SMIM19
SMPD1
SMPD2
SMPD4
SMYD2
SNAP47
SNAPC2
SNAPIN
SNRK
SNRNP25
SNRPD2
Snrpg
SNTA1
SNTB2
SNX12
SNX14
SNX15
SNX19
SNX21
SNX3
SNX30
SNX5
SOAT1
SORT1
SOS2
SPAST
SPC25
SPCS3

SPDL1
SPECC1
SPEG
SPEN
Spg20
SPG21
SPHK2
SPICE1
SPNS1
SPPL2B
SPR
SPRYD3
SPSB2
SPSB4
SPTSSA
SRD5A3
SREK1
SRP14
SRP9
SRPK2
SRR
SRSF9
SS18L2
SSBP3
SSBP4
SSNA1
SSRP1
SSU72
ST6GAL1
ST6GALNAC6
STAC2
STAM
STARD8

STEAP3
STIM1
STK11
STK11IP
STK38
STRADA
STT3A
STT3B
STUB1
STX17
STX5
STX7
STX8
STXBP2
SUCLA2
SUDS3
SUGP2
SUMF1
SUMO3
SUOX
SURF1
SVIL
SWSAP1
SYNE1
SYNGR1
SYNGR2
SYNRG
SYPL1
SYS1
SYT11
TAB1
TACC2
TADA1

TADA2A
TADA2B
TAF10
TAF13
TAF6
TALDO1
TAMM41
TANGO2
TARBP2
TARS2
TAX1BP3
TAZ
TBC1D14
TBC1D15
TBC1D17
TBC1D20
TBC1D22A
TBC1D24
TBC1D8
TBCD
TBCEL
TBL1XR1
TBL2
TCEAL8
Tceb2
TCEB3
TCF3
TCTA
TCTN3
TDP2
TEC
TECPR1
TEF

TELO2
TEN1
TEP1
TERF2
TERF2IP
TESK1
TEX2
TEX261
TEX264
TFCP2
TFEB
TGFBR2
THAP11
THAP7
THNSL1
THOP1
THTPA
THYN1
TIA1
TIAL1
TIMM10B
TIMM13
TIMM8B
TIMP2
TINF2
TIPIN
TK2
TKT
TLE1
TLN1
TM2D2
TM7SF3
TM9SF1

TM9SF2
TMC4
TMC6
TMCO1
TMED10
TMED3
TMED4
TMEM101
TMEM106C
TMEM109
TMEM129
TMEM134
TMEM141
TMEM147
TMEM14C
TMEM150A
TMEM158
TMEM160
TMEM161A
TMEM165
TMEM175
TMEM184C
TMEM186
TMEM189
TMEM19
TMEM199
TMEM203
TMEM205
TMEM206
TMEM214
TMEM216
TMEM218
Tmem223

TMEM230
TMEM242
TMEM259
TMEM261
TMEM33
TMEM38B
TMEM41A
TMEM41B
TMEM42
TMEM43
TMEM51
TMEM55A
TMEM60
TMEM71
TMEM80
TMEM82
TMEM8A
TMEM97
TMLHE
TMPPE
TMUB1
TMUB2
TMX2
TNFRSF12A
TNFRSF21
TNKS2
TNRC6A
TNRC6B
TNRC6C
TNS1
TOB1
TOB2
TOM1

TOMM40L
Tomm5
TOP2B
TOP3B
TOPBP1
Toporsos
TOR1A
TOR2A
TOR4A
TP53BP1
TP53INP1
TP53INP2
TPCN1
TPCN2
TPD52
TPD52L2
TPGS1
TPGS2
TPI1
TPMT
TPP1
TPRA1
TPRG1L
TPRKB
TPRN
TPST2
TRADD
TRAF7
TRAM1
TRAPPC1
TRAPPC2L
TRIAP1
TRIM11

TRIM16
TRIM2
TRIM24
TRIM27
TRIM28
TRIM32
TRIM37
TRIM39
TRIM41
TRIM44
TRIM62
TRIM8
TRIP12
TRMT10B
TRMT112
TRMT2A
TRRAP
TSC1
TSC22D3
TSC22D4
TSEN15
TSN
TSNAX
TSPAN14
TSPAN4
TSPAN5
TSSC4
TST
TTC13
TTC14
TTC3
TTC30A
TTC30B

TTC33
TTC4
TTC5
TTI1
TTLL1
TTYH2
TUBA4A
TUBD1
TUBG1
TUBGCP4
TUBGCP5
TXN
TXN2
TXNDC11
TXNDC17
TXNDC5
TXNRD2
TXNRD3
TYROBP
TYSND1
UAP1L1
Uba52
UBAC1
UBAC2
UBALD1
UBE2I
UBE2L3
UBE2O
UBE2R2
UBE2T
UBE3A
UBE3B
UBE3C

UBE4B
UBFD1
UBN1
UBQLN2
UBR3
UBR7
UBTD1
UBXN1
UBXN6
UCHL5
UCKL1
UFD1L
UFSP2
UGDH
UGP2
UHRF1BP1L
UIMC1
ULK1
ULK2
UMPS
UNC119
UQCC2
UQCC3
UQCR10
UQCR11
UQCRB
URI1
UROD
USF2
USP20
USP21
USP34
USP4

USP45
USP48
USP5
UVRAG
VAC14
VAMP4
VAMP8
VAPB
VAT1
VCP
VDAC1
VEGFA
VEGFB
VIPAS39
VKORC1
VMP1
VPS11
VPS18
VPS26B
VPS28
VPS39
VPS41
VPS45
VPS53
VPS9D1
VRK3
VTI1A
VTI1B
WAS
WASH1
WBP1
WBP1L
WBP2

WBP5
WDR13
WDR26
WDR45
WDR6
WDR77
WDTC1
WEE1
WIPF1
WIPI1
WIPI2
WNK1
WRAP53
WRNIP1
WSB2
WWC2
WWOX
WWP2
XPA
XPC
XPO1
XPO7
XPR1
XRCC1
XRCC2
XRCC6
XYLB
YEATS4
YIPF2
YLPM1
YPEL3
YPEL5
YWHAB

YWHAH
ZBED3
ZBTB1
Zbtb11os1
ZBTB14
ZBTB2
ZBTB20
ZBTB25
ZBTB44
ZBTB45
ZBTB48
ZC3H14
ZC3H8
ZCCHC17
ZDHHC1
ZDHHC12
ZDHHC14
ZDHHC16
ZDHHC4
ZDHHC7
ZDHHC8
ZDHHC9
ZFAND2A
ZFAND5
ZFAND6
Zfp101
ZFP36L2
Zfp472
Zfp52
Zfp53
Zfp61
Zfp740
Zfp810

Zfp87
Zfp871
ZFP91
Zfp932
Zfp942
Zfp944
Zfp954
Zfp955a
Zfp955b
Zfp958
ZFYVE19
ZFYVE21
ZFYVE26
ZKSCAN1
ZKSCAN3
ZKSCAN5
ZMAT3
ZMYM1
ZMYND8
ZNF106
ZNF148
ZNF207
ZNF219
ZNF235
ZNF260
ZNF280C
ZNF292
ZNF316
ZNF318
ZNF329
ZNF362
ZNF385A
ZNF426

ZNF451
ZNF467
ZNF503
ZNF524
ZNF566
ZNF688
ZNF703
ZNF704
ZNF706
ZNF707
ZNF746
ZNF761
ZNF764
ZNHIT2
ZNRD1
Znrd1as
ZPR1
Zrsr1
ZSCAN12
ZSCAN26
ZSWIM1
ZW10
ZWINT
ZXDC
ZYG11B
